# KRAS is vulnerable to reversible switch-II pocket engagement in cells

**DOI:** 10.1101/2021.10.15.464544

**Authors:** James D. Vasta, D. Matthew Peacock, Qinheng Zheng, Joel A. Walker, Ziyang Zhang, Chad A. Zimprich, Morgan R. Thomas, Michael T. Beck, Brock F. Binkowski, Cesear R. Corona, Matthew B. Robers, Kevan M. Shokat

**Author notes:** These authors contributed equally (James D. Vasta, D. Matthew Peacock, Qinheng Zheng).

## Abstract

Current small molecule inhibitors of KRAS(G12C) bind irreversibly in the switch-II pocket, exploiting the strong nucleophilicity of the acquired cysteine as well as the preponderance of the GDP-bound form of this mutant. Nevertheless, many oncogenic KRAS mutants lack these two features, and it remains unknown whether targeting the switch-II pocket is a practical therapeutic approach for KRAS mutants beyond G12C. Here we use NMR spectroscopy and a novel cellular KRAS engagement assay to address this question by examining a collection of SII-P ligands from the literature and from our own laboratory. We show that the switch-II pockets of many GTP hydrolysis-deficient KRAS hotspot (G12, G13, Q61) mutants are accessible using non-covalent ligands, and that this accessibility is not necessarily coupled to the GDP state of KRAS. The results we describe here emphasize the switch-II pocket as a privileged drug binding site on KRAS and unveil new therapeutic opportunities in RAS-driven cancer.

## Introduction

The KRAS proto-oncogene is the most frequently mutated oncogene in cancer^1^. Glycine-12 mutations are the most common, with KRAS(G12D) representing the most common substitution in pancreatic ductal adenocarcinoma (PDAC) and colorectal tumors^1^. KRAS had long been considered “undruggable” until the identification of covalent drugs targeting KRAS(G12C).^2,3^ The drug sotorasib (AMG510) was recently approved for treatment of patients with the KRAS(G12C) mutation with six additional drugs targeting this same mutant currently under clinical investigation^4–6^. Several features unique to KRAS(G12C) enabled this allele to be the first KRAS mutant to be drugged. The somatic mutation of glycine 12 to cysteine provided the opportunity to exploit covalent drug discovery methods which are not applicable to the other common KRAS alleles (e.g., G12D/G12V/Q61H). Sotorasib and other known irreversible KRAS(G12C) drugs bind to the switch-II pocket (SII-P) and only engage the inactive GDP-state of KRAS(G12C)^2,5–10^. A rare example of a molecule reported to target the active GTP-state was recently disclosed which relies on a “molecular glue” mechanism involving the recruitment of cyclophilin not widely applicable to other KRAS(G12C) inhibitors.^11^ Fortunately, KRAS(G12C) is also unique among the KRAS oncogenes in maintaining wild-type like intrinsic GTPase activity – thereby allowing for successful GDP-state targeting for this allele^12^. In order to effectively inhibit other oncogenic KRAS alleles that do not adequately sample the GDP-state in cells, drugs which bind reversibly to the GTP-state will likely be required.

Studies using engineered proteins and cyclic peptides to probe the SII-P of KRAS have revealed the dynamic nature of this pocket and support the possibility that KRAS-GTP may adopt conformations favorable to SII-P engagement^13–17^. However, proteins and most cyclic peptides are impermeable to cell membranes, making them difficult to use as drug leads. The recent flurry of drug discovery aimed at targeting KRAS reported in the literature and patent filings might provide suitable small molecule leads for reversible KRAS inhibition. However, the nucleotide state requirements of these molecules are unknown. Furthermore, robust methods to directly measure in-cell non-covalent engagement of KRAS that do not rely on downstream signals or phenotypic effects are surrogates are unknown.

In this study, we investigated the reversible binding of KRAS small molecule inhibitors to determine which hotspot mutants are vulnerable to SII-P engagement in cells. We used HSQC NMR spectroscopy to directly observe reversible binding to the SII-P of the KRAS *in vitro* and determine the nucleotide state dependency of binding. We developed a bifunctional cell-permeable fluorescent probe from the SI/II pocket inhibitor BI-2852, and this probe was utilized in a competitive Bioluminescence Resonance Energy Transfer (BRET) format^18,19^ to quantify SII-P engagement to multimeric RAS complexes in live cells. These studies represent the first observation and quantification of direct target engagement of non-G12C oncogenic KRAS mutants in cells by reversible binders. Our results expose a wide scope of vulnerability to SII-P engagement across hotspot mutants and should help guide the development of future inhibitors and therapeutics.

## Results

### Reversible binding to the KRAS SII-P is observed *in vitro* by NMR spectroscopy

The well-established and clinically validated inhibitors of KRAS(G12C) such as ARS-1620, AMG510, and MRTX849 (Fig 1A) rely on a covalent reaction with the nucleophilic cysteine 12 in the GDP-state. Although these molecules bind the same pocket and are similar chemotypes, MRTX849 (and the closely related MRTX1257) possess unique structural elements proposed to increase the reversible component of their binding, resulting in a measurable *K*_I_ of 3.7 μM for the reaction of MRTX849 with KRAS(G12C).^7^. We sought to determine whether these molecules also bind RAS proteins lacking the G12C mutation and whether their reversible affinity is specific to the inactive GDP-state.

**Figure 1.**
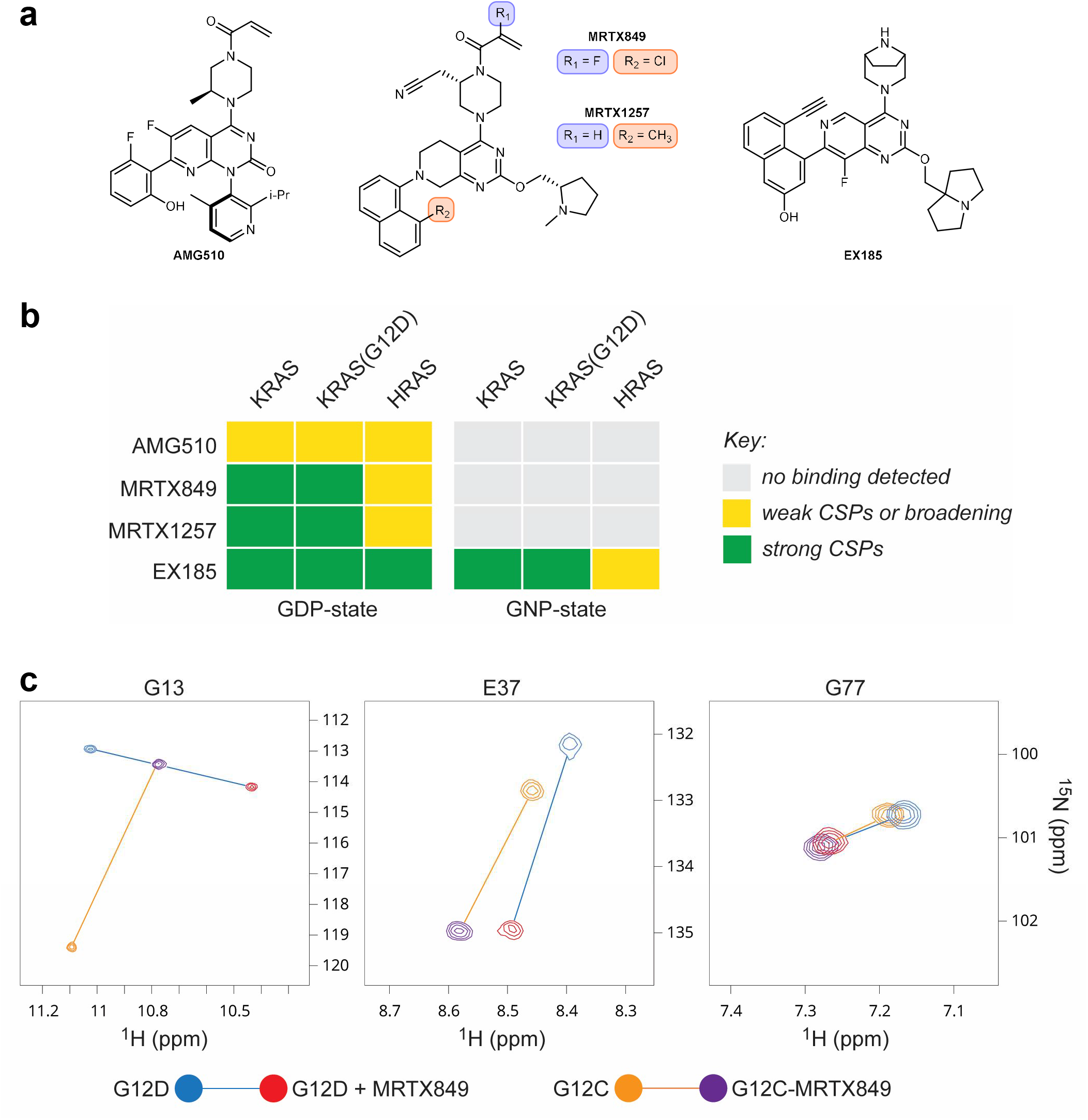
*In vitro* non-covalent binding to the KRAS SIIP determined by NMR spectroscopy. **a.** Chemical structures of AMG510, MRTX849, MRTX1257, and EX185. **b.** Summary of the effects of SIIP-binders on ^1^H-^15^N HSQC NMR spectra of RAS proteins. **c.** Examples of CSPs of GDP-loaded KRAS in the presence of MRTX849 and comparison of irreversible binding to KRAS(G12C) and reversible binding to KRAS(G12D). Spectra recorded at pH 7.4 and 298 K with 100 μM U-^15^N protein and 200 μM ligand.

Cell-free analysis of binding to RAS proteins was performed via protein-observed NMR spectroscopy (Fig 1). We expressed uniformly ^15^N-labelled KRAS 1-169, KRAS(G12C) 1-169, KRAS(G12D) 1-169, and HRAS 1-166 proteins and acquired a series of ^1^H-^15^N HSQC NMR spectra. The addition of either MRTX849 or MRTX1257 (200 μM) to the GDP-loaded state of either KRAS or KRAS(G12D) protein (100 μM) resulted in the formation of a new complex with strong chemical shift perturbations (CSPs) from the peaks of the unbound protein (Fig 1B-C). The same CSPs were observed under more dilute conditions (50 μM protein and 100 μM ligand) for all four of these protein-ligand combinations, and no exchange between the bound and unbound protein could be detected in a sample containing KRAS-GDP protein sub-stoichiometrically ligated by MRTX849. Although the lack of chemical exchange poses a challenge to assigning most peaks of the protein-ligand complexes to their respective residues, when these spectra were compared to those of KRAS(G12C) and the covalent KRAS(G12C)-MRTX849 complex, similarities in perturbations of some well-resolved peaks are clear (e.g., G77) and support a similar binding mode between the non-covalent (WT and G12D) and covalent (G12C) protein-ligand complexes (Fig 1C, Fig S1). These data indicate that MRTX849 and MRTX1257 tightly and non-covalently bind GDP-loaded KRAS proteins lacking the G12C mutation with *K*_D_ and *k*_off_ values too small to be quantified by HSQC NMR spectroscopy.

In contrast, no significant effects were observed on the spectra of KRAS or KRAS(G12D) proteins containing the non-hydrolyzable GTP analog GPPNHP (GNP) with 200 μM of either MRTX849 or MRTX1257 (Fig 1B). Furthermore, only weak CSPs were observed from the addition of either molecule to GDP-loaded HRAS 1-166 under the same conditions. Instead, concentration-dependent broadening and/or decreases in volume were observed for many peaks corresponding to residues in the SII-P, suggesting only weak occupancy of the HRAS SII-P even at the highest concentration tested (100 μM protein and 300 μM ligand). The results of these HSQC experiments show that MRTX849 and MRTX1257 bind KRAS proteins with high selectivity for the inactive GDP-loaded state and for the K-isoform over HRAS.

A series of similar ^1^H-^15^N HSQC NMR experiments provided some evidence for weak binding of AMG510 (Fig 1A) to the SII-P of GDP-loaded KRAS and HRAS proteins (Fig 1B). Peaks corresponding to residues in the SII-P broadened and exhibited weak (generally less than line-widths) CSPs in the presence of 200 μM of AMG510. These experiments suggest that the reversible affinity of AMG510 to RAS proteins is likely too weak to be relevant to in-cell experiments conducted at lower concentrations, and that AMG510 must rely on the irreversible reaction at the mutant cysteine 12 for its inhibitory activity, which is consistent with previously published data^5^.

Recently, compounds reported to target KRAS(G12D) were disclosed in patent applications by multiple groups^20–23^. We selected and synthesized an example from these patent filings^23^ (EX185) with structural features similar to MRTX849/1257 (Fig 1A). We found that EX185 bound GDP-loaded KRAS and KRAS(G12D) by HSQC NMR spectroscopy, similarly to MRTX849/1257 (Fig 1B). However, in stark contrast, EX185 also bound the active GNP-state of these proteins, resulting in new complexes with strong CSPs from the unbound proteins. Identical CSPs were observed in samples containing either 50 μM protein and 100 μM ligand or 100 μM protein and 200 μM ligand. Since EX185 tightly bound both nucleotide states of KRAS and KRAS(G12D) by HSQC NMR spectroscopy, we determined its nucleotide state preference by adding a sub-stoichiometric amount of EX185 (50 μM) to a sample containing a 1:1 mixture of GDP- and GNP-loaded KRAS protein (100 μM each) (Fig S2). This mixture resulted in exclusive formation of the GDP-KRAS-EX185 complex, and the same experiment with KRAS(G12D) yielded the same result. These results suggest that the relative affinity of EX185 to the GDP-state over the GNP-state of KRAS proteins is greater than the noise limit of the spectra (>10 for most peaks).

These cell-free NMR experiments show that MRTX849 and MRTX1257 engage KRAS proteins even in the absence of a nucleophilic mutant cysteine 12. However, this engagement is selective for the inactive GDP-loaded state of the protein. The more recently disclosed EX185, in contrast, engages both nucleotide states – albeit with preference for the inactive GDP-loaded protein – and might present an opportunity to inhibit even constitutively active (GTP-loaded) KRAS hotspot mutants. However, these NMR experiments require high concentrations of proteins and do not quantify the potency of these tightly-binding compounds. Furthermore, *in vitro* binding assays may not be representative of the in-cell vulnerability of a regulated, effector-bound, and membrane-localized protein such as KRAS.

### KRAS is broadly vulnerable to reversible SII-P target engagement in cells

With our NMR results supporting the potential of KRAS and its hotspot mutants to be vulnerable to non-covalent SII-P occupancy, we asked whether these SII-P ligands engage KRAS in cells. We first assessed the anti-proliferative effects of MRTX849 in a number of G12C and non-G12C KRAS-mutant and KRAS-wildtype cell lines (Fig S3A). Although MRTX849 inhibited the growth of SW1990 [KRAS(G12D)] and HCT-116 [KRAS(G13D)] at micromolar concentrations, it also had the same effect on HEK-293 (KRAS wildtype, RAS-independent) (Lim 2021) and A375 (BRAF V600E, RAS-independent), suggesting the antiproliferative effects may originate from RAS-independent toxicity (Fig S3A). We also measured the ability of MRTX849 to inhibit ERK phosphorylation in a similar panel of cell lines (Fig S3B). We corroborated the strong potency of MRTX849 in KRAS(G12C) driven cell lineages. However, in non-G12C driven cell lineages, the non-specific cytotoxic effects were observed over the same concentration range as the inhibitory effects on ERK phosphorylation (Fig S3B), thus preventing a clear confirmation of cellular target engagement.

The interference from off-target toxic effects in these assays precluded the analysis of target engagement and prompted us to develop new approaches to determine ligand-RAS interaction in cells. To more directly query biophysical engagement of KRAS and HRAS to small molecule target engagement in cells, a BRET reporter system was developed. We synthesized a pan-RAS BRET probe by conjugating a fluorophore to a derivative of the reversible SI/II-P inhibitor BI-2852 (Fig 2A). Recognizing the multimeric and membrane-localized nature of RAS ^24–27^, we sought to generate a BRET signal conditionally within membrane-associated RAS complexes^28^. We configured a luminescent complementation-based system that was dependent upon RAS lipidation as the BRET donor (Fig 2B-C, Fig S4). When cells expressing the BRET donor complexes were treated with the SI/II BRET probe, we observed a strong BRET signal that was readily competed by unmodified BI-2852 in cells (Fig 2D, Fig S4). To evaluate the sensitivity of the SI/II-P BRET probe to allosteric target engagement within the SII-P, live HEK-293 cells expressing NanoBiT-KRAS(G12C) were challenged with SII-P ligands AMG510 or ARS-1620 in the presence of the SI/II-P BRET probe. Time- and dose-dependent competition was observed between AMG510 or ARS-1620 and the BRET probe (Fig 2E, Fig S5A). At a 2 h timepoint, BRET results with both AMG510 and ARS-1620 closely matched the potency of endogenous target engagement and phospho-ERK inhibition at identical timepoints in a number of G12C-driven lineages (Mia PaCa-2, NCI-H358), corroborating the accuracy of the BRET method as a proxy for engagement in an endogenous cellular setting (Fig 2E, Fig S5B). AMG510 demonstrated exquisite engagement selectivity for KRAS(G12C) compared to KRAS WT, other KRAS hotspot mutants, and HRAS WT (Fig S5C), consistent with previous reports for functional selectivity between KRAS(G12C) and non-G12C driven cancer cell lines. Additional SII-P inhibitors were evaluated at KRAS(G12C) complexes, including MRTX849 and MRTX1257 (Fig S5D). Each produced BRET target engagement results that agreed closely with published cellular potency at KRAS(G12C) lineages^7^. MRTX849/1257 were the most potent KRAS(G12C) inhibitors in the analysis, in close agreement with previous studies ^7^. Together the results for engagement of KRAS(G12C) with SII-P ligands support the potential of the BRET target engagement system to report on KRAS in its endogenous cellular setting, and that this system can be used to accurately query engagement across oncogenic KRAS mutants in live cells.

**Figure 2.**
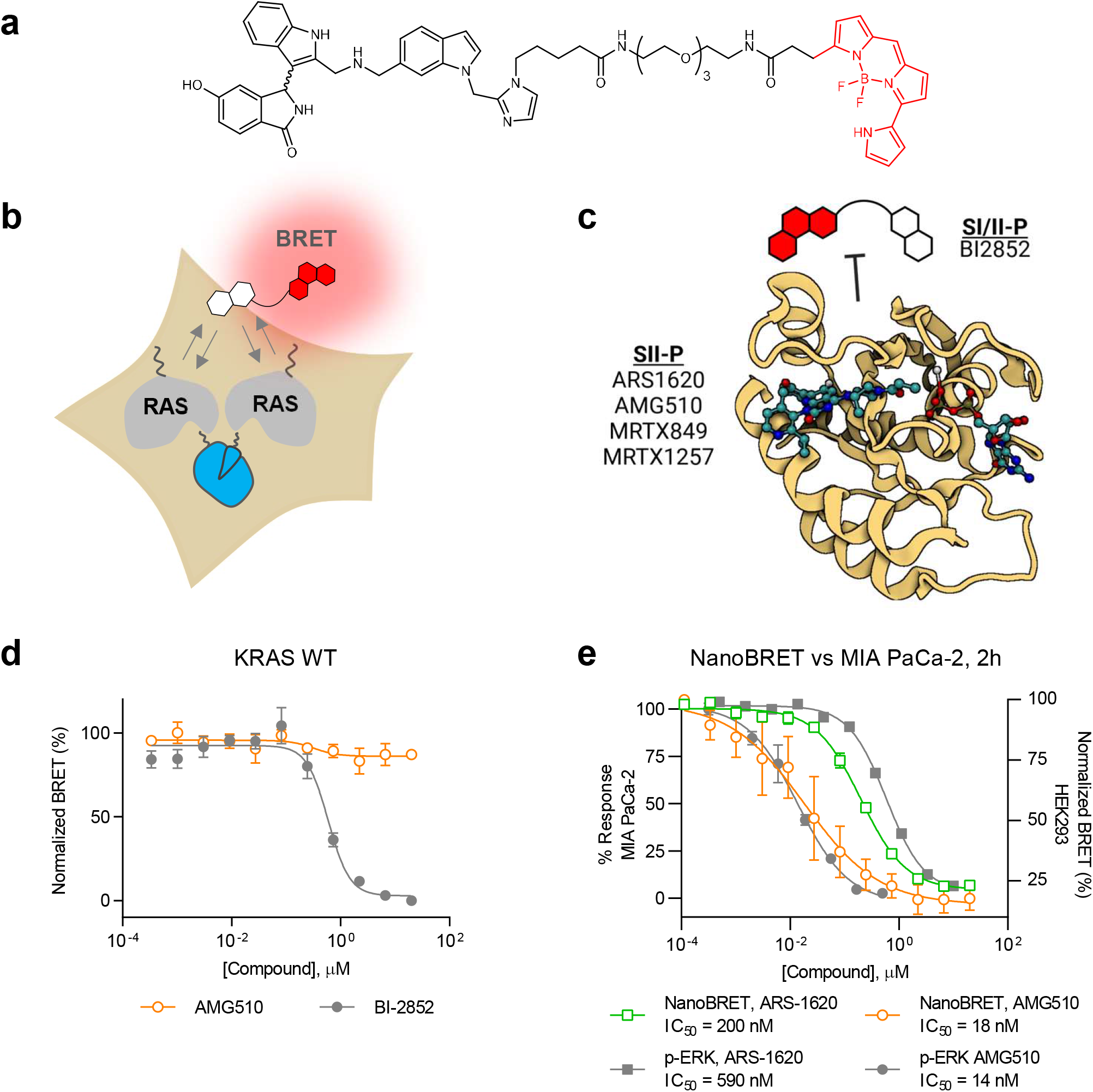
Target engagement assay for RAS. **a.** Chemical structure of RAS Switch I/II pocket BRET probe. **b.** Illustration of the RAS Cellular Target Engagement assay. A luminescent complex is formed between RAS multimers in live cells. **c.** In a competitive displacement format, a switch I/II BRET probe is used to quantify binding of unmodified ligands to RAS in live cells. Image created with BioRender.com from PDB 6OIM (AMG510-KRAS(G12C)). **d.** Live cell target engagement is observed at wildtype KRAS with BI-2852, but not with AMG-510. BRET data are normalized based on complete occupancy (0% BRET) and no occupancy (100% BRET) of the BRET probe. **e.** Comparison of BRET target engagement data at KRAS (G12C) versus phospho-ERK from MIA PaCa cells as reported by Canon et al. 2019. Data are means of 4 independent experiments ± S.E.M. (n=4).

This BRET target engagement system enabled us to directly assess the engagement of wild type KRAS and numerous critical hotspot mutants by SII-P ligands. For example, following 2 h incubation, engagement of both MRTX849 and MRTX1257 was observed for wildtype KRAS complexes in the sub-micromolar range (IC_50_ < 600 nM) (Table 1, Fig 3B, Fig S6A). When KRAS hotspot mutants were evaluated, a wide spectrum of engagement was observed (Table 1, Fig 3B-C, Fig S6B−G). Though no engagement of MRTX849/1257 was observed for KRAS Q61R, and only weak engagement was observed for KRAS(G12V) and KRAS(G12D), modest engagement was observed for the remaining KRAS hotspot mutants in the single-digit micromolar range (IC_50_ ranging from 1-5 μM). Among the KRAS hotspot mutants excluding KRAS(G12C), the most potent engagement was observed for G13D, Q61H, and Q61L (Fig S6D−F).

**Table 1.** Engagement potency values (IC_50_, nM)^a^ for SII-P binders across RAS variants in HEK-293 cells using the BRET assay. ^a^IC_50_ values are the mean ± S.E. of at least three independent experiments, collected after a 2 hr incubation. ^b^Not determined. ^c^IC_50_ values are from a single biological replicate with 4 technical replicates.

| Compound | KRAS WT | KRAS(G12C) | KRAS(G12D) | KRAS(G12V) | KRAS(G13D) |
| --- | --- | --- | --- | --- | --- |
| ARS-1620 | N/D<sup>b</sup> | 210 ± 20 | N/D<sup>b</sup> | N/D<sup>b</sup> | N/D<sup>b</sup> |
| AMG510 | No Binding | 30 ± 10 | No Binding | No Binding | No Binding |
| MRTX849 | 550 ± 80 | 3.3 ± 0.9 | 3900 ± 300 | >60,000 | 1800 ± 300 |
| MRTX1257 | 600 ± 20 | 1.9 ± 0.3 | 5000 ± 1000 | >60,000 | 1700 ± 300 |
| CPD1 | 730 ± 60 | > 10,000 | > 30,000 | > 30,000 | 1130 ± 60 |
| CPD2 | 580 ± 80 | > 10,000 | > 30,000 | > 30,000 | 1180 ± 14 |
| CPD3 | > 10,000 | > 100,000 | > 100,000 | > 100,000 | > 30,000 |
| CPD4 | 1600 ± 300 | > 10,000 | > 10,000 | > 30,000 | 3400 ± 400 |
| CPD5 | 3000 ± 1000 | > 30000 | > 30,000 | >30,000 | 5000 ± 1000 |
| CPD6 | 410 ± 90 | 3000 ± 800 | > 10,000 | >30,000 | 850 ± 50 |
| EX185 | 110 ± 50 | 290<sup>c</sup> | 90 ± 20 | 700 ± 200 | 240<sup>c</sup> |

**Table 1. Engagement potency values (IC<sub>50</sub>, nM)<sup>a</sup> for SH-P binders across RAS variants in HEK-293 cells using the BRET assay.**
| Compound | KRAS(Q61H) | KRAS(Q61L) | KRAS(Q61R) | HRAS WT |
| --- | --- | --- | --- | --- |
| ARS-1620 | N/D<sup>b</sup> | N/D<sup>b</sup> | N/D<sup>b</sup> | N/D<sup>b</sup> |
| AMG510 | No Binding | No Binding | No Binding | No Binding |
| MRTX849 | 1200 ± 200 | 1900 ± 200 | > 60,000 | > 60,000 |
| MRTX1257 | 700 ± 100 | 2200 ± 200 | > 60,000 | > 60,000 |
| CPD1 | 1300 ± 100 | 2000 ± 300 | > 100,000 | > 100,000 |
| CPD2 | 1300 ± 200 | 3000 ± 300 | > 100,000 | > 100,000 |
| CPD3 | > 30,000 | > 30,000 | > 100,000 | > 100,000 |
| CPD4 | 3600 ± 600 | 7400 ± 700 | > 30,000 | > 30,000 |
| CPD5 | 6000 ± 2000 | > 10,000 | > 100,000 | > 100,000 |
| CPD6 | 900 ± 40 | 1000 ± 100 | > 30,000 | > 100,000 |
| EX185 | 130<sup>c</sup> | 290<sup>c</sup> | 1400<sup>c</sup> | > 10,000 |

**Figure 3.**
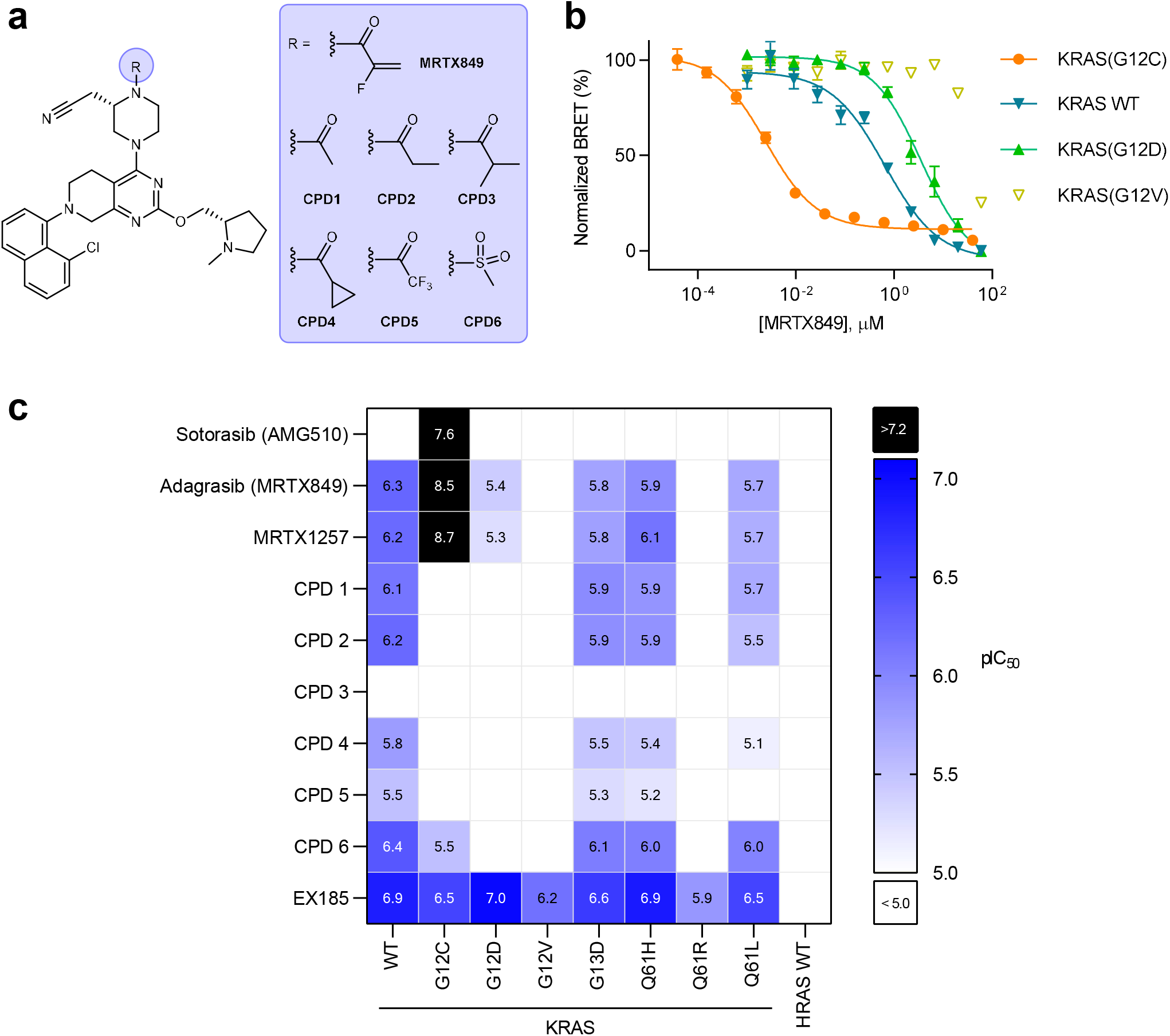
Profiling in-cell target engagement of SII-P binding molecules. **a.** Chemical structures of MRTX849, non-covalent derivatives CPD1−6, and EX185. **b.** BRET target engagement profiles for MRTX849 at KRAS WT and G12 hotspot mutants. BRET data are normalized based on complete occupancy (0% BRET) and no occupancy (100% BRET) of the BRET probe. Data are representative of three independent experiments, where each data point is the mean of 4 technical replicates ± S.D. (n=3). **c.** Summary of BRET target engagement across KRAS hotspot mutants and HRAS. pIC_50_ values were calculated as −Log_10_(IC_50_ (M)). Combinations that exhibited incomplete engagement at the highest concentration tested (10^−4^ M) were grouped as pIC_50_ < 5.0 (white cells).

Wild type HRAS as well as two oncogenic HRAS mutants (G12C and G12V) were also evaluated for SII-P vulnerability using the BRET assay. No engagement was observed for wild type HRAS with AMG510, MRTX849, or MRTX1257 (Fig S5C, Fig S6H). Though HRAS(G12V) was also not vulnerable to SII-P engagement (Fig S6I), HRAS(G12C) showed vulnerability to both AMG510 and MRTX849 (Fig S6J). AMG510 demonstrated similar intracellular affinity towards HRAS(G12C) compared to KRAS(G12C), but MRTX849 demonstrated affinity for HRAS(G12C) that was 3 orders of magnitude weaker than that observed for KRAS(G12C), suggesting that the MRTX849 scaffold preferentially engages the K-isoform of RAS, which is consistent with our NMR spectroscopy results.

We next sought to accurately assess the contribution to SII-P engagement from non-covalent ligand-protein interactions. Due to the potential differences in the steric and electrostatic environments for the SII-P among the RAS variants, we synthesized derivatives of MRTX849 lacking the covalent acrylamide warhead and positioning groups with varied steric and electronic properties proximal to residue 12 (CPD1-6, Fig 3A), and we evaluated these compounds with the BRET target engagement assay (Fig 3C, Table 1, Fig S7A). For most RAS variants, the saturated amide and sulfonamide derivatives (CPD1−6) demonstrated comparable rank order vulnerability to those of MRTX849 and MRTX1257. Among non-G12C variants, wild type KRAS remained the most vulnerable of all RAS isoforms to reversible engagement, followed closely by hotspot KRAS mutants G13D and Q61H. KRAS(G12V), KRAS(Q61R), and wild type HRAS showed weak to no engagement across all saturated amides, similar to the results observed with MRTX849/1257. Engagement of KRAS(G12C) by most of the saturated amide derivatives was significantly impaired in the absence of the covalent mechanism, with the exception of the sulfonamide (CPD6), which demonstrated modest single digit micromolar affinity. KRAS(G12D), which showed weak engagement by MRTX849/1257, was less vulnerable to the saturated amides, suggesting that more significant chemical modifications will be necessary to effectively target this oncogene.

Within the saturated amide series, CPD1 and CPD2 containing acetamide and propionamide moieties, respectively, were generally well tolerated and showed engagement potencies similar to MRTX849 for most non-G12C-RAS isoforms. CPD6 presenting a sulfonamide was also well tolerated, in most cases demonstrating comparable engagement potency to the CPD1 and CPD2, except in the case of KRAS(G12C) where it was found to be moderately selective compared to other derivatives. CPD5 presenting an electron deficient trifluoroacetamide demonstrated right-shifted moderate to weak potency in all cases compared to CPD1 and CPD2, suggesting the importance of polar interactions with the amide carbonyl^29^. CPD3, the most sterically bulky analog presenting an isobutyramide, was poorly tolerated and demonstrated the weakest engagement potency among all of the amides across all RAS isoforms. Posing a ring constraint to the branched isopropyl group (i.e. the cyclopropyl carboxamide presented in CPD4) improved the potency compared to CPD3, but still demonstrated only moderate to weak potency in most cases. Overall, we observed weak but detectable cellular engagement of a spectrum of RAS mutants by these amide derivatives with IC_50_s in the micromolar range. These compounds elicited cytotoxic effects in a RAS-independent cell line at similar concentrations (Fig S7B); however, our BRET system permitted the direct measurement of SII-P engagement without the interference from off-target toxicity.

### EX185 engages KRAS hotspot mutants in cells and drives antiproliferation

Because our NMR results demonstrated the unique capability of EX185 to bind to both the GDP-state and the GTP-state of KRAS(G12D) in a cell-free system, we next evaluated this compound in a cellular setting using our BRET assay. Potent target engagement (Table 1, Fig 3C, Fig 4A, IC_50_ value of 90 nM) was observed for EX185 with KRAS(G12D), greatly surpassing engagement potency of the GDP state-selective MRTX849 derivatives. Though EX185 and MRTX849 have some similar structural features, it is unknown whether both molecules engage KRAS in a similar pose within the SII-P. To provide support for engagement of EX185 within the KRAS SII-P, we also assayed engagement of KRAS(Y96D), which contains a previously reported mutation conferring resistance to described SII-P inhibitors including MRTX849^11^(Fig S8A). KRAS(Y96D) engagement was not observed with MRTX849, CPD2, or EX185 by the BRET-based assay. This finding, in conjunction with the finding that all of the MRTX chemotypes in this study show weak to no binding to HRAS variants (in which residue 95 is a glutamine), is consistent with binding to the SII-P in a similar pose.

EX185 was also evaluated for inhibition of KRAS(G12D):effector interactions in cells using a NanoBiT protein-protein interaction assay. EX185 demonstrated time- and dose-dependent inhibition of the KRAS(G12D):CRAF(RBD) interaction (Fig S8B), providing support for functional disruption of MAPK signaling. Phospho-ERK and cell viability analysis in SW1990 cells confirmed that engagement with EX185 translated into inhibition of mitogenic signaling and an anti-proliferative effect in a G12D-driven lineage (Figure 4B-C), with anti-proliferative potency (70 nM) in close agreement with the BRET readout. Unlike the MRTX849 derivatives, nonspecific cytotoxicity did not confound the anti-proliferative results, as EX185 did not inhibit proliferation in a panel of control cell lines (Fig S8C). Taken together, these results along with the NMR findings indicate that GTP state compatibility may support the superior SII-P engagement for KRAS(G12D) in cells. We therefore attempted extend the utility of EX185 to additional KRAS hotspot alleles. EX185 engaged numerous KRAS Q61, G12, and G13 mutant alleles (Table 1, Fig 3C, Fig 4A, Fig S8D). Notably, EX185 engaged KRAS(G12V) in cells, which is the most GTP-biased G12 allele described. While the potency against individual mutants may require tailored chemical optimization, and engagement of wildtype KRAS may constrain the therapeutic window, our observation that a SII-P ligand can engage several GTP hydrolysis-deficient KRAS mutants signifies exciting opportunities to drug these KRAS mutants through this pocket.

**Figure 4.**
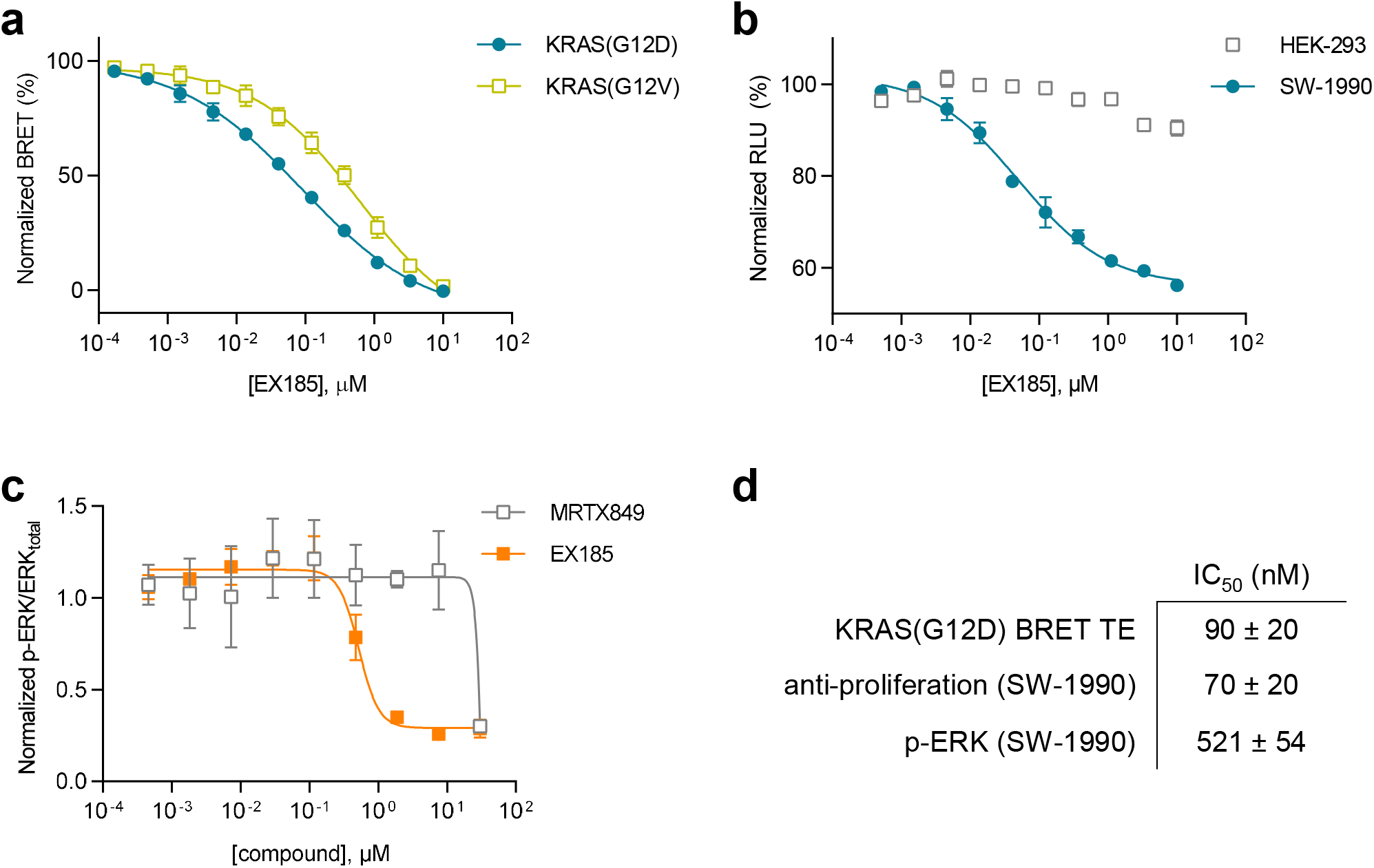
Characterization of engagement of KRAS with EX185. **a.** BRET target engagement profiles for EX185 at KRAS(G12D) and KRAS(G12V). BRET data are normalized based on complete occupancy (0% BRET) and no occupancy (100% BRET) of the BRET probe. Data are means of 3 independent experiments ± S.E.M. (n=3). **b.** EX185-driven antiproliferation (cell-titer glow) is observed in SW-1990 [KRAS(G12D)] but not in HEK-293 [KRAS-independent] cells. Data are the means ± S.E.M. of 3 independent experiments, each performed with at least 3 technical replicates (n=3). **c.** EX185 inhibits phospho-ERK. Data are means ± S.E.M. of 3 independent experiments (n = 3). **d.** Summary of IC_50_ values for EX185 target engagement (BRET) at KRAS(G12D) compared to anti-proliferation and inhibition of phospho-ERK in SW-1990 cells.

## Discussion

Here we report subfamily-wide engagement of KRAS hotspot mutants with the preclinical inhibitor MRTX849 and structurally related molecules. This is the first evidence of intracellular SII-P vulnerability across the prevalent oncogenic KRAS mutants including KRAS(G12D). To characterize target engagement across RAS species, we combined *in vitro* and intracellular biophysical approaches. NMR spectroscopy provided a defined system to observe reversible, non-covalent binding and to determine the impact of nucleotide status on KRAS vulnerability. However, cell-free methods are incapable of simulating the intracellular architecture where target engagement would naturally occur. To query engagement in cells, we developed a SI/II-P BRET probe that was competent to detect a variety of intracellular engagement mechanisms including ligands selective for either SI/II or SII pockets. The BRET method reported here conditionally measures engagement at membrane-localized RAS complexes in cells. Target engagement results with known SII-P covalent inhibitors matched both engagement and MAPK inhibition within endogenous G12C-driven lineages, supporting the accuracy of the engineered BRET method.

Although AMG510 and MRTX849 are related chemotypes, the unique structural features of MRTX849/1257 provide additional reversible affinity to the SII-P of KRAS. We observed this directly with NMR spectroscopy, finding that MRTX849/1257 could still engage the SII-P of wild-type KRAS and KRAS(G12D) in the absence of the nucleophilic cysteine. Moreover, previous reports of MRTX849 series SAR support this observation^7^. Inclusion of the chiral cyanomethyl group on the piperazine moiety adjacent to the electrophile dramatically enhances cellular potency by over 2 orders of magnitude, and structural studies suggest that the cyano moiety makes a key hydrogen bond interaction with the backbone nitrogen of Gly10. This enhanced cellular potency for KRAS(G12C) also correlates with a measurable reversible affinity component calculated from kinetic data (reported *K*_I_ of 3.7 μM for KRAS(G12C) and increased specificity constant (*k*_inact_/*K*_I_) for KRAS(G12C)) compared to that reported for AMG510 or ARS-1620. Taken together, it is likely that this cyanomethyl substituent is important for the interaction of MRTX849/1257 with non-G12C KRAS alleles.

Expanding beyond inhibition of KRAS(G12C), the BRET system enabled us to observe engagement of wild-type KRAS and of the majority of KRAS hotspot mutants including G12D. As measured in the BRET system, the rank-order vulnerability of KRAS hotspot mutants to SII-P engagement with MRTX849/1257 and related non-covalent inhibitors (CPD1–6) did not fully correlate with reported rates of intrinsic hydrolysis using purified RAS proteins^12^. Specifically, the G13D, Q61H, and Q61L mutants reportedly have among the lowest intrinsic hydrolysis rates of the hotspot mutants evaluated here, as determined in cell-free systems. Accordingly, these alleles should be among the least vulnerable to SII-P target engagement by GDP-state specific inhibitors, even when considering the potential for steric and conformational effects to confer differential affinity. However, MRTX849/1257 and the non-covalent inhibitors CPD1, CPD2, and CPD6 engaged these mutants nearly as potently as they did WT protein, and significantly more potently than they engaged the G12D and G12V mutants. In the case of G13D, this result may be explained by the high nucleotide exchange rate measured for this mutant.^12,30,31^ In the cases of the Q61 mutants, earlier reports have noted similar discrepancies; a GDP-state specific degrader was able to target KRAS(Q61H) in cells^32^, and KRAS(Q61L) was observed to possess a higher hydrolysis rate in a cellular context than in cell free systems.^31^ These earlier reports and our in-cell BRET data suggest that the nucleotide states of RAS proteins in a cellular setting may deviate from those quantified in a biochemically-defined system, emphasizing the need for direct measurements of target engagement in cells when evaluating RAS-targeted inhibitors. Another factor that we cannot rule out is the potential contribution of RAS proteins in extracellular fractions to the behavior of the cellular BRET assay. RAS proteins in extracellular fractions could potentially contribute to this cellular BRET assay, which may have different properties compared to intracellular RAS proteins.

Notably, as assessed via intracellular BRET and biochemically via NMR, complete engagement of wildtype HRAS was not observed with any of the SII-P inhibitors evaluated here as high as the compound solubility limits. In contrast to wildtype HRAS, HRAS(G12C) engaged the preclinical SII-P inhibitors in BRET assays, albeit with an inverse rank-order to that of KRAS(G12C); MRTX849 and MRTX1257 were far less potent towards HRAS(G12C) compared to KRAS(G12C), while AMG510 engaged both isoforms with similar potency. This suggests that AMG510 could be a potential therapeutic agent for HRAS(G12C)-driven tumors. Residue 95 is the only amino acid in the SII-P binding site to differ between K- and HRAS. In the published co-crystal structure of KRAS(G12C)/MRTX849 (PDB 6UT0), histidine 95 stacks underneath the *N*-methylpyrrolidine substituent and forms a hydrogen bond to the pyrimidine core of the molecule.^7^ In contrast, H95 adopts a very different solvent-exposed conformation within the KRAS(G12C)/AMG510 co-crystal due to the presence of the isopropylpyridine moiety in AMG510 (PDB 6OIM).^5^ In HRAS proteins, the corresponding residue is a glutamine, and the lack of this key interaction in HRAS could explain the contrasting vulnerability of K- and HRAS to MRTX849 and MRTX1257. Furthermore, recent studies have identified secondary mutations in KRAS(G12C) cell lines responsible for acquired resistance to MRTX849 and AMG510 treatment in both clinic-derived tumor samples and in resistance mutagenesis experiments.^11,33,34^ KRAS proteins with mutations at Y96 were found to be resistant to inhibition by both MRTX849 and AMG510, and proteins with mutations at H95 were found to be resistant to inhibition by MRTX849. These resistance studies further support interactions around H95 as critical to the binding of SII-P-targeted inhibitors.

As MRTX849/1257 demonstrate SII-P engagement across KRAS hotspot mutants, this chemotype may serve as the basis for development of allele-specific KRAS inhibitors beyond G12C. However, the GDP-state bias may limit the efficacy of this chemotype against a wider array of KRAS hotspot mutants that may predominantly reside in a GTP state *in vivo*. For example, KRAS(G12D) was only weakly engaged, and KRAS(G12V) and Q61R were largely inaccessible to MRTX849/1257. Thus, more significant chemical modifications will likely be necessary to target these oncogenes, and engaging both nucleotide states of KRAS may be required. At the time of preparing this manuscript, structures of novel KRAS(G12D) inhibitors were disclosed that were structurally similar the MRTX849 chemotype^20–23^. We found that one such example, EX185^23^, can bind GPPNHP-loaded KRAS and KRAS(G12D) 1-169 by NMR and engage KRAS(G12D) in cells by our BRET-based assay with < 100 nM affinity. The increased affinity to KRAS(G12D) translated into potent inhibition of RAF effector interactions as well as potent antiproliferative effect. Though detailed analyses of this new chemotype’s binding mode have not yet been published, its ability to also access the active nucleotide state of KRAS SII-P is likely a key contributor to its increased engagement potency against KRAS(G12D) in cells. Among the KRAS hotspot mutants, KRAS(G12V) is expected to be even more heavily biased towards the GTP-state compared to G12D^12^. Consistent with GTP state accessibility, EX185 engaged KRAS(G12V) in cells with sub-micromolar affinity. Together our target engagement and NMR spectroscopy results support a broad opportunity to target KRAS SII-P in a manner decoupled from nucleotide status.

We have shown that KRAS hotspot mutants offer wider opportunities for SII-P engagement than previously understood; in particular, that some proteins bearing activating mutations may be more accessible to GDP-state inhibition in some cellular contexts than predicted based solely on biochemical GTP hydrolysis rates. Furthermore, recently disclosed chemotypes capable of directly binding the active GTP-loaded state present even wider opportunities for SII-P engagement across KRAS hotspot mutants. Thus, our work highlights the importance of methods to directly assay target engagement in cells to compliment phenotypic assays and *in vitro* biochemical assays. The BRET-based and NMR assays reported in this work provide a reliable workflow to rapidly profile direct target engagement across a variety of RAS hotspot mutants, which should be broadly enabling for SII-P inhibitor discovery. Similarly, these assays may also become important tools to assess KRAS secondary mutations which are already emerging in clinical settings^11,33,34^. These capabilities should aid in the evaluation and optimization of new and improved medicines for RAS-driven cancers and prevalent RASopathies.

## Supporting information

Supplementary Note HSQC

## Acknowledgements

D.M.P. is supported by a Ruth Kirschstein NRSA from the NCI of the NIH (F32CA253966). The content of this publication is solely the responsibility of the authors and does not necessarily represent the official views of the NIH. Q.Z. is the Connie and Bob Lurie Fellow of the Damon Runyon Cancer Research Foundation (DRG-2434-21). Z.Z. is a Damon Runyon Fellow supported by the Damon Runyon Cancer Research Foundation (DRG-2281-17). K.M.S. would like to acknowledge support from the NIH (5R01CA244550), the Mark Foundation for Cancer Research EXTOL, the Samuel Waxman Cancer Research Foundation, and the Howard Hughes Medical Institute. We would also like to thank Dr. Mark Kelly (UCSF) for his advice and assistance with NMR spectroscopy experiments.

## Author contributions

D.M.P. performed and analyzed the data from protein NMR spectroscopy experiments. J.A.W., J.D.V., and C.R.C. designed and synthesized the BRET probe. M.R.T. designed the RAS expression constructs. M.B.R. and J.D.V. designed the in-cell RAS BRET assay system. C.A.Z. performed and analyzed the data from the BRET assays. Q.Z. synthesized the non-covalent SII-P inhibitors. J.D.V. and M.B.R. performed and analyzed the cell-titer glow anti-proliferation experiments. B.F.B. and M.T.B. designed and performed the RAS-RAF interaction assay. Q.Z. and Z.Z. performed and analyzed the data from the ERK phosphorylation assays. K.M.S. and M.B.R. guided the study and supervised the research from their respective groups. J.D.V., D.M.P., Q.Z., Z.Z., K.M.S., and M.B.R. wrote the manuscript.

