## Supplementary Information and Figures for "KRAS is vulnerable to reversible switch-II pocket engagement in cells"

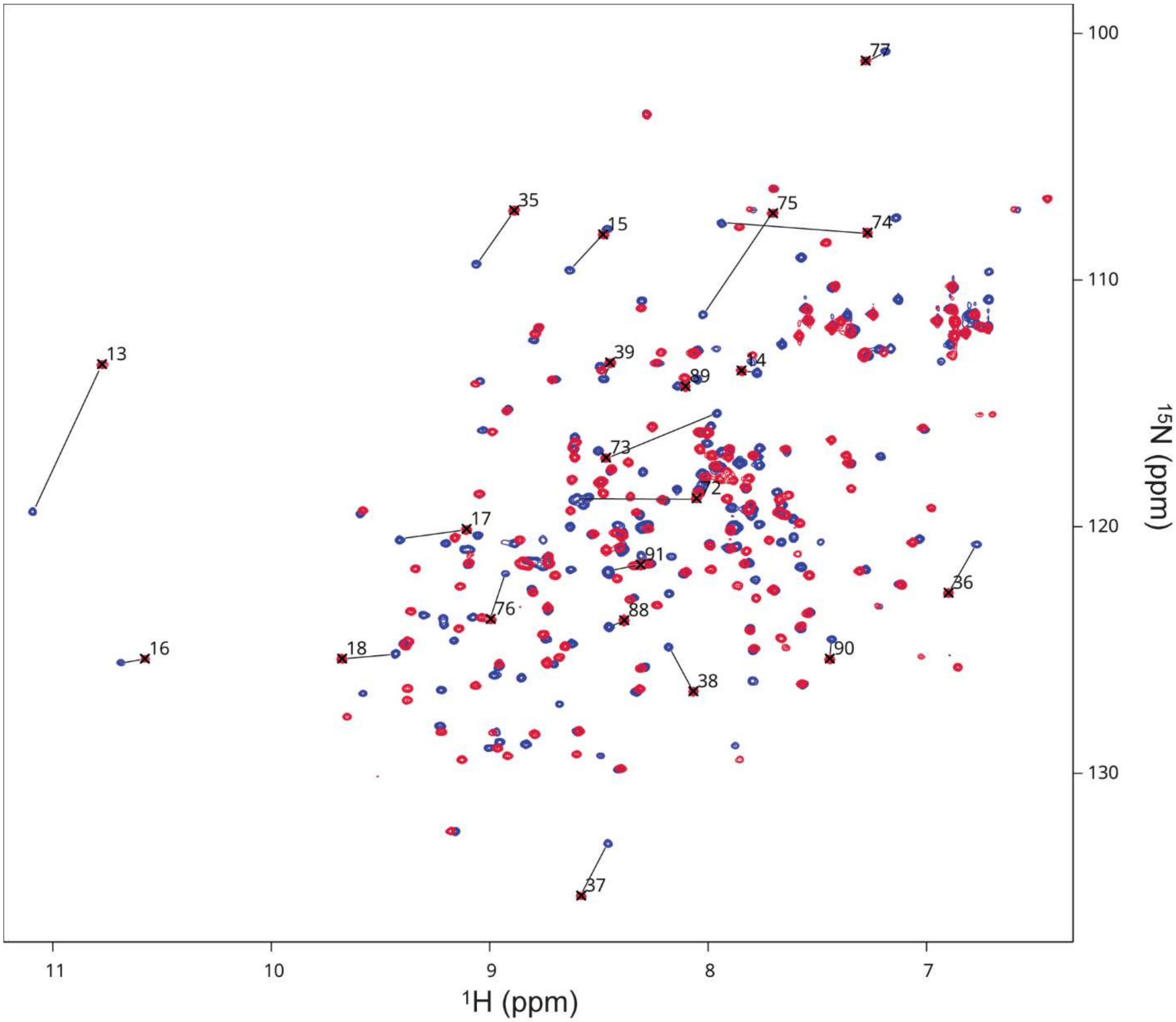

**Figure S1. Effect of MRTX849 on the HSQC NMR spectrum of KRAS(G12C)-GDP.** Superimposed HSQC NMR spectra of 100 µM U-$^{15}$N KRAS(G12C)-GDP 1-169 (blue) and MRTX849-KRAS(G12C)-GDP 1-169 (red) at pH 7.4 and 298 K. Selected peaks corresponding to residues inside the SII-P that could be confidently reassigned to the covalent G12C-MRTX849 adduct based on 3D NOESY-HSQC correlations are noted, and lines connecting them to the analogous peaks of unmodified protein (BMRB 27646) are drawn.

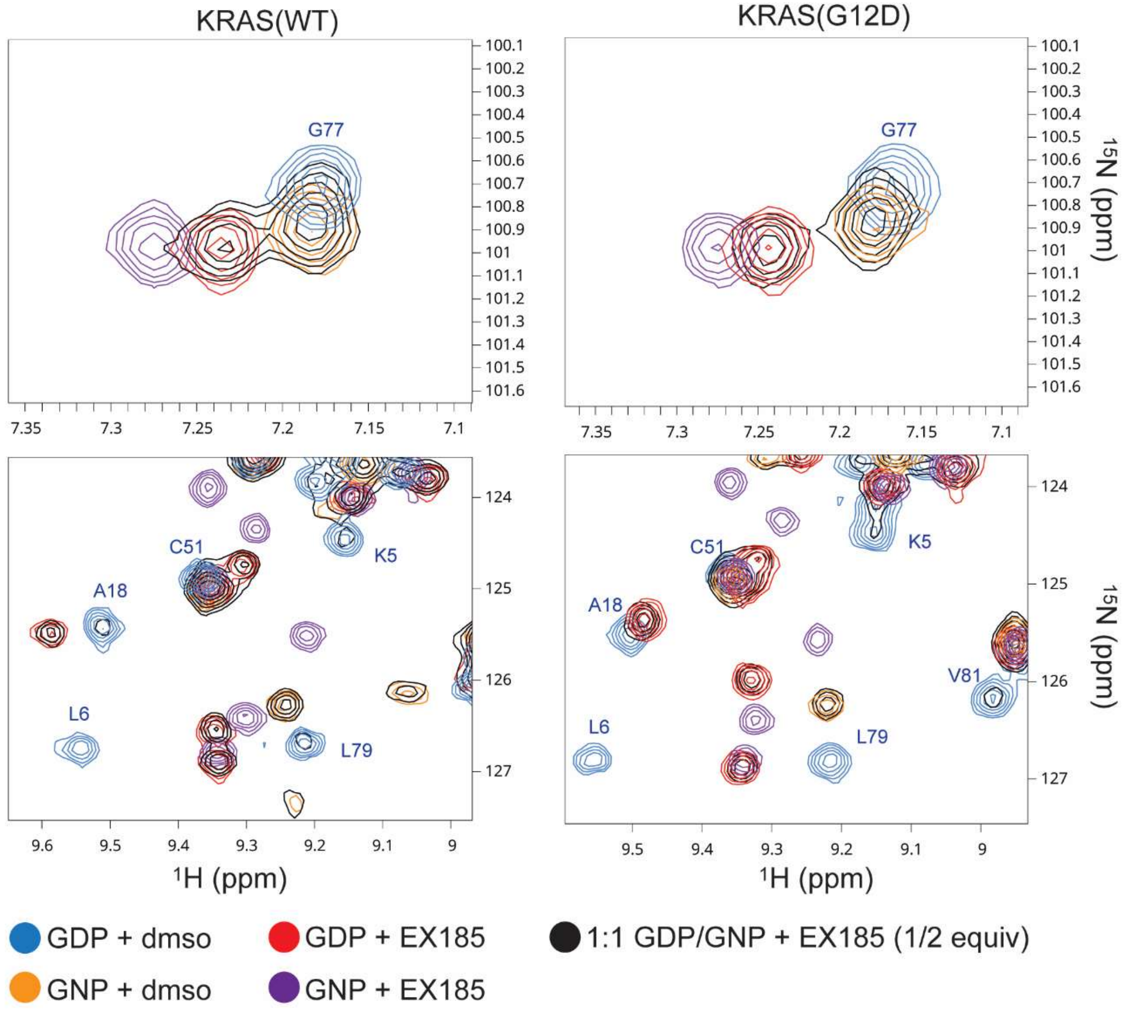

**Figure S2. KRAS nucleotide state preference of EX185.** Superimposed HSQC NMR spectra of 100 µM WT (left) and G12D (right) U-$^{15}$N KRAS-GDP 1-169 with (red) and without (blue) 200 µM EX185, 100 µM U-$^{15}$N KRAS-GNP 1-169 with (purple) and without (orange) 200 µM EX185, and a 1:1 mixture of the two nucleotide states (100 µM each) with 50 µM EX185 (black). Spectra recorded at pH 7.4 and 298 K with 10% dmso-$d_6$. Regions chosen to show examples of peaks that are well-resolved between the four possible complexes. No well-resolved peaks of EX185-KRAS-GNP in the mixed samples could be detected above the noise limit of the spectra (noise <10% of peak heights for most peaks). Assignments of KRAS-GDP are noted in blue text (BMRB 27720 for WT and 27719 for G12D).

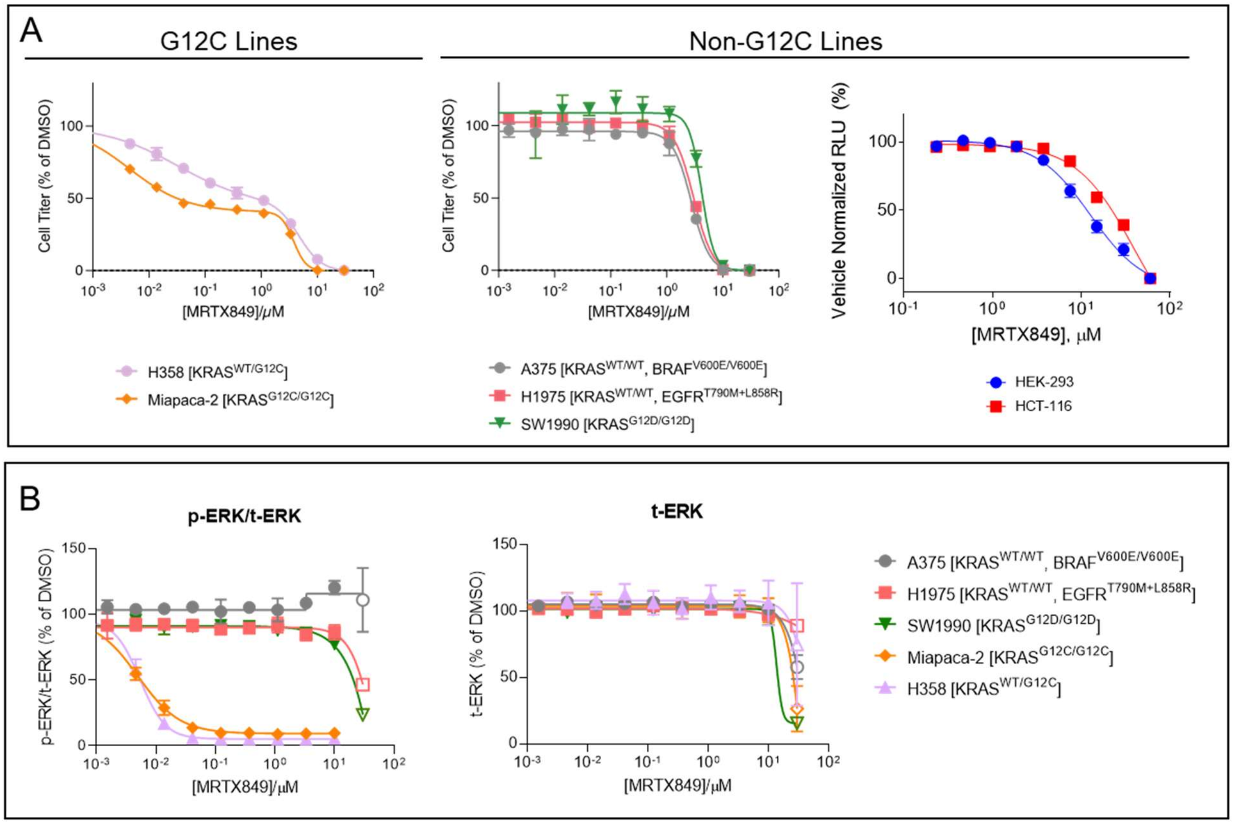

**Figure S3. Characterization of the antiproliferative effects of MRTX849.** A. Cell growth of KRAS(G12C), left, and non-KRAS(G12C) cell lines, right, in the presence of MRTX849 evaluated using CellTiter-Glo. For KRAS(G12C) lines, the curve is fitted to a biphasic inhibitory model with variable hill slopes; for non-KRAS(G12C) lines, the curve curve is fitted to a monophasic inhibitory model with a variable hill slope. For H358, Miapaca-2, A375, H1975, and SW1990 cells, data are mean of at least 3 technical replicates ± S.D. (n=1). For HEK-293 and HCT-116 cells, data are the mean of technical duplicates, (n=1). B. Effect of MRTX849 on phosphorylation of ERK. Left. Phospho-ERK (p-ERK) normalized to total ERK (t-ERK) data across 5 different cell lines. Hollowed data points represent concentrations over which cytotoxic effects are observed as a reduction in t-ERK levels (Right). Data are the mean of technical triplicates (n=1).

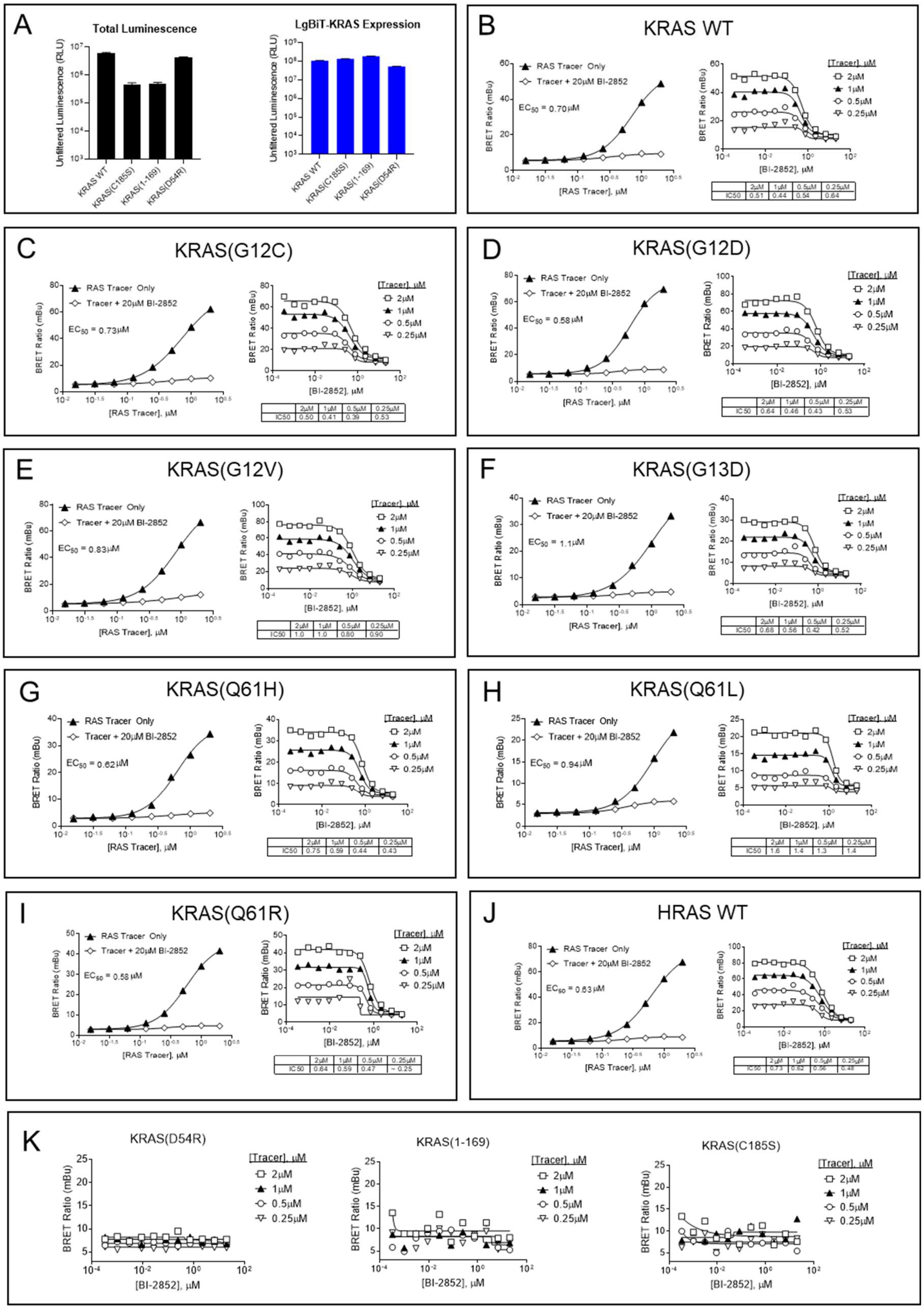
A
Total Luminescence
Unfiltered Luminescence (RLU)
KRAS WT
KRAS(C185S)
KRAS(1-169)
KRAS(D54R)
LgBiT-KRAS Expression
B
KRAS WT
RAS Tracer Only
Tracer + 20μM BI-2852
EC50 = 0.70μM
BRET Ratio (mBu)
[RAS Tracer], μM
[BI-2852], μM
[Tracer], μM
2μM
1μM
0.5μM
0.25μM
IC50 0.51 0.44 0.54 0.64
C
KRAS(G12C)
EC50 = 0.73μM
IC50 0.50 0.41 0.39 0.53
D
KRAS(G12D)
EC50 = 0.58μM
IC50 0.64 0.46 0.43 0.53
E
KRAS(G12V)
EC50 = 0.83μM
IC50 1.0 1.0 0.80 0.90
F
KRAS(G13D)
EC50 = 1.1μM
IC50 0.68 0.56 0.42 0.52
G
KRAS(Q61H)
EC50 = 0.62μM
IC50 0.75 0.59 0.44 0.43
H
KRAS(Q61L)
EC50 = 0.94μM
IC50 1.6 1.4 1.3 1.4
I
KRAS(Q61R)
EC50 = 0.58μM
IC50 0.64 0.59 0.47 ~ 0.25
J
HRAS WT
EC50 = 0.63μM
IC50 0.73 0.62 0.56 0.48
K
KRAS(D54R)
KRAS(1-169)
KRAS(C185S)

**Figure S4. Characterization of SI/II-P BRET assay for RAS in live cells.** The N-terminus of each HRAS or KRAS4B variant was tagged with complementation-based NanoBiT (SmBiT and LgBiT) reporters [1,2]. Full-length RAS constructs with intact C-terminu were utilized to preserve lipidation via the native membrane localization domain. Co-expression of full length LgBiT-RAS with SmBiT-RAS generated a luminescent signal selectively at homo-multimeric RAS compexes in cells. A. Luminescent complexes are observed for KRAS dimers (left). Removal of the HVR or mutation at C185S results in a loss of luminescence, supporting RAS lipidation signals as critical to assay signal. Mutation at D54Q was used to confirm dimerization and SII/I binding. Similar expression levels were observed for the LgBiT fusions under each condition (right) after detection with HiBiT peptide. Results are mean ± S.D. of 12 technical replicates. B-J. Characterization of BRET between RAS dimers and the SI/II-P BRET probe for wildtype KRAS and HRAS, as well as KRAS hotspot mutants. Titration of BI-2852 confirms specific BRET in live cells (n=1). K. D54R and mutations to critical lipidation sites on KRAS are essential for the BRET signal with the SI/II-P BRET probe (n=1).

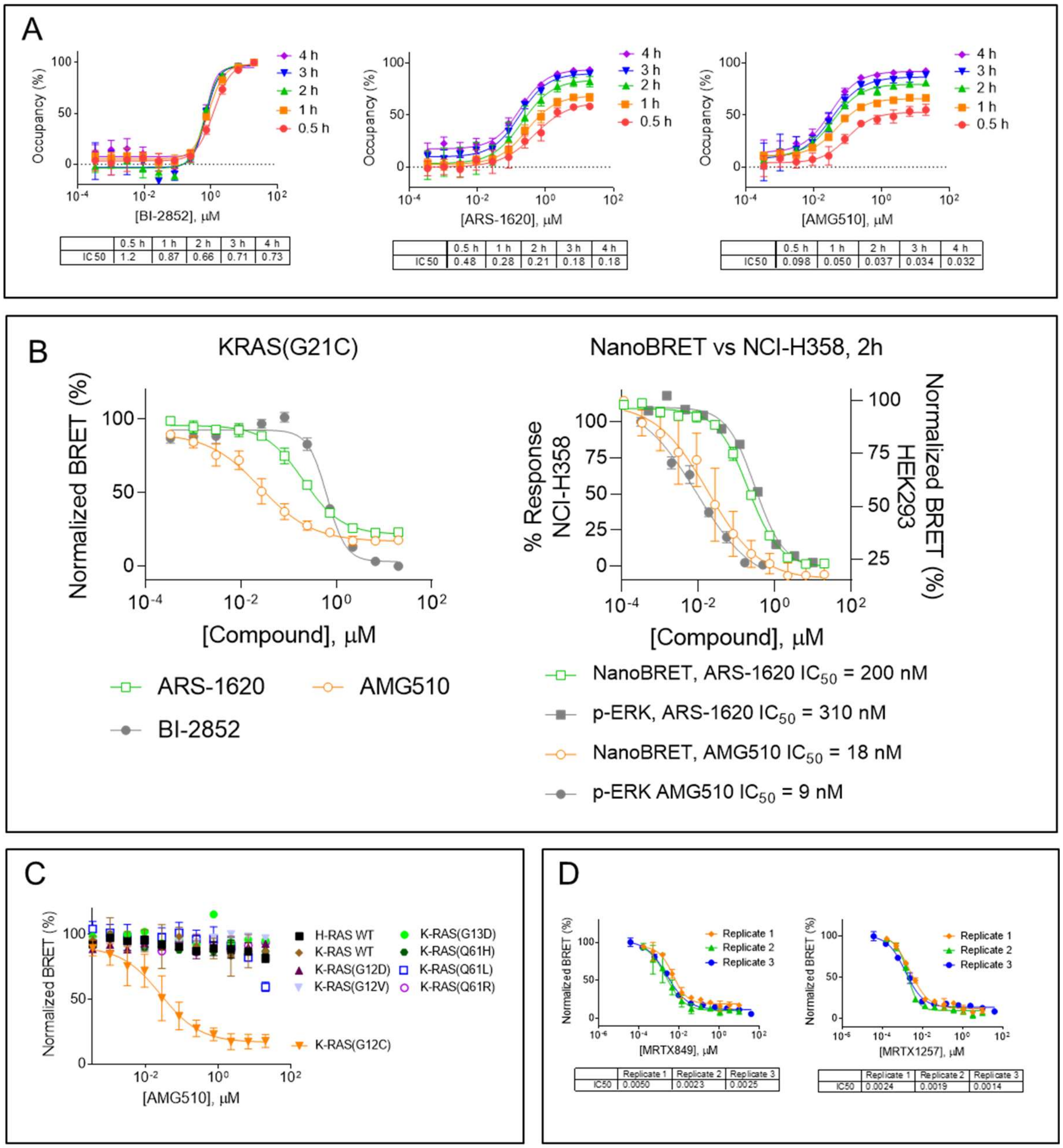

**Figure S5. Characterization of SII-P engagement at KRAS(G12C).** A. Protracted KRAS(G12C) engagement is observed for SII-P ligands, while rapid engagement is observed for BI-2852. Individual data points are the mean ± S.D. of 4 technical replicates (n=2). B. Left: BI-2852 as well as SII-P ligands ARS-1620 and AMG510 engage KRAS(G12C) in live cells. Right: Comparison of target engagement versus phospho-ERK as reported by Canon et al.[3] Data are mean of 4 independent experiments ± S.E.M. (n=4). C. Live cell BRET target engagement analysis with AMG510 confirming selectivity for KRAS(G12C) versus other RAS variants. Results are the mean ± S.E. of 1−4 independent experiments (N = 1−4). D. Target engagement is observed with MRTX849 and MRTX1257 at KRAS(G12C) in live cells. Individual data points are mean ± S.D of 4 technical replicates (n = 3).

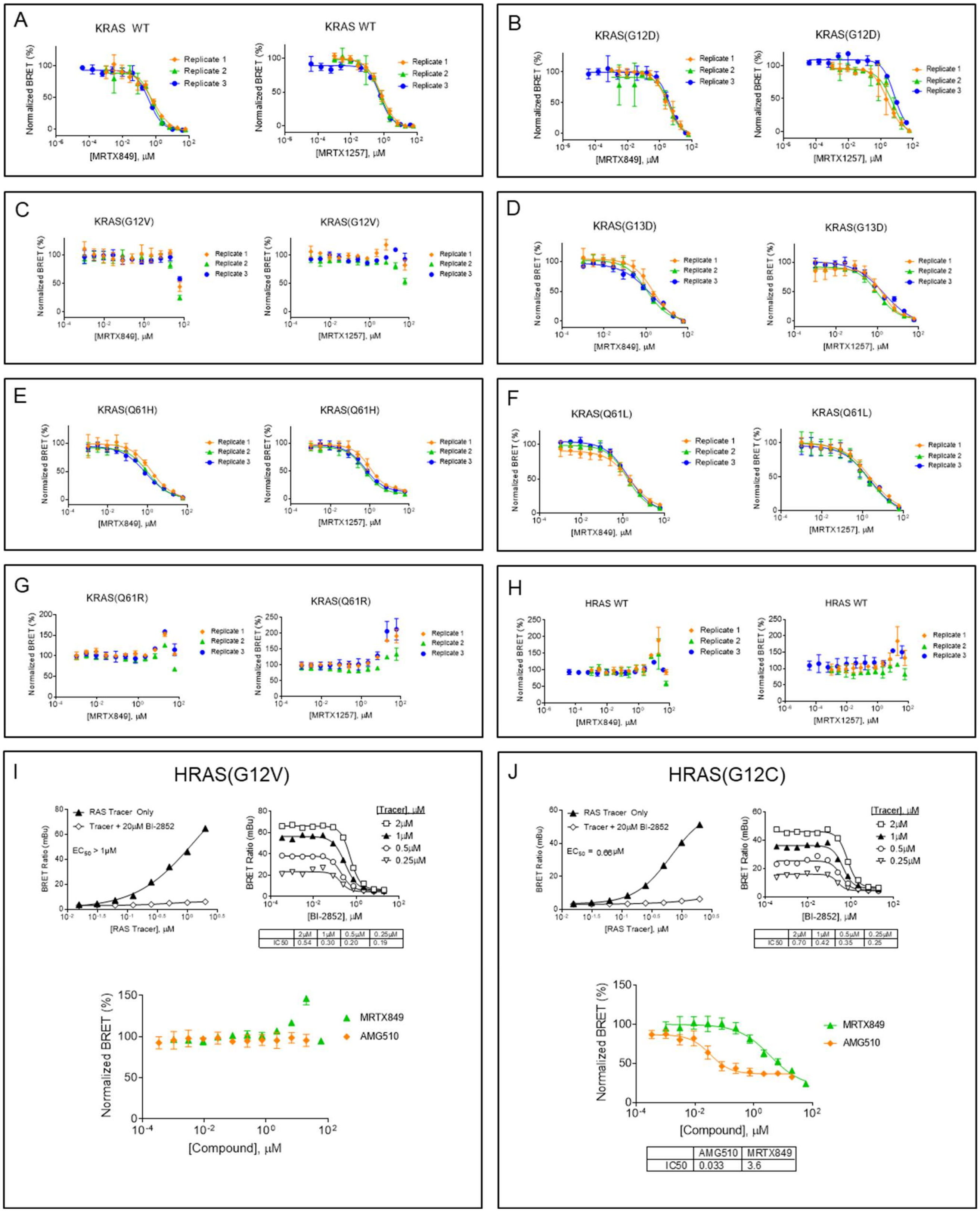

| | 2μM | 1μM | 0.5μM | 0.25μM |
|---|---|---|---|---|
| IC50 | 0.54 | 0.30 | 0.20 | 0.19 |

| | 2μM | 1μM | 0.5μM | 0.25μM |
|---|---|---|---|---|
| IC50 | 0.70 | 0.42 | 0.35 | 0.25 |

| | AMG510 | MRTX849 |
|---|---|---|
| IC50 | 0.033 | 3.6 |

**Figure S6. Observation of engagement of KRAS and key hotspot mutants with MRTX849 and MRTX1257.** A−H. Reproducibility of MRTX849 and MRTX1257 engagement across RAS isoforms. Data are mean ± S.D. of 4 technical replicates (n = 3). I, J. Characterization of BRET assay (n = 1) and SII-P engagement for HRAS(G12V) and HRAS(G12C). Data for tracer binding and BI-2852 engagement are technical singlicates (n = 1), and data for MRTX849 and AMG510 engagement are the mean ± S.D. of 4 technical replicates (n = 1).

A

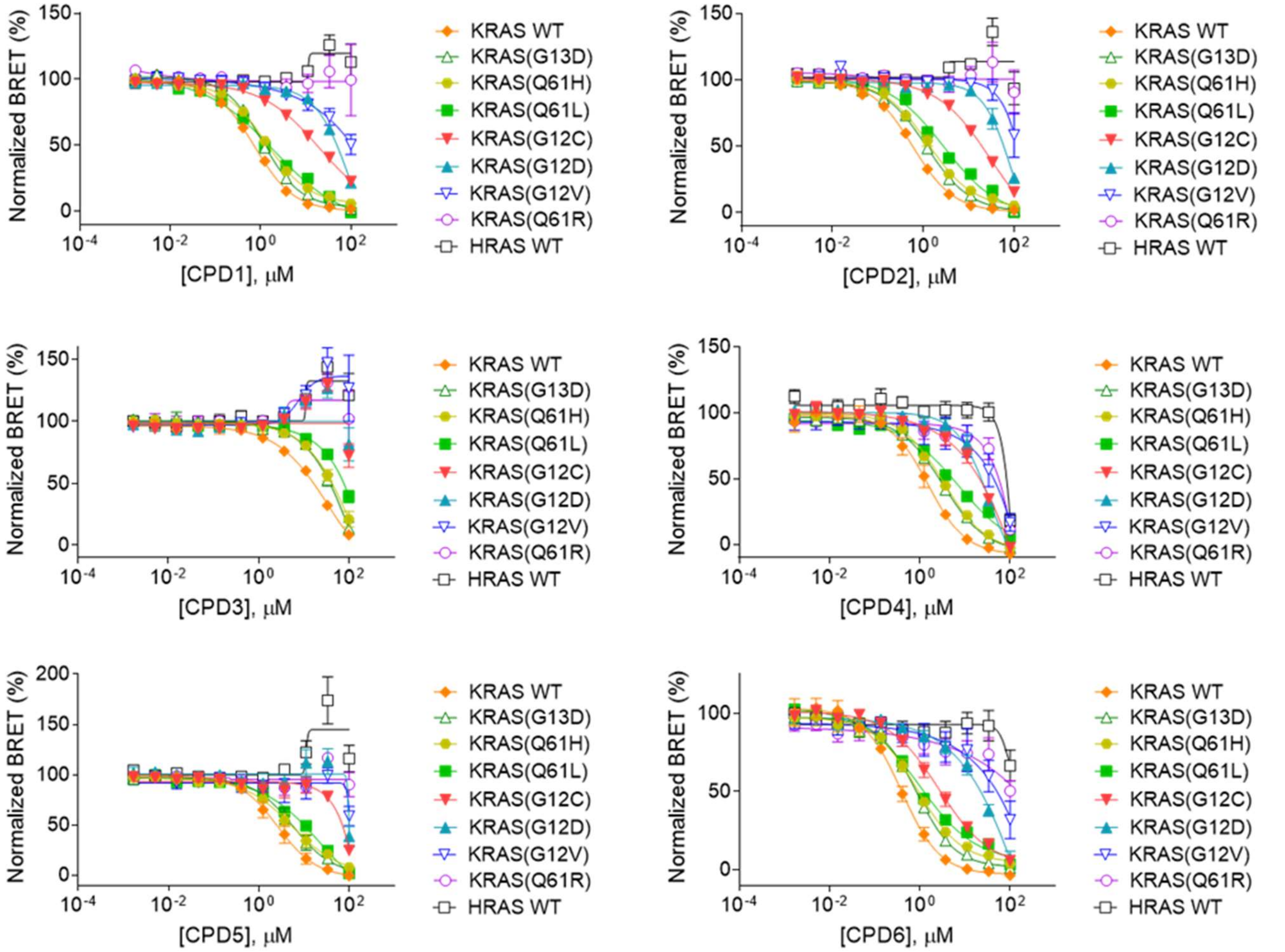

B

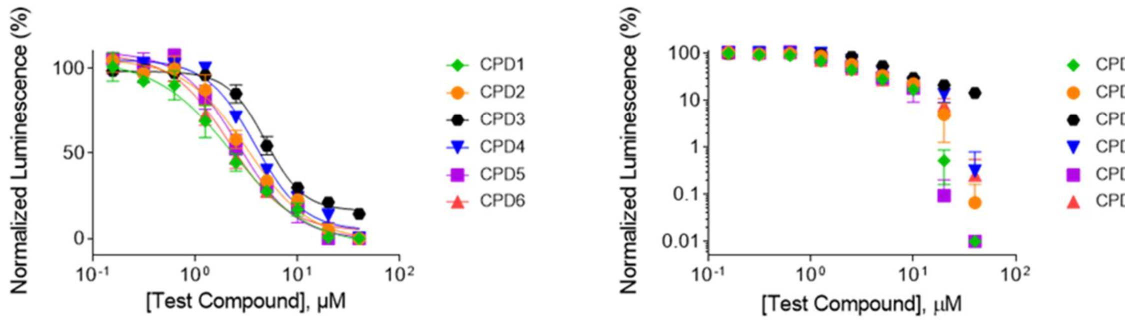

| | CPD1 | CPD2 | CPD3 | CPD4 | CPD5 | CPD6 |
|---|---|---|---|---|---|---|
| IC50 | 2.3 | 3.2 | 4.9 | 3.8 | 2.7 | 2.1 |

| | CPD1 | CPD2 | CPD3 | CPD4 | CPD5 | CPD6 |
|---|---|---|---|---|---|---|
| IC50 | 2.3 | 3.2 | 4.9 | 3.8 | 2.7 | 2.1 |

**Figure S7. Characterization of RAS engagement for reversible saturated amides of MRTX849.** A. BRET target engagement profiles of CPD1−6 for KRAS and HRAS, as well as KRAS hotspot mutants. Data are mean of 3 independent experiments, each performed with 4 technical replicates ± S.E.M. (n=3). B. Antiproliferative/cytotoxic effect of 1−6 in RAS-independent HEK293 cells using CellTiter-Glo. Data are depicted on a linear (left) and a log (right) scale to better visualize the biphasic activity of CPD1−6. Data are mean of technical triplicates (n=1).

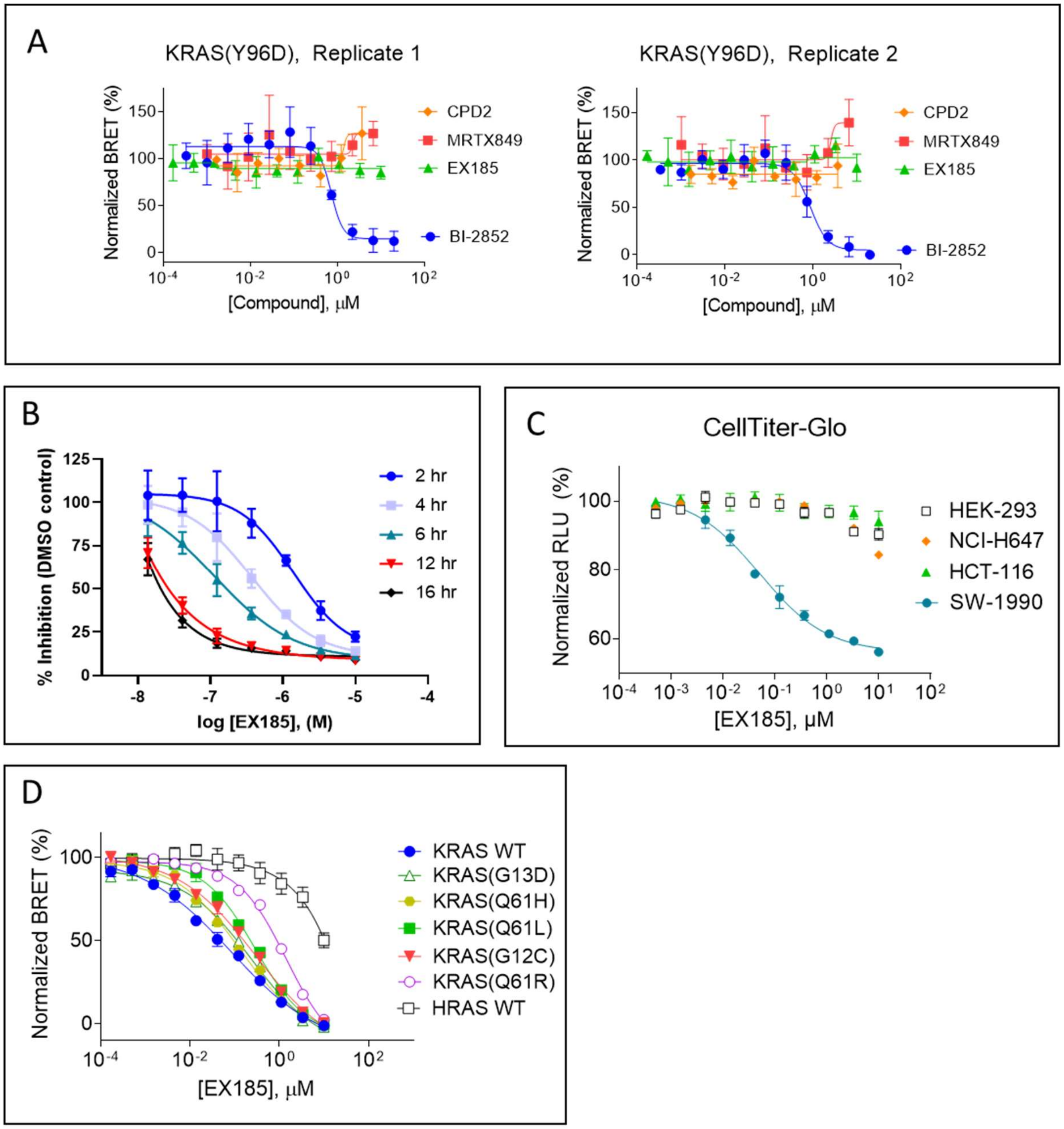

**Figure S8. Characterization of EX185.** A. Characterization of engagement of compounds to KRAS(Y96D). Individual data are mean of 4 technical replicates ± S.D. (n=2). B. Inhibition of KRAS(G12D) : CRAF(RBD) in live cells using NanoBiT. Individual data points are the mean ± S.D. of 6 technical replicates (n=1). C. Anti-proliferative effects of EX185 in various cell lines. Data are mean of 3 independent experiments, each performed with at least 3 technical replicates ± S.E.M. (n=3). D. Engagement of EX185 to KRAS hotspot mutants. For KRAS WT, individual data points are the mean ± S.E. of 4 independent experiments (n=4). For all other RAS variants, individual data points are the mean of 4 technical replicates ± S.D. (n=1).
