## Supplementary Note HSQC for "KRAS is vulnerable to reversible switch-II pocket engagement in cells"

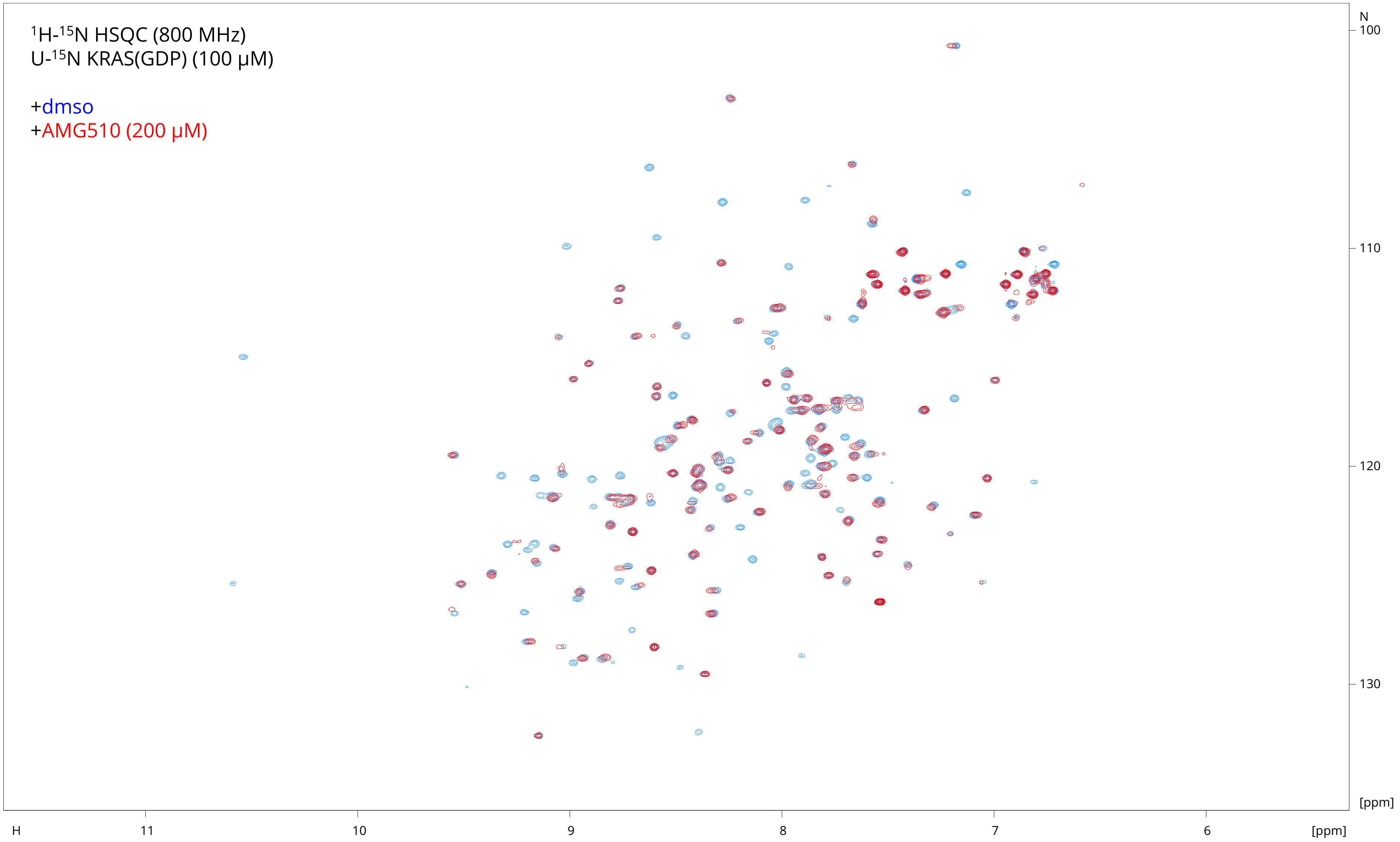

$^1$H-$^{15}$N HSQC (800 MHz)
U-$^{15}$N KRAS(GDP) (100 µM)
+dmso
+AMG510 (200 µM)
N
100
110
120
130
[ppm]
H
11
10
9
8
7
6
[ppm]

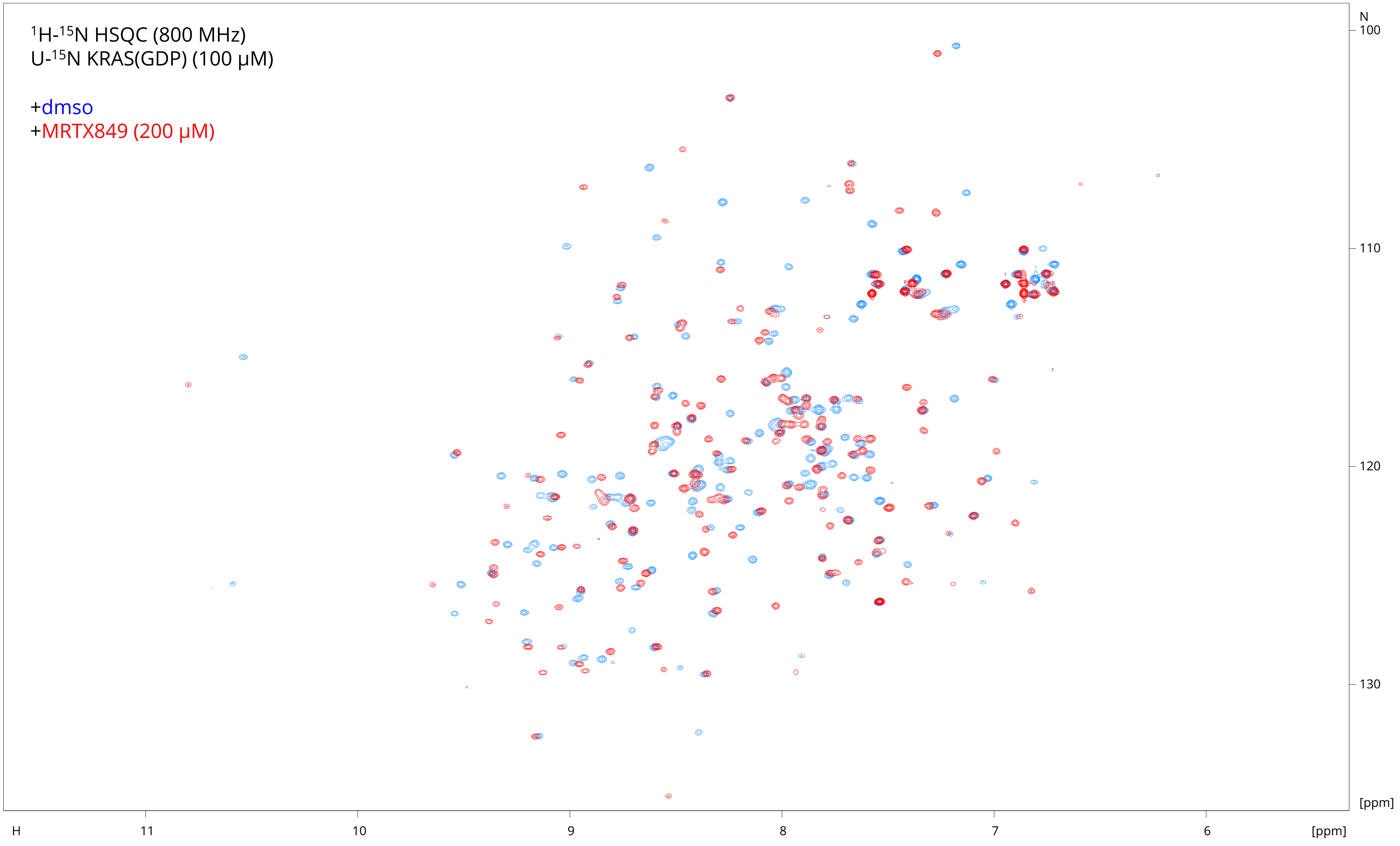

1H-15N HSQC (800 MHz)
U-15N KRAS(GDP) (100 µM)
+dmso
+MRTX849 (200 µM)
N
100
110
120
130
[ppm]
H
11
10
9
8
7
6
[ppm]

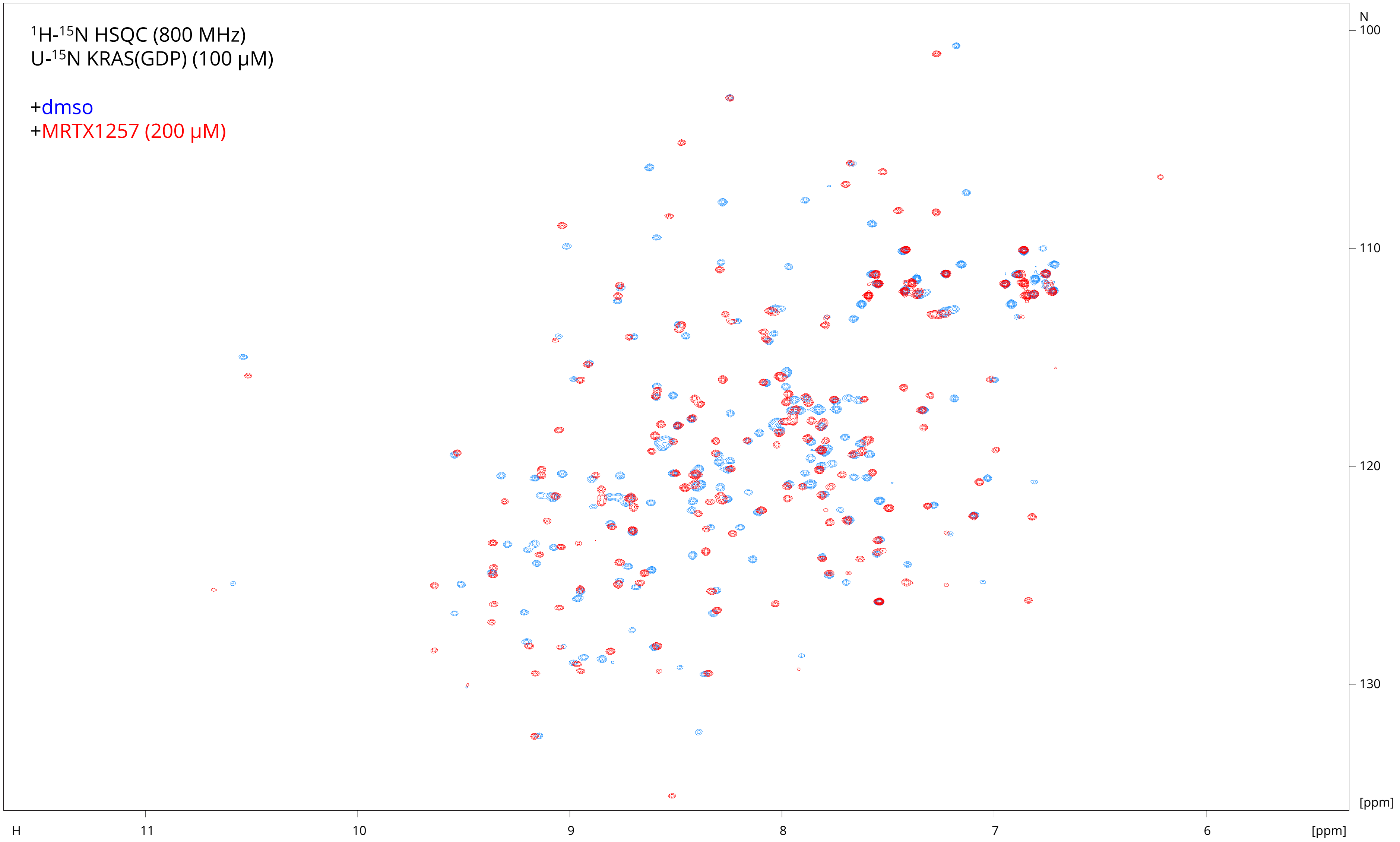

1H-15N HSQC (800 MHz)
U-15N KRAS(GDP) (100 μM)
+dmso
+MRTX1257 (200 μM)
N
100
110
120
130
[ppm]
H
11
10
9
8
7
6
[ppm]

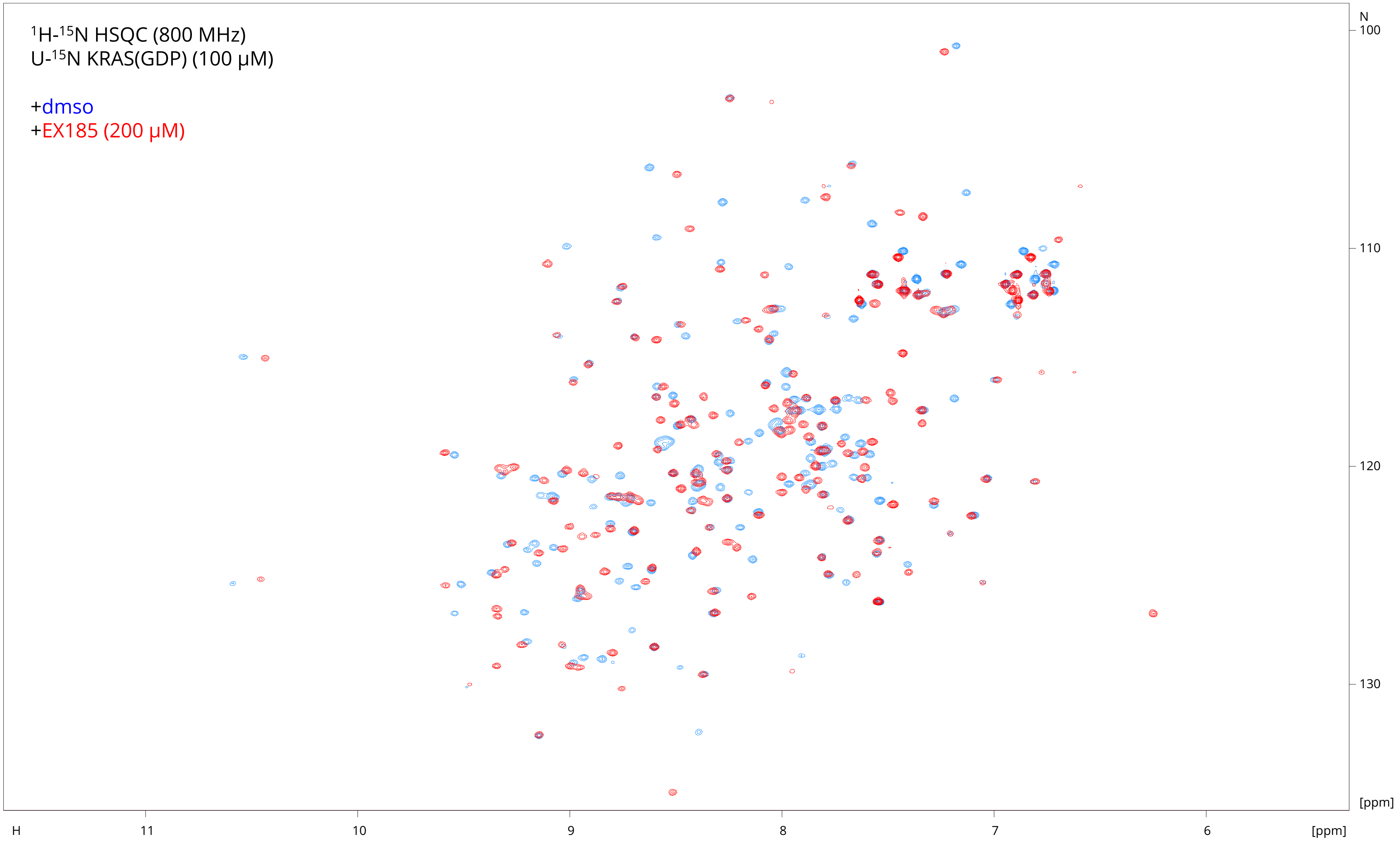

$^{1}H$-$^{15}N$ HSQC (800 MHz)
U-$^{15}N$ KRAS(GDP) (100 µM)
+dmso
+EX185 (200 µM)
N
100
110
120
130
[ppm]
H
11
10
9
8
7
6
[ppm]

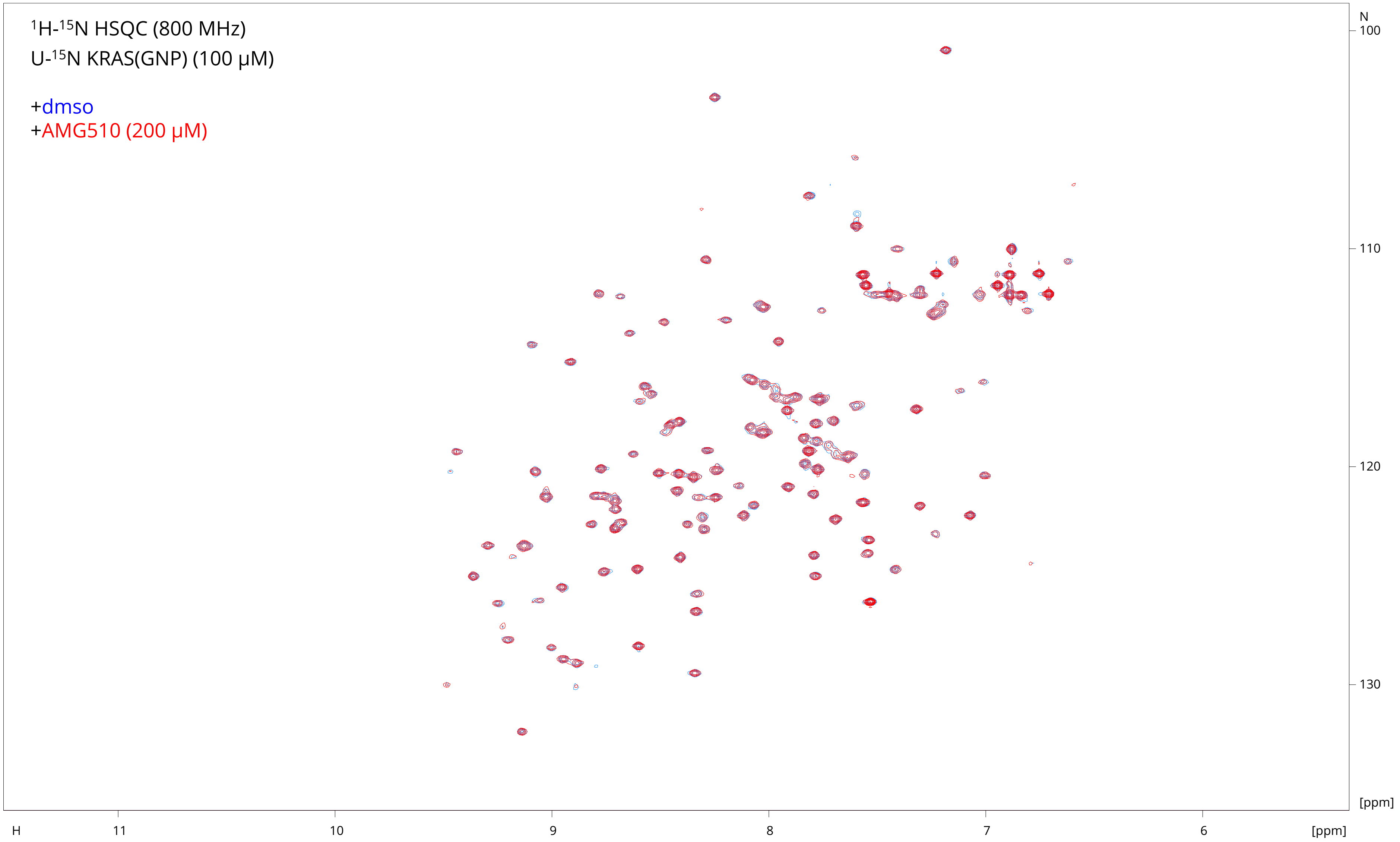

$^{1}H$-$^{15}N$ HSQC (800 MHz)
U-$^{15}N$ KRAS(GNP) (100 µM)
+dmso
+AMG510 (200 µM)
N
100
110
120
130
[ppm]
H
11
10
9
8
7
6
[ppm]

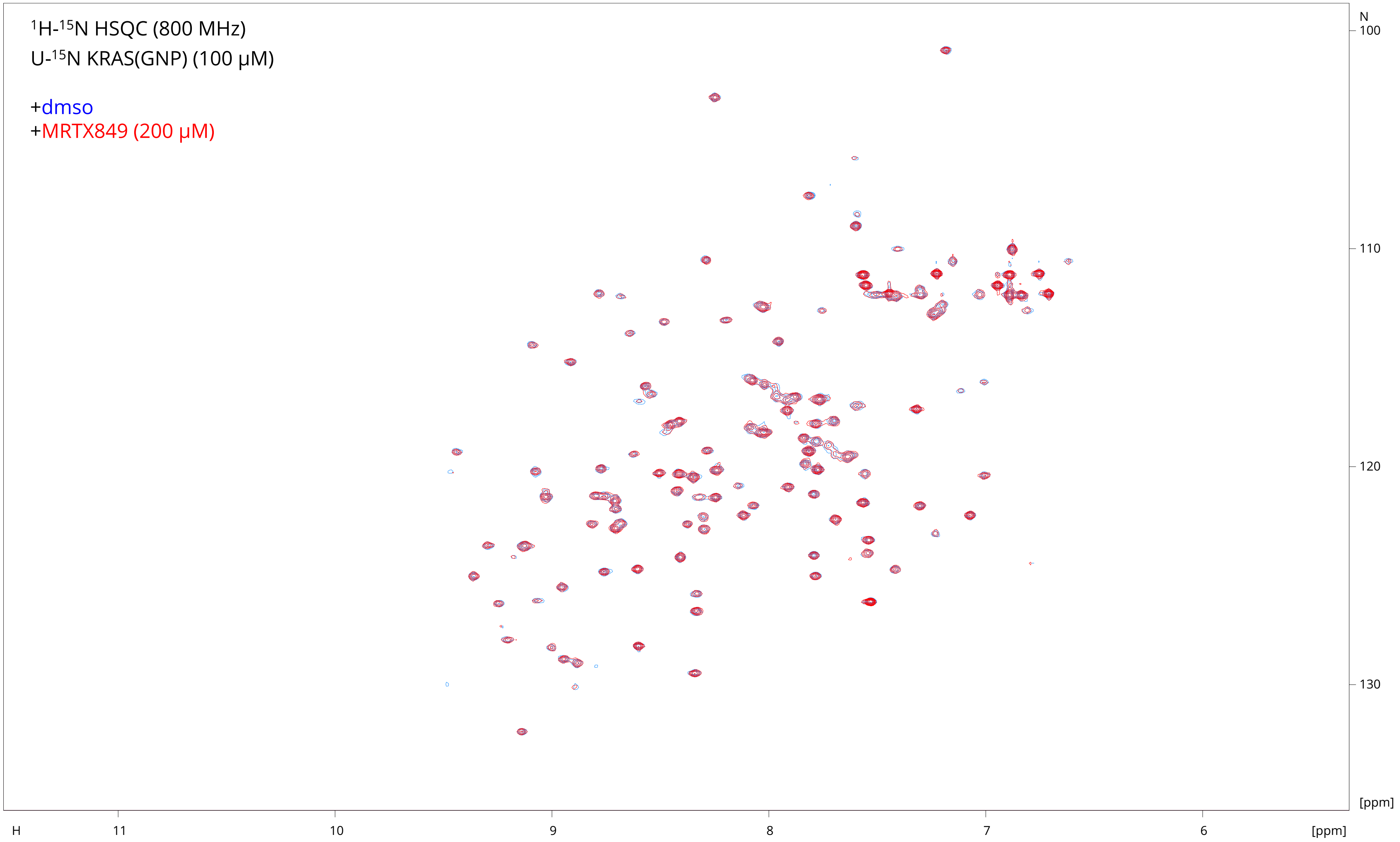

$^{1}H$-$^{15}N$ HSQC (800 MHz)
U-$^{15}N$ KRAS(GNP) (100 µM)
+dmso
+MRTX849 (200 µM)
N
100
110
120
130
[ppm]
H
11
10
9
8
7
6
[ppm]

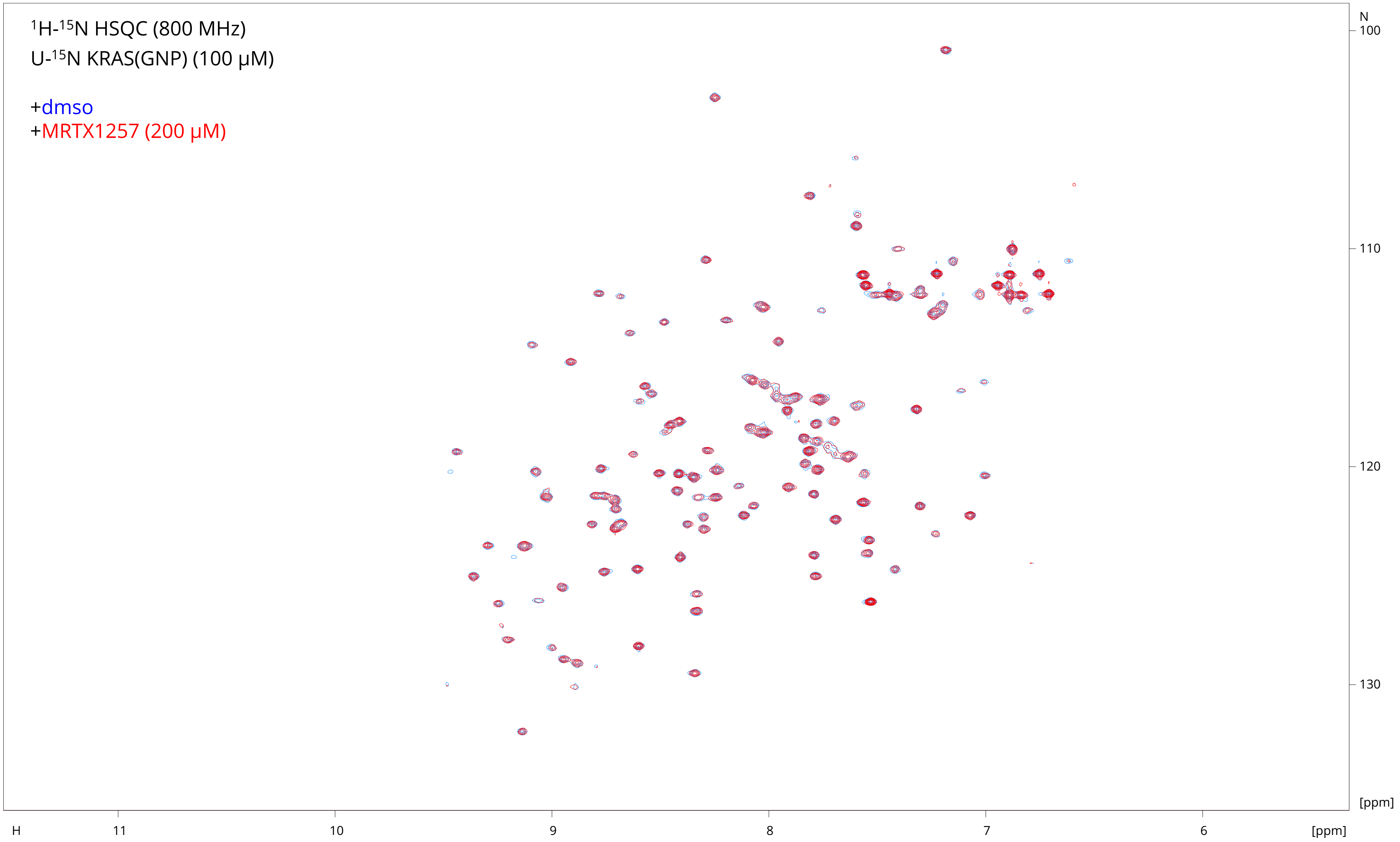

$^{1}H$-$^{15}N$ HSQC (800 MHz)
U-$^{15}N$ KRAS(GNP) (100 µM)
+dmso
+MRTX1257 (200 µM)
N
100
110
120
130
[ppm]
H
11
10
9
8
7
6
[ppm]

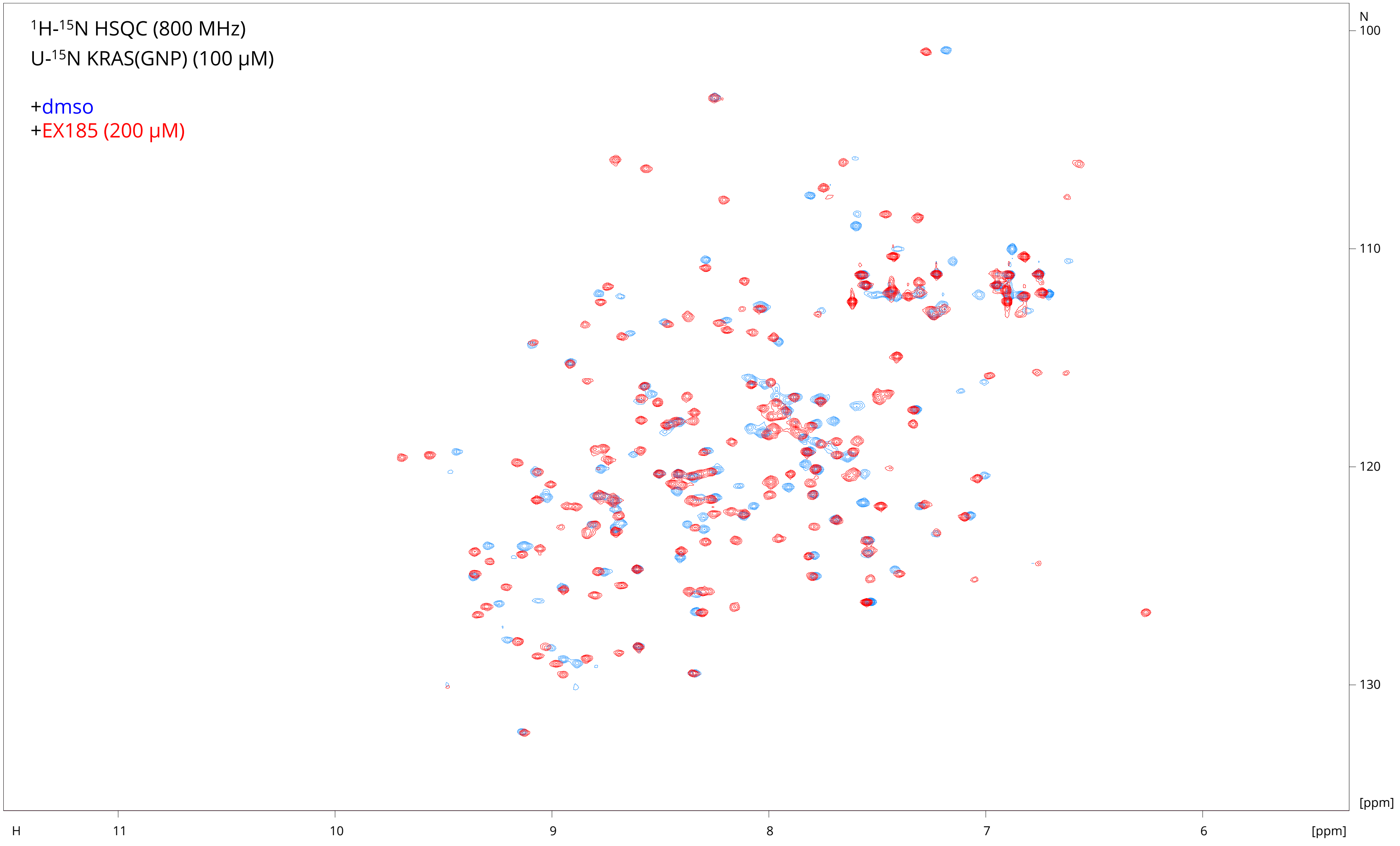

$^{1}$H-$^{15}$N HSQC (800 MHz)
U-$^{15}$N KRAS(GNP) (100 μM)
+dmso
+EX185 (200 μM)
N
100
110
120
130
[ppm]
H
11
10
9
8
7
6
[ppm]

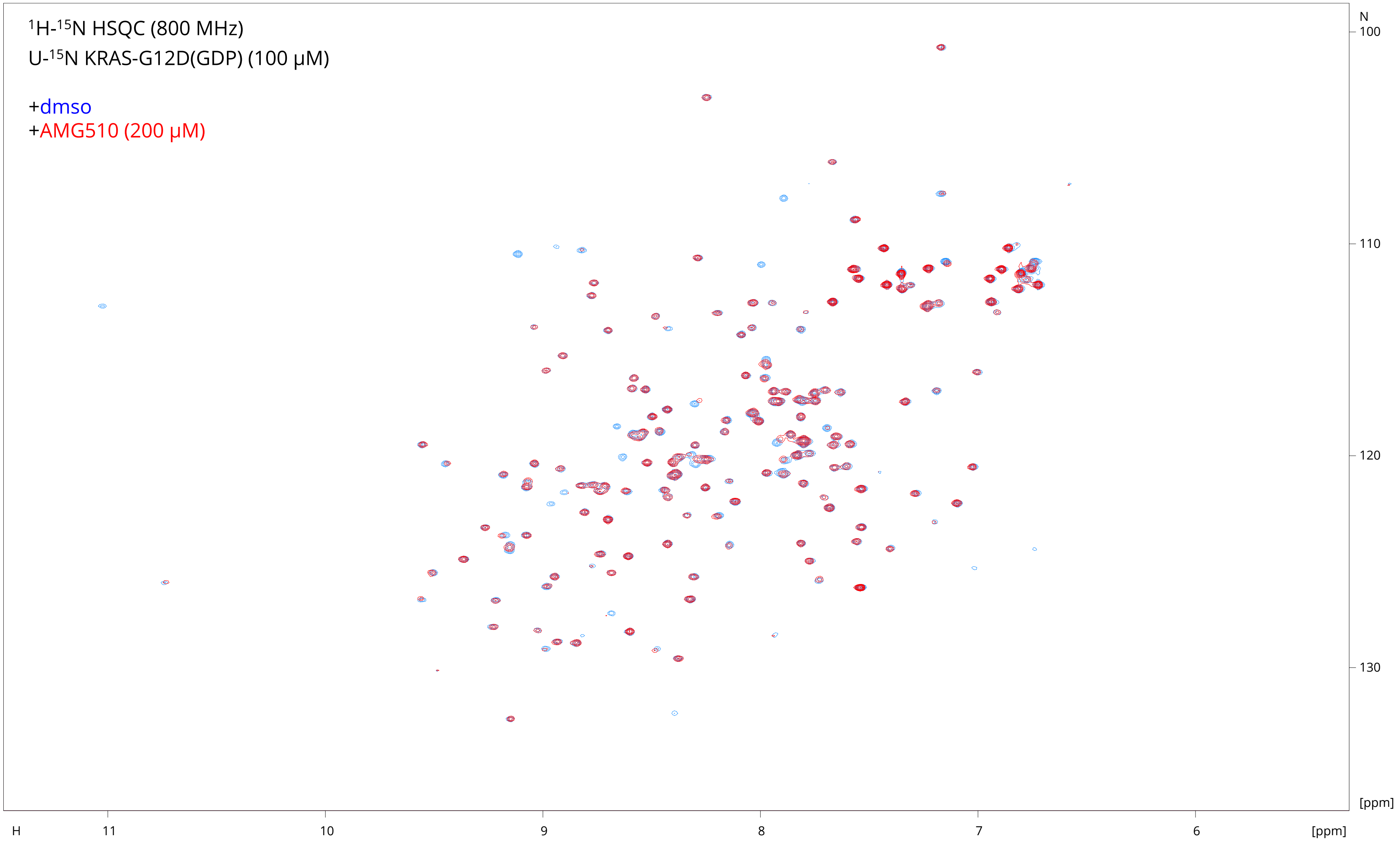

1H-15N HSQC (800 MHz)
U-15N KRAS-G12D(GDP) (100 µM)
+dmso
+AMG510 (200 µM)
N
100
110
120
130
[ppm]
H
11
10
9
8
7
6
[ppm]

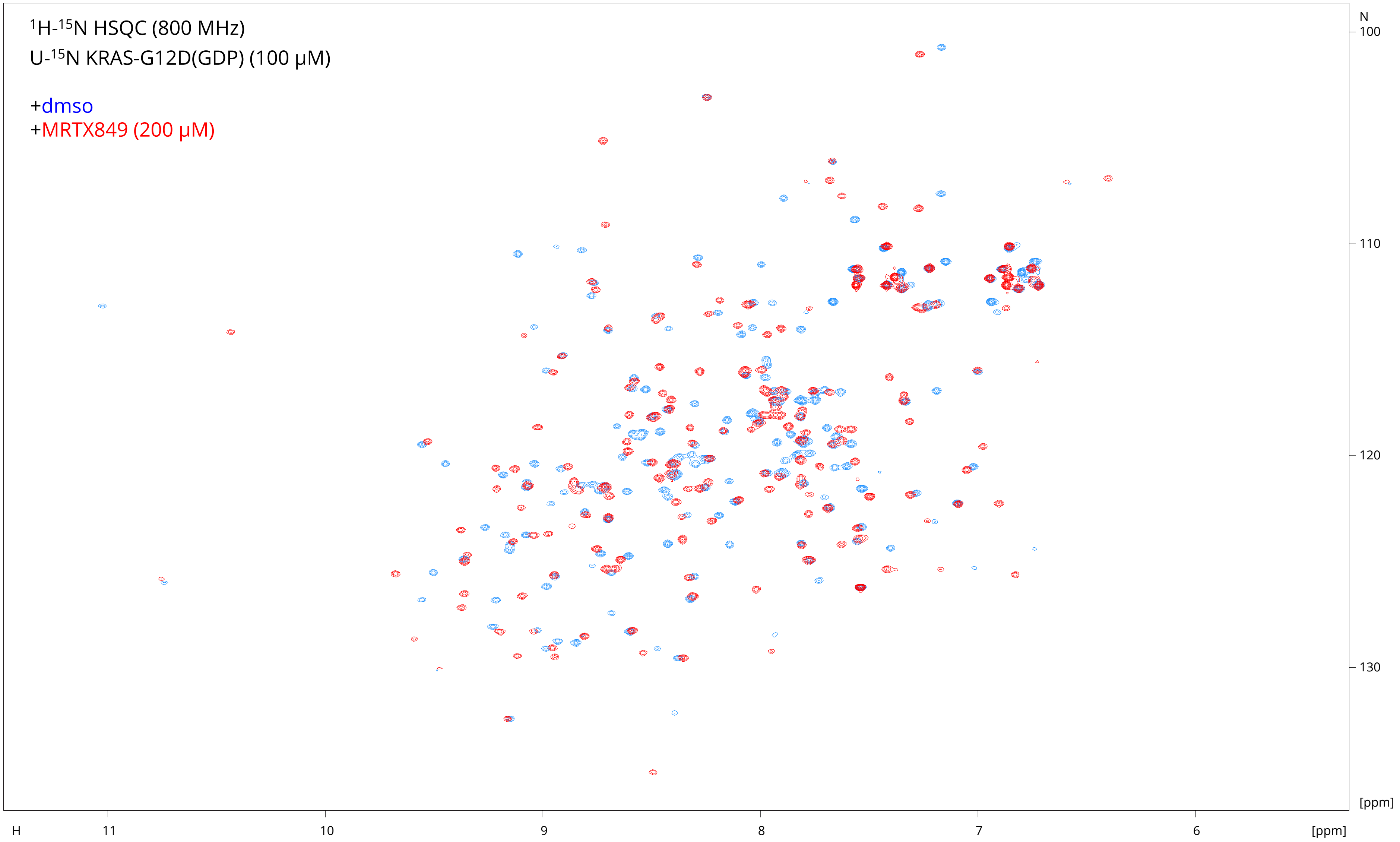

1H-15N HSQC (800 MHz)
U-15N KRAS-G12D(GDP) (100 μM)
+dmso
+MRTX849 (200 μM)
N
100
110
120
130
[ppm]
H
11
10
9
8
7
6
[ppm]

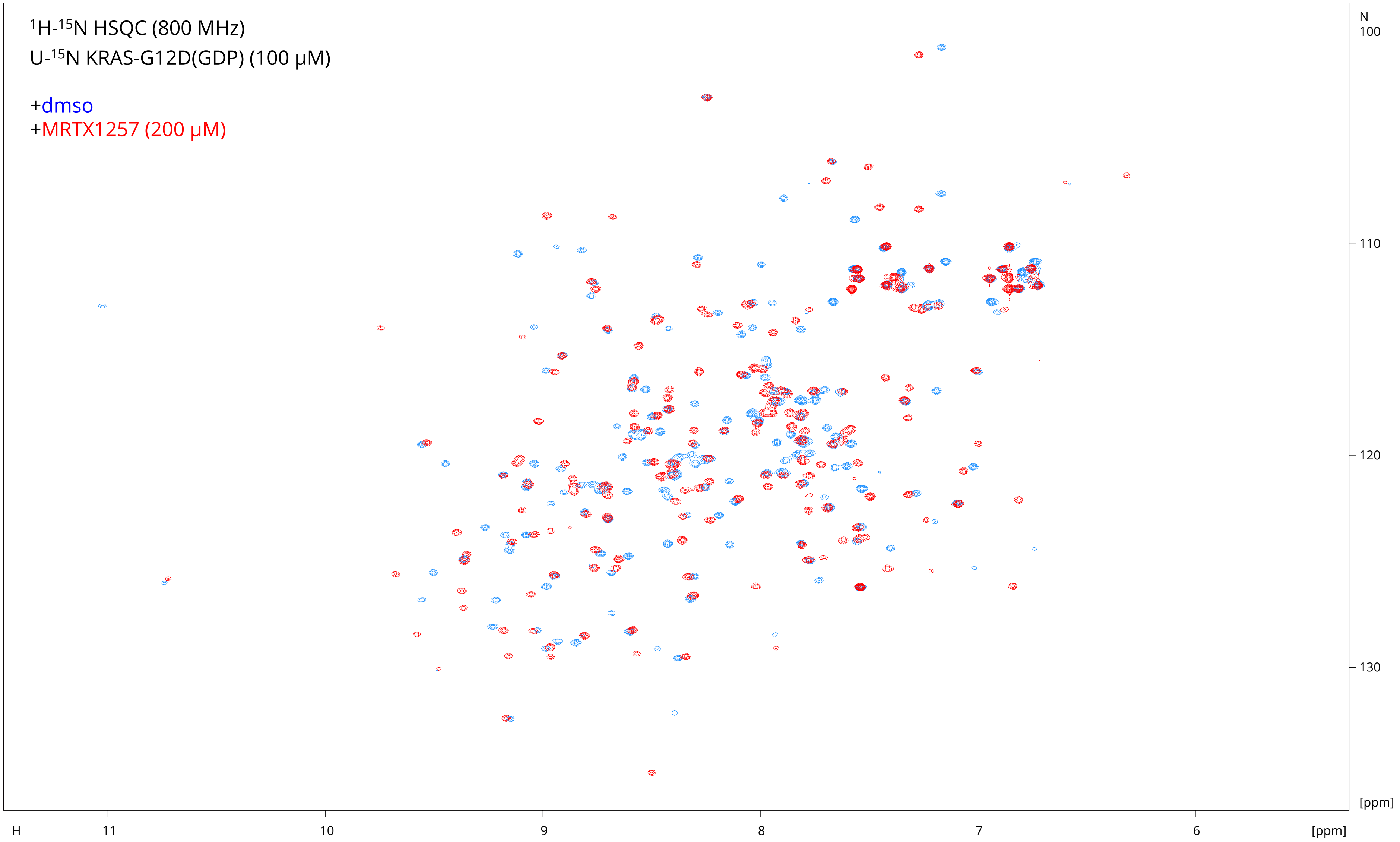

1H-15N HSQC (800 MHz)
U-15N KRAS-G12D(GDP) (100 μM)
+dmso
+MRTX1257 (200 μM)
N
100
110
120
130
[ppm]
H
11
10
9
8
7
6
[ppm]

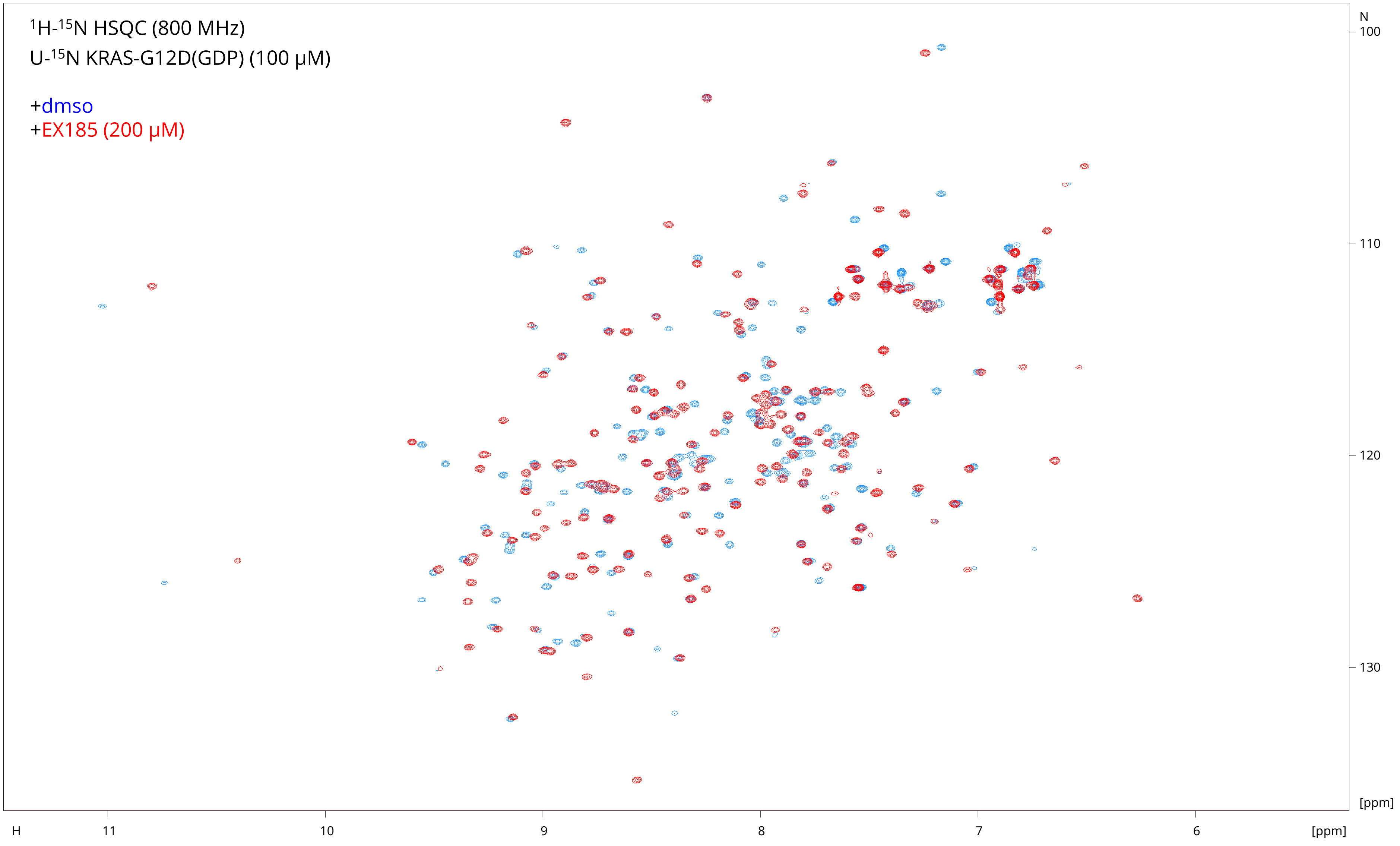

$^1H$-$^{15}N$ HSQC (800 MHz)
U-$^{15}N$ KRAS-G12D(GDP) (100 μM)
+dmso
+EX185 (200 μM)
N
100
110
120
130
[ppm]
H
11
10
9
8
7
6
[ppm]

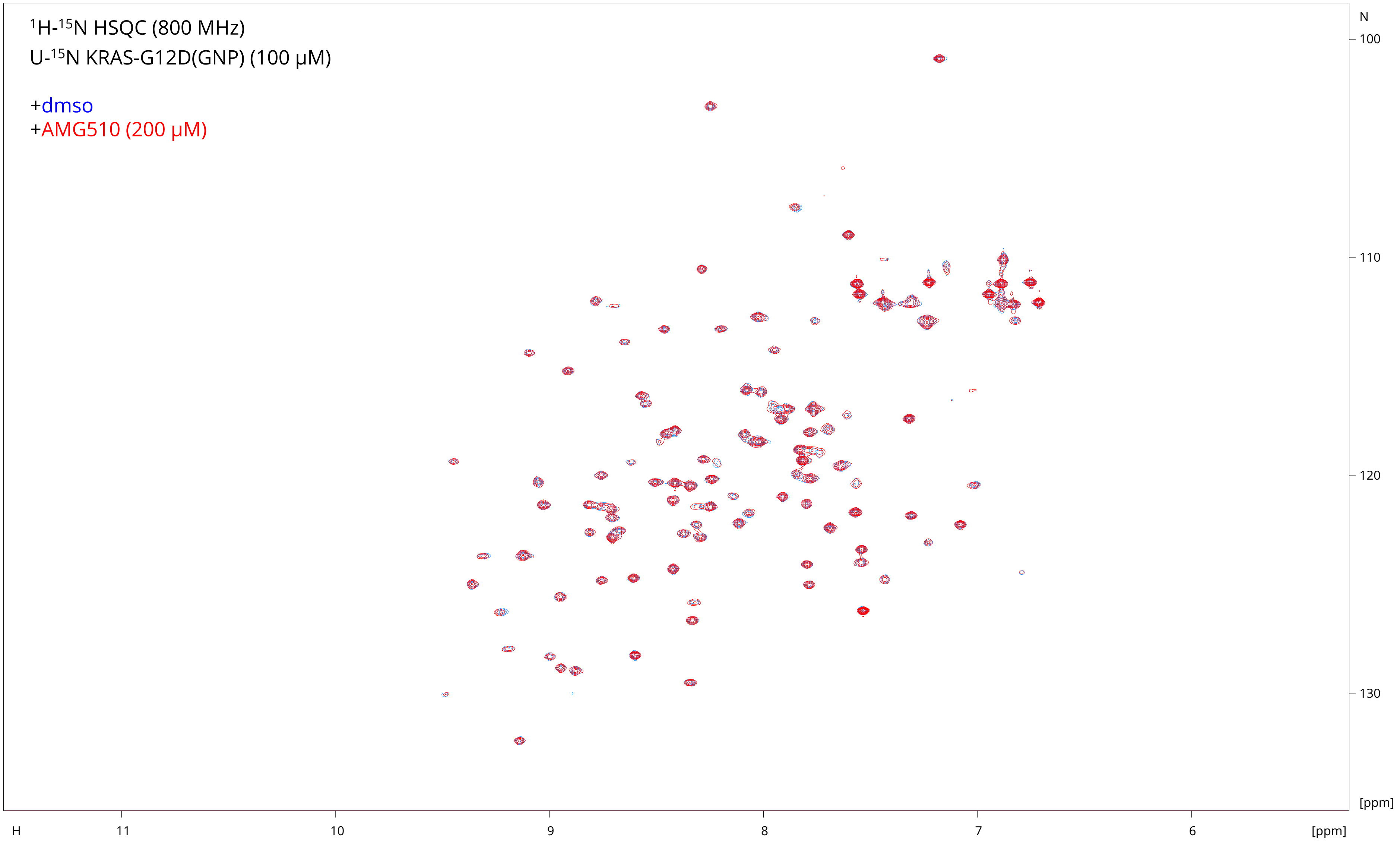

¹H-¹⁵N HSQC (800 MHz)
U-¹⁵N KRAS-G12D(GNP) (100 µM)
+dmso
+AMG510 (200 µM)
N
100
110
120
130
[ppm]
H
11
10
9
8
7
6
[ppm]

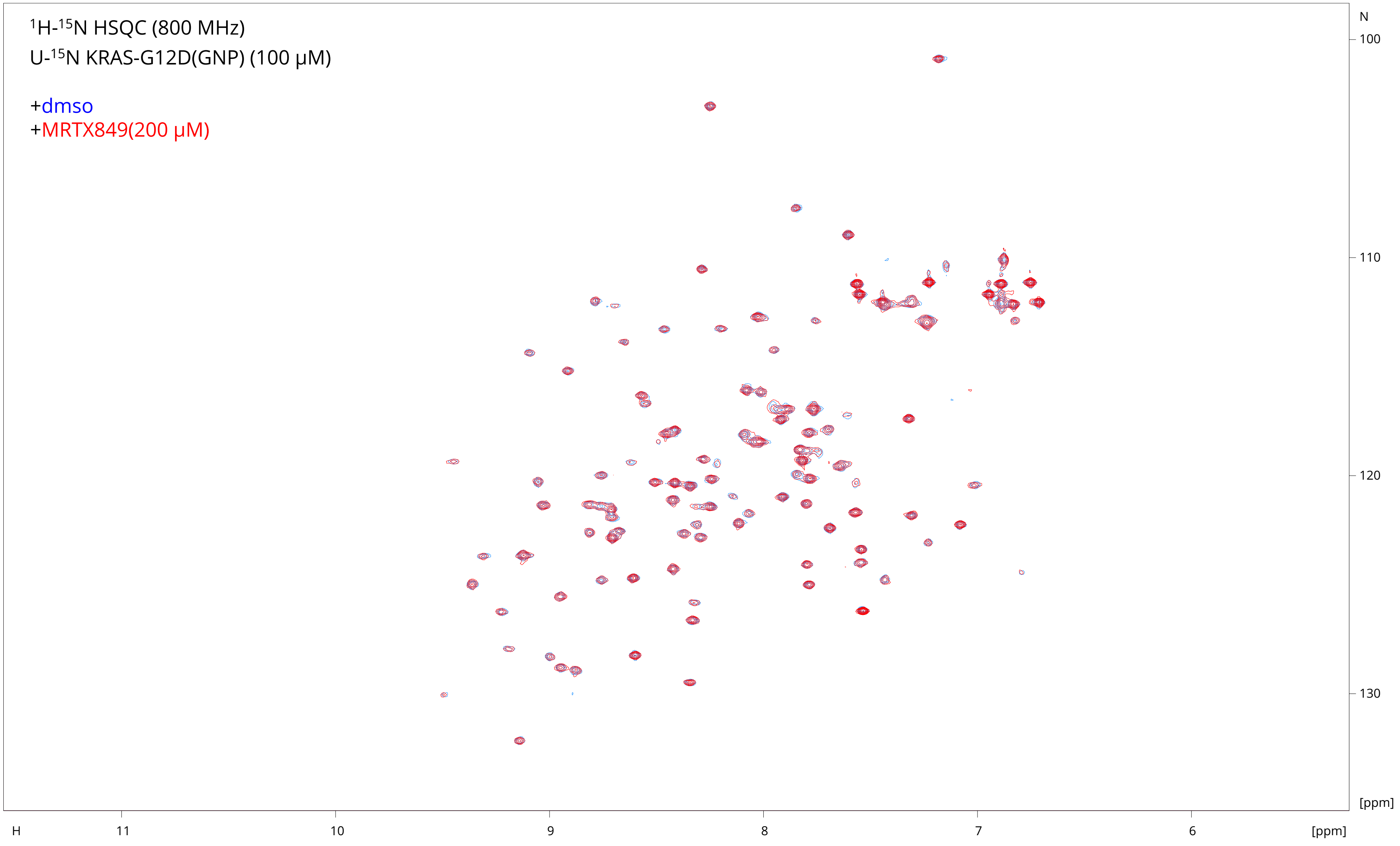

1H-15N HSQC (800 MHz)
U-15N KRAS-G12D(GNP) (100 μM)
+dmso
+MRTX849(200 μM)
N
100
110
120
130
[ppm]
H
11
10
9
8
7
6
[ppm]

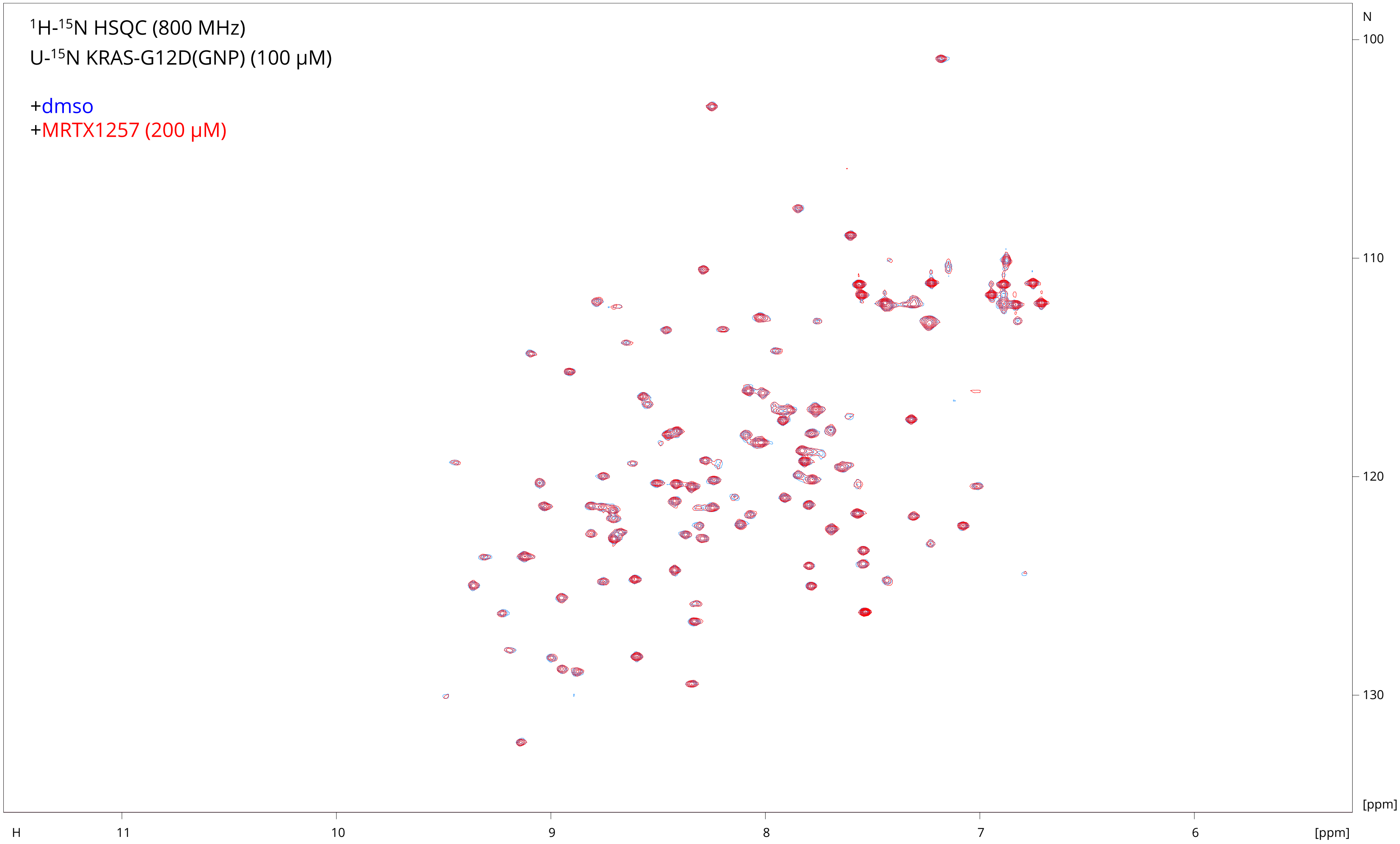

$^1$H-$^{15}$N HSQC (800 MHz)
U-$^{15}$N KRAS-G12D(GNP) (100 µM)
+dmso
+MRTX1257 (200 µM)
N
100
110
120
130
[ppm]
H
11
10
9
8
7
6
[ppm]

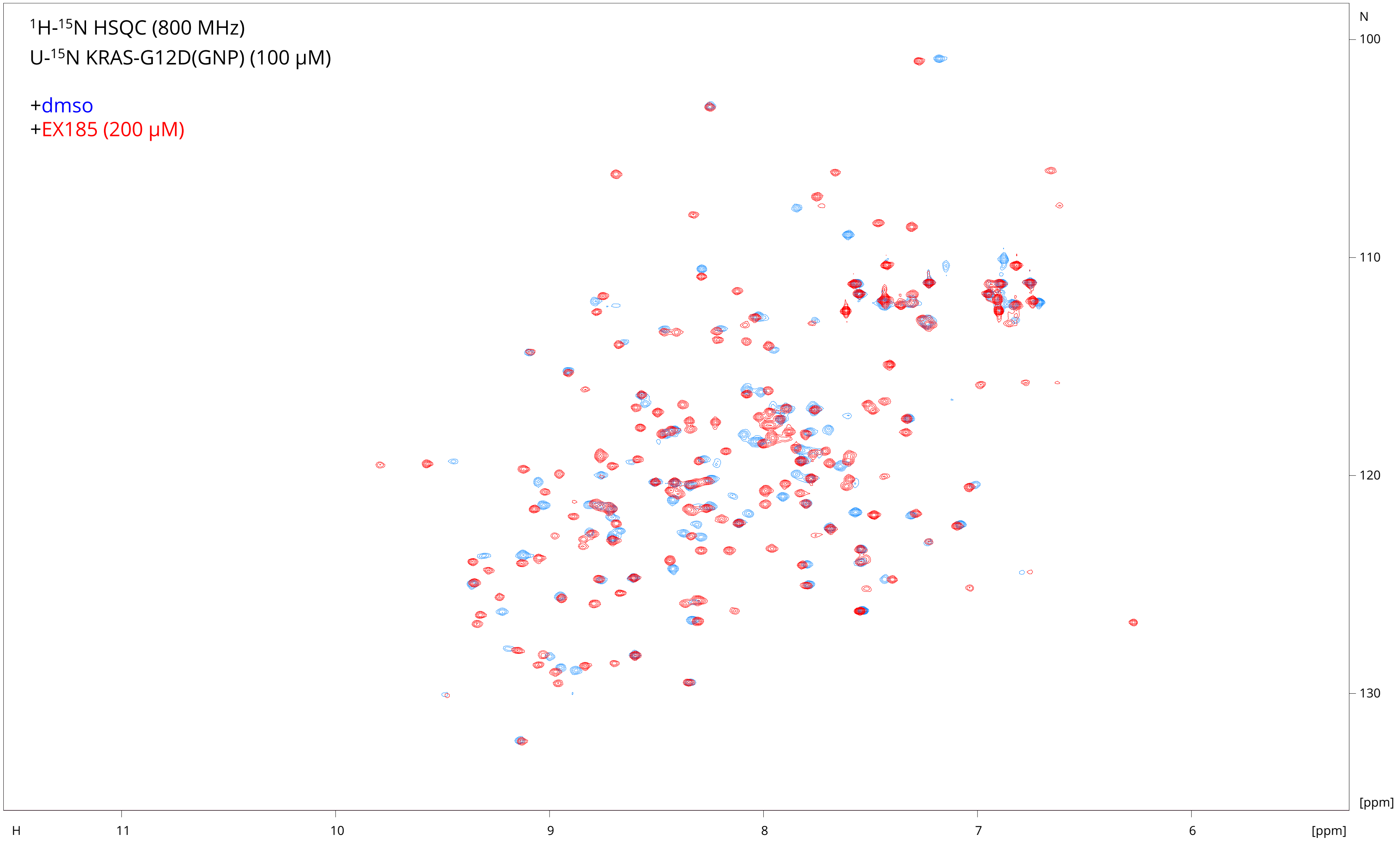

$^1$H-$^{15}$N HSQC (800 MHz)
U-$^{15}$N KRAS-G12D(GNP) (100 µM)
+dmso
+EX185 (200 µM)
N
100
110
120
130
[ppm]
H
11
10
9
8
7
6
[ppm]

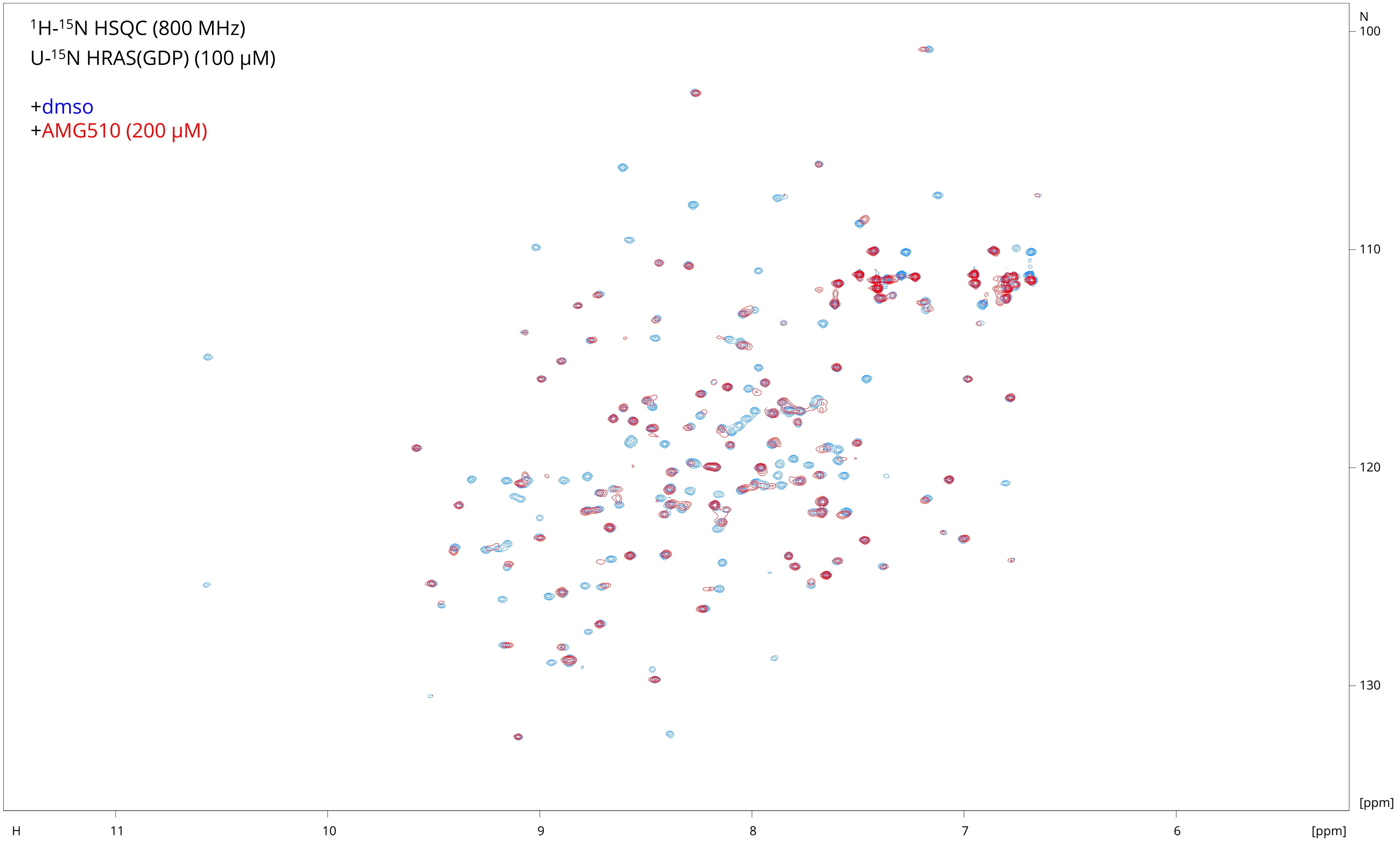
$^{1}H$-$^{15}N$ HSQC (800 MHz)
U-$^{15}N$ HRAS(GDP) (100 μM)
+dmso
+AMG510 (200 μM)
N
100
110
120
130
[ppm]
H
11
10
9
8
7
6
[ppm]

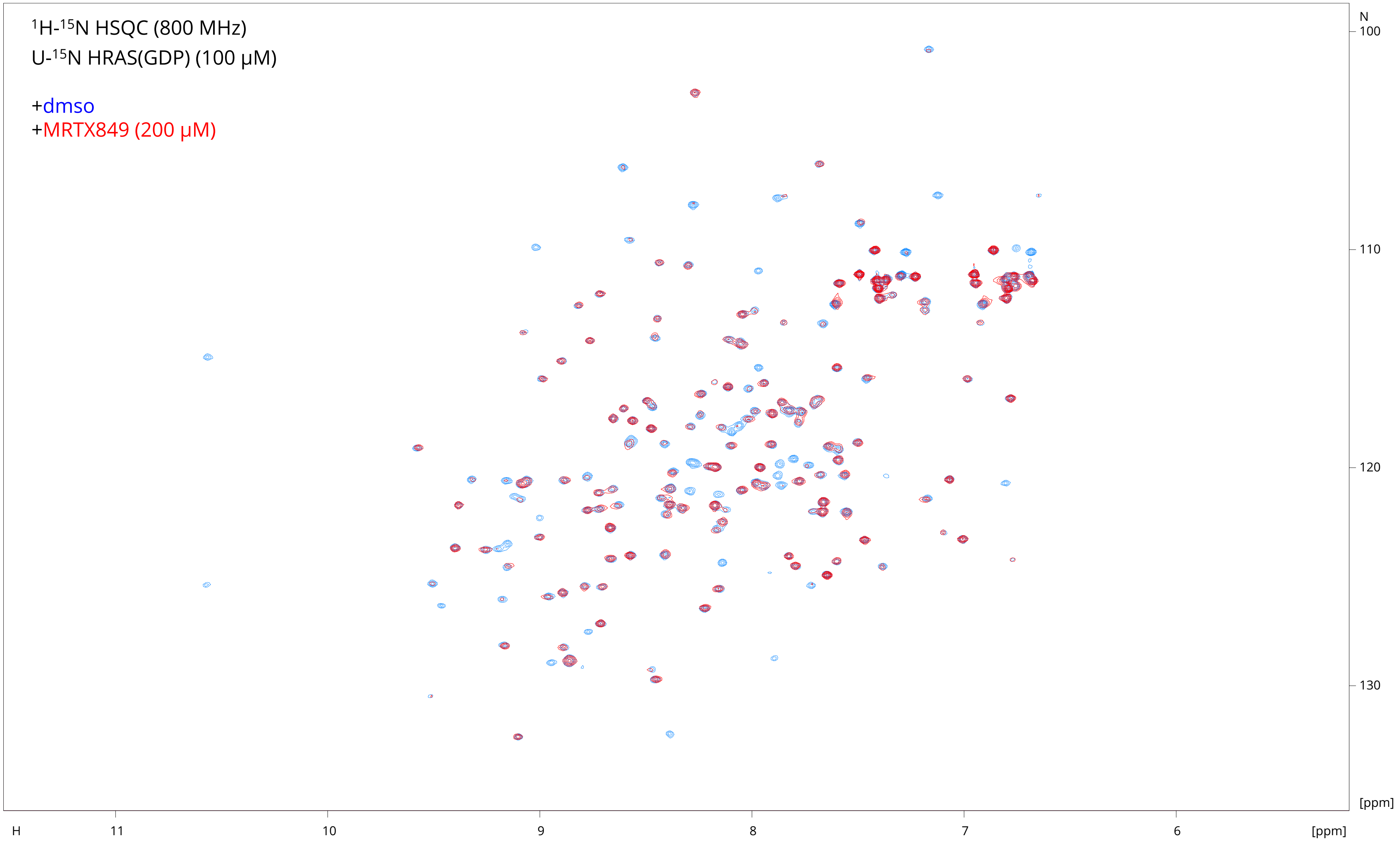

$^{1}H$-$^{15}N$ HSQC (800 MHz)
U-$^{15}N$ HRAS(GDP) (100 μM)
+dmso
+MRTX849 (200 μM)
N
100
110
120
130
[ppm]
H
11
10
9
8
7
6
[ppm]

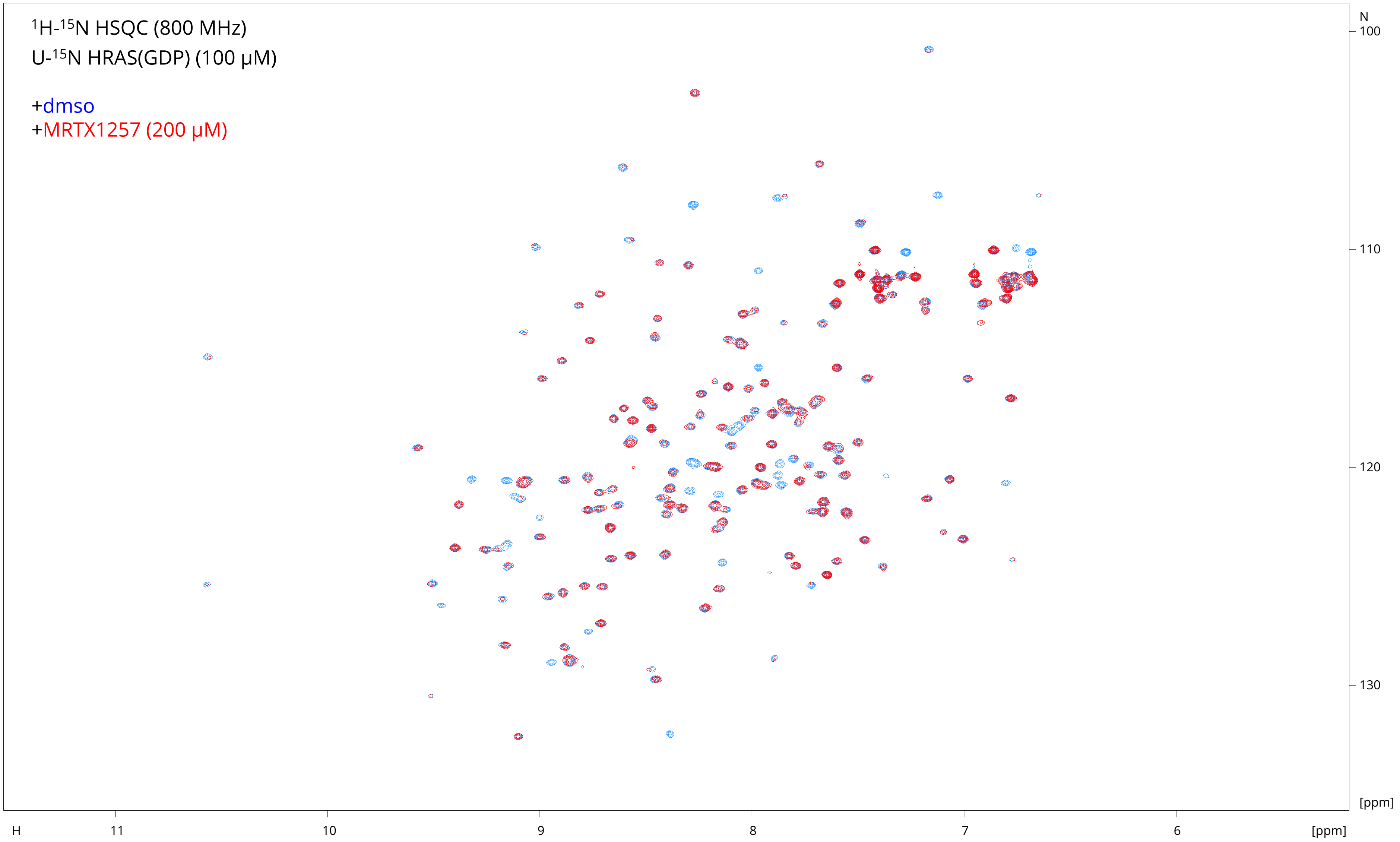
$^{1}$H-$^{15}$N HSQC (800 MHz)
U-$^{15}$N HRAS(GDP) (100 μM)
+dmso
+MRTX1257 (200 μM)
N
100
110
120
130
[ppm]
H
11
10
9
8
7
6
[ppm]

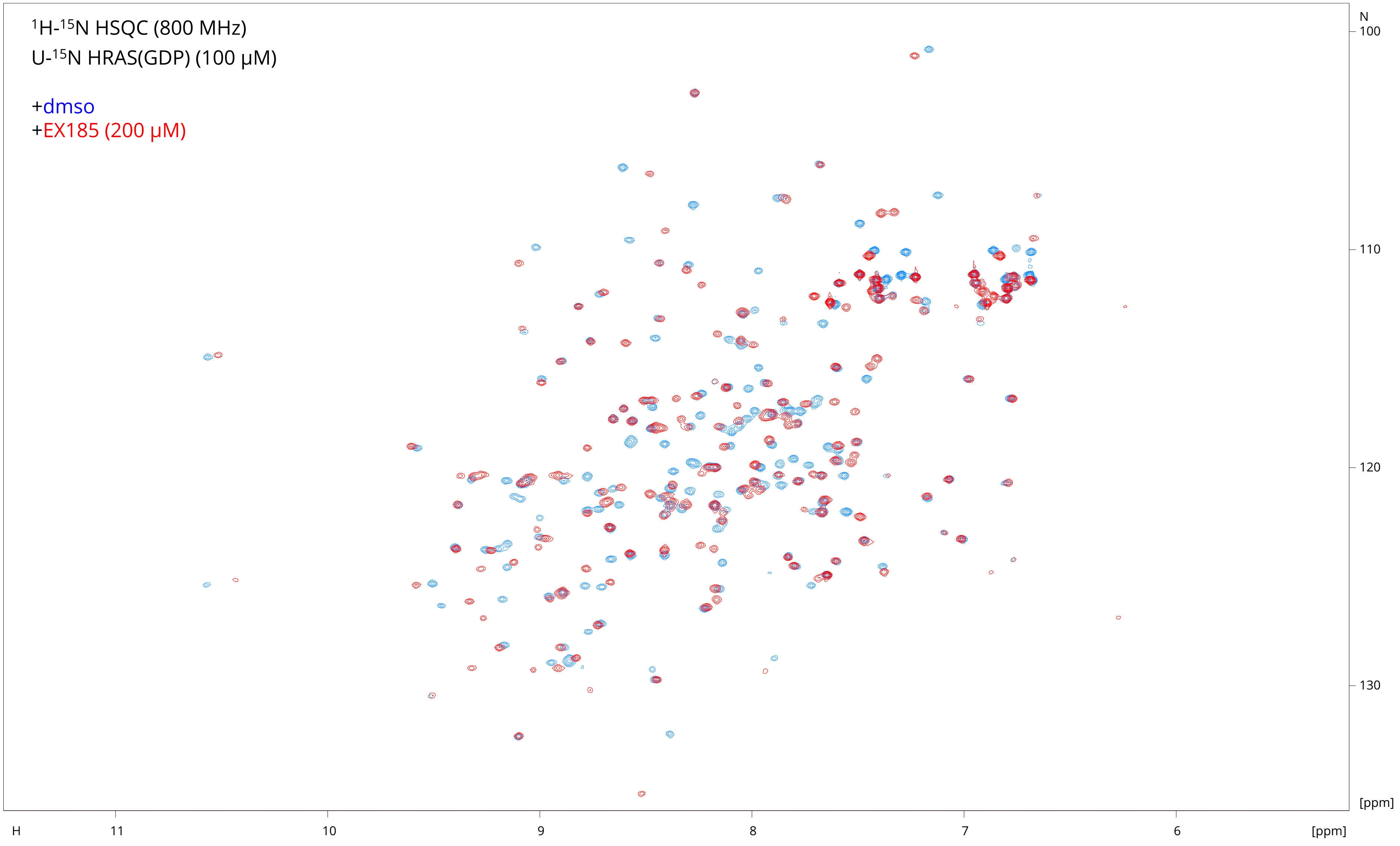

$^{1}H$-$^{15}N$ HSQC (800 MHz)
U-$^{15}N$ HRAS(GDP) (100 µM)
+dmso
+EX185 (200 µM)
N
100
110
120
130
[ppm]
H
11
10
9
8
7
6
[ppm]

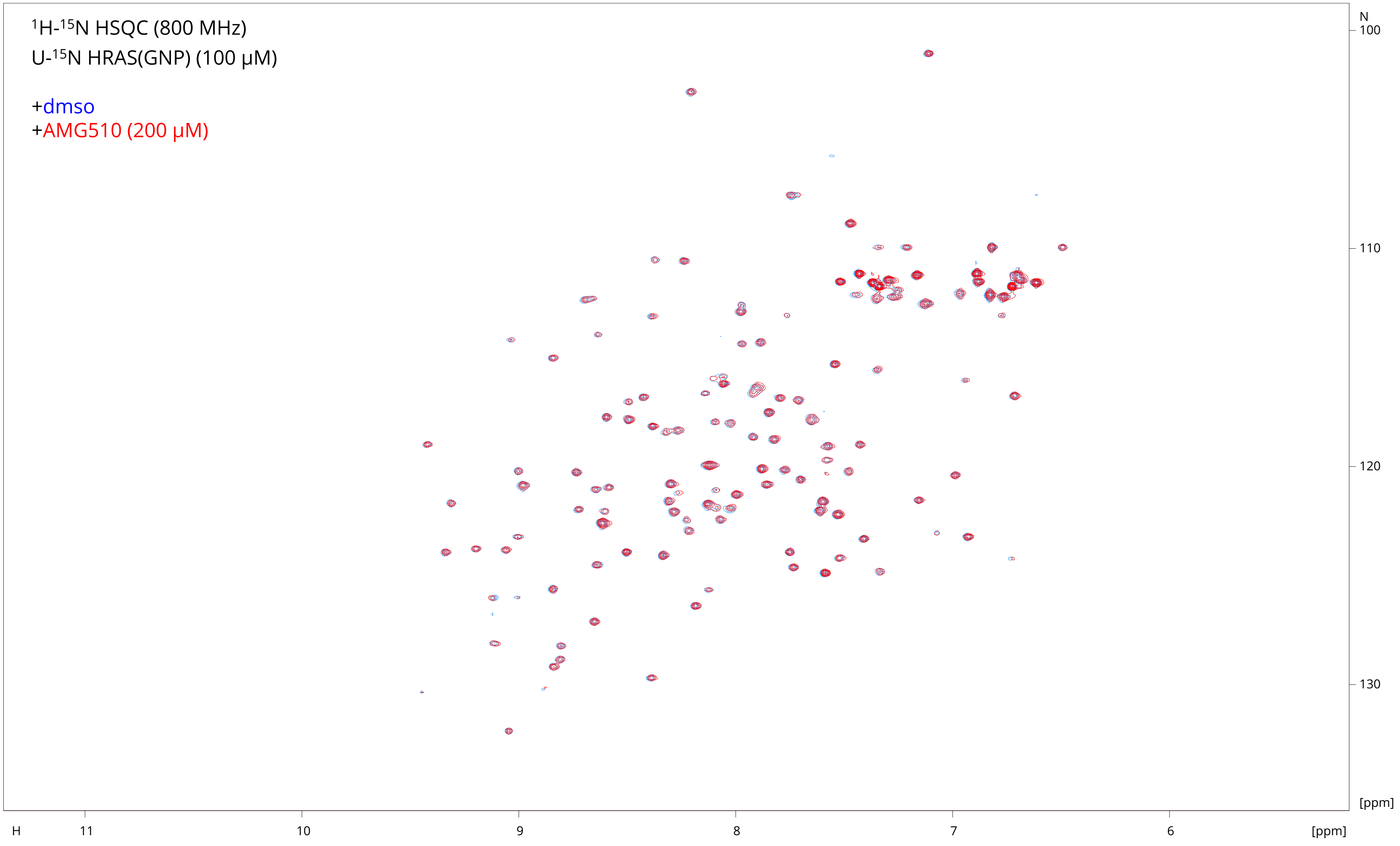

$^{1}$H-$^{15}$N HSQC (800 MHz)
U-$^{15}$N HRAS(GNP) (100 µM)
+dmso
+AMG510 (200 µM)
N
100
110
120
130
[ppm]
H
11
10
9
8
7
6
[ppm]

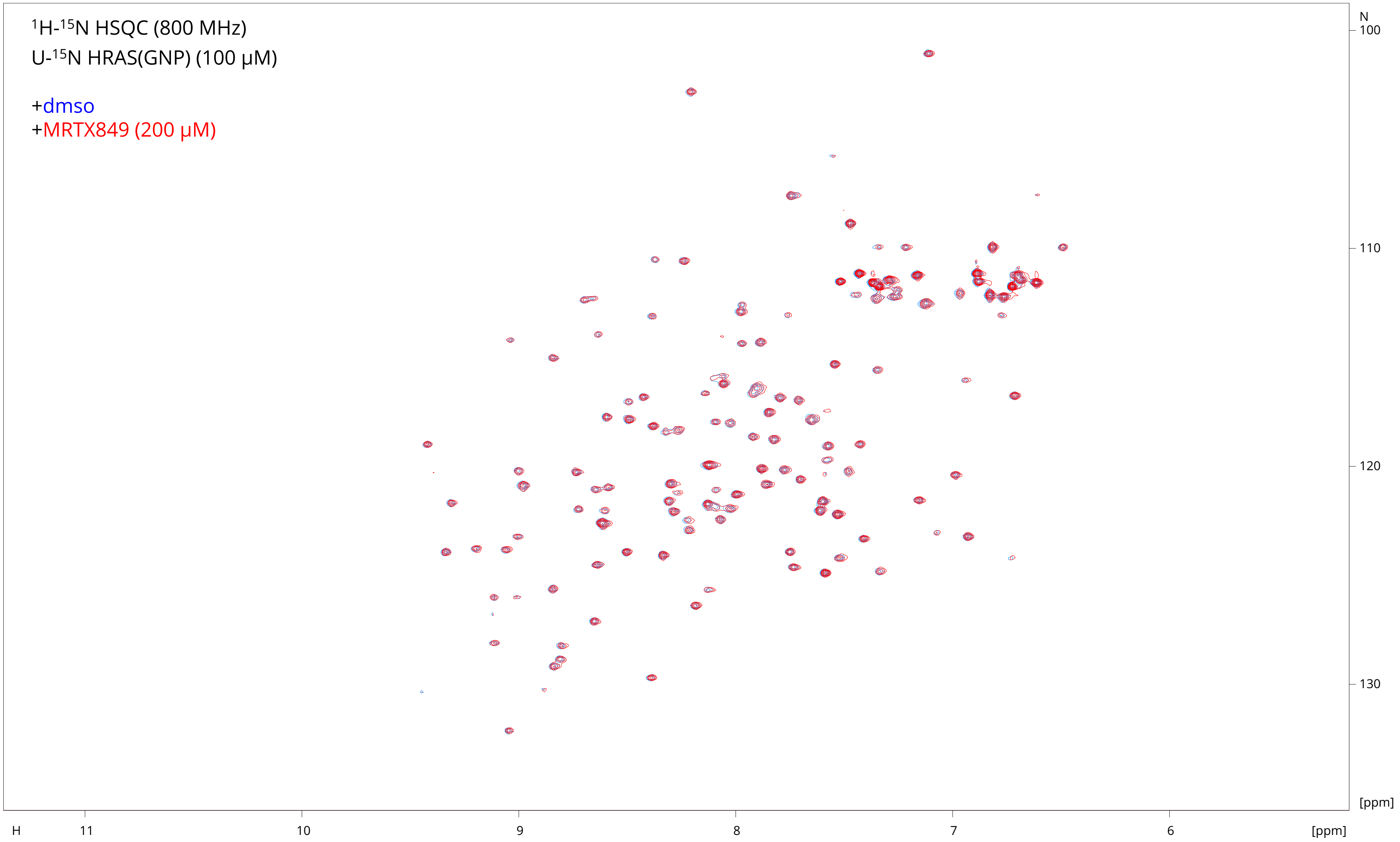

1H-15N HSQC (800 MHz)
U-15N HRAS(GNP) (100 μM)
+dmso
+MRTX849 (200 μM)
N
100
110
120
130
[ppm]
H
11
10
9
8
7
6
[ppm]

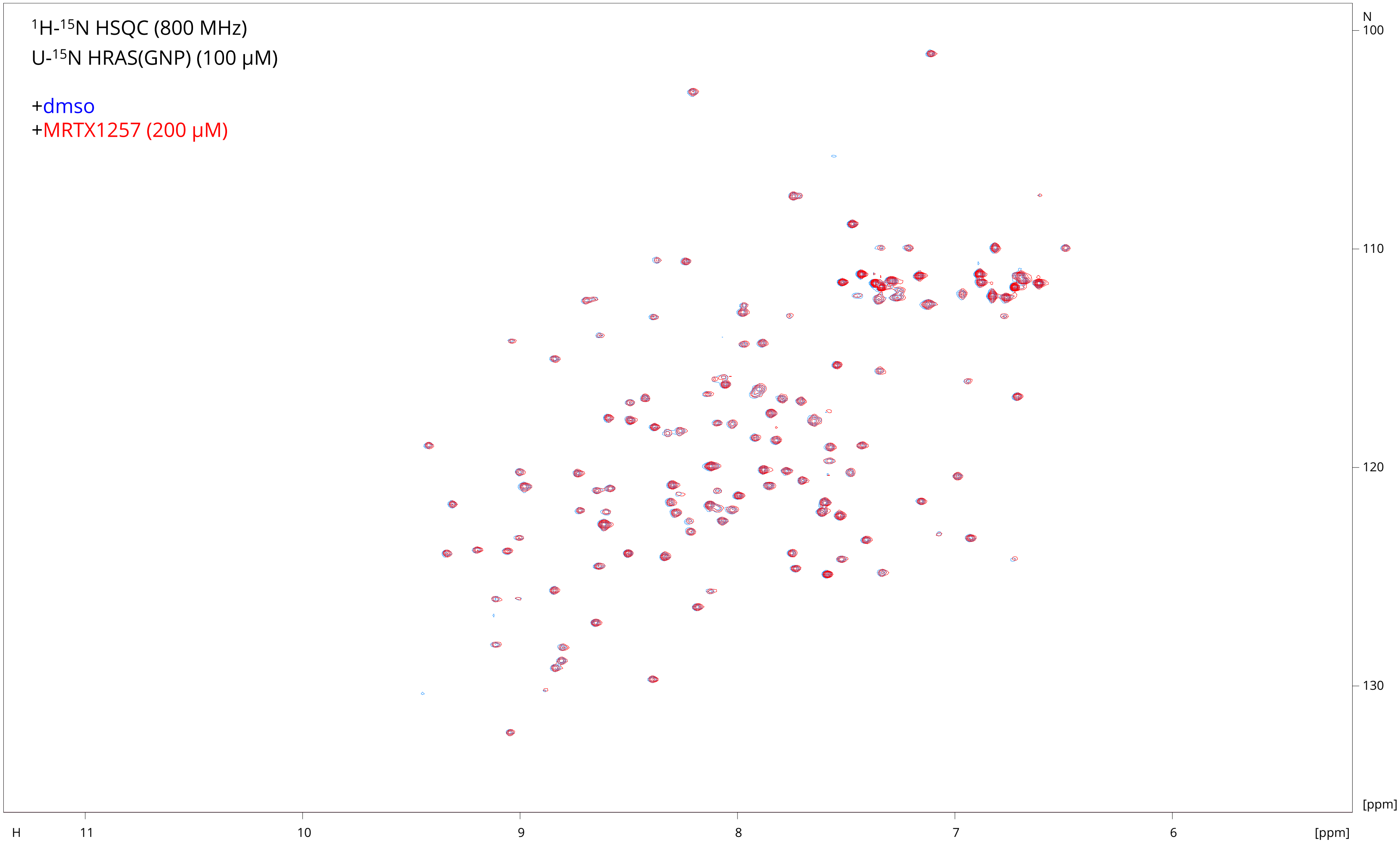

$^{1}H$-$^{15}N$ HSQC (800 MHz)
U-$^{15}N$ HRAS(GNP) (100 μM)
+dmso
+MRTX1257 (200 μM)
N
100
110
120
130
[ppm]
H
11
10
9
8
7
6
[ppm]

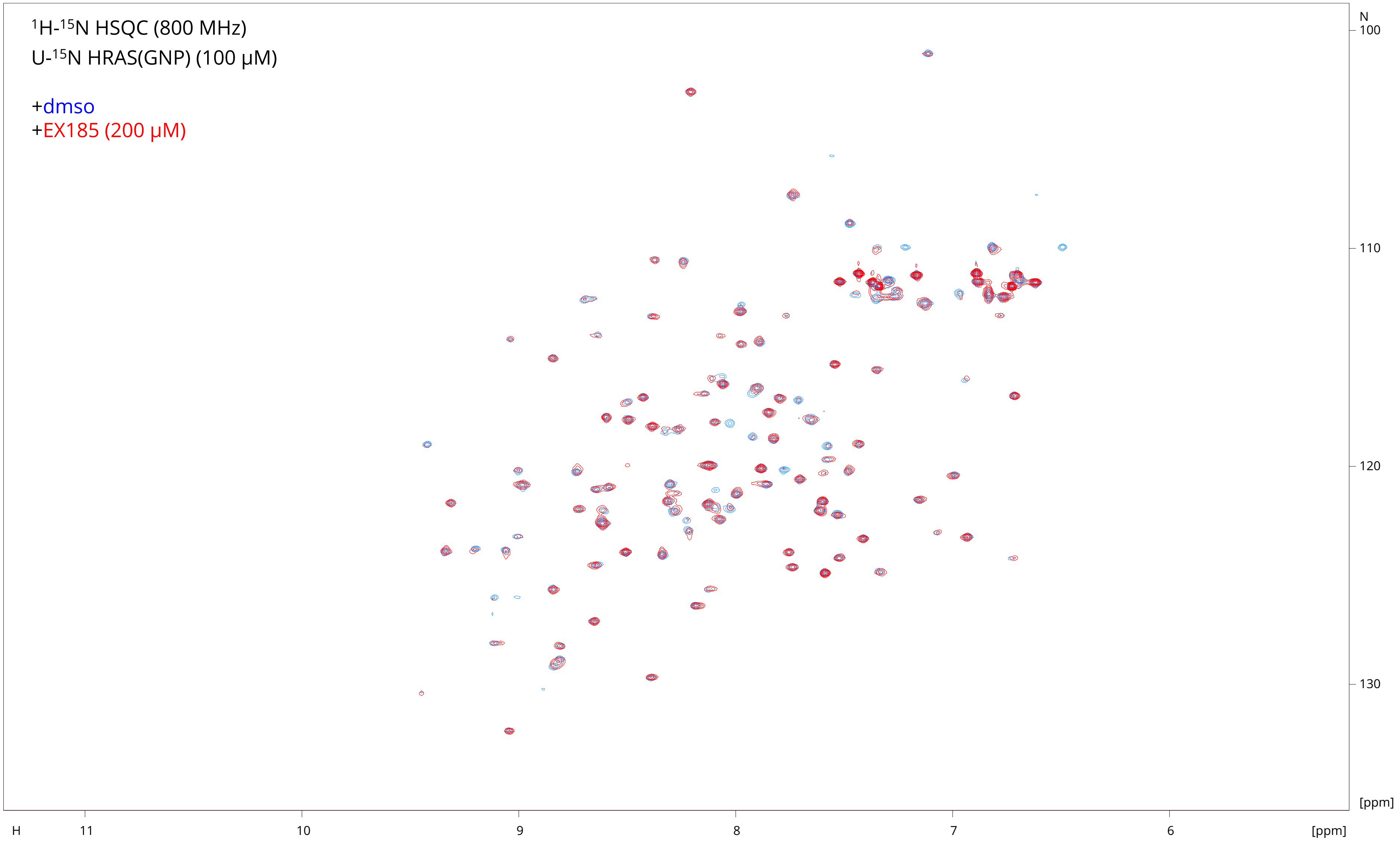

$^{1}H$-$^{15}N$ HSQC (800 MHz)
U-$^{15}N$ HRAS(GNP) (100 μM)
+dmso
+EX185 (200 μM)
N
100
110
120
130
[ppm]
H
11
10
9
8
7
6
[ppm]
