## Supplementary material for "KRAS is vulnerable to reversible switch-II pocket engagement in cells": Methods

**CONTACT FOR REAGENT AND RESOURCE SHARING;**

Further information and requests for resources and reagents should be directed to and will be fulfilled by the Lead Contact Matthew B. Robers.

**EXPERIMENTAL MODELS AND SUBJECT DETAILS**

HEK-293 cells (ATCC), A-375 cells (ATCC), HCT-116 cells (ATCC), NCI-H358 cells (ATCC), NCI-H647 cells (ATCC), Mia PaCa-2 Cells (ATCC), and SW-1990 cells (ATCC) were cultured in DMEM (Gibco) + 10% FBS (Seradigm), with incubation in a humidified, 37°C/5% CO_2_ incubator. H1975 cells (ATCC) were cultured in RPMI 1640 (GIBCO) + 10% FBS, with incubation in a humidified, 37°C/5% CO_2_ incubator.

**METHOD DETAILS**

*Cell Transfections and BRET measurements*

For cellular BRET measurements, a luciferase donor signal was produced at multimeric RAS using the NanoBiT approach. N-terminal LgBiT- or SmBiT-RAS fusions were encoded in pNB3K and pNB4K (respectively) expression vectors (Promega), including flexible 15 residue linkers (GSSGGGGSGGGGSSG) between the tag and each RAS isoform. All RAS ORFs were full length unless otherwise noted. HEK-293 cells were transfected with SmBiT-RAS and LgBiT-RAS fusion constructs using FuGENE HD (Promega) according to the manufacturer’s protocol. Briefly, SmBiT-RAS and LgBiT-RAS constructs were diluted together into Transfection Carrier DNA (Promega) at a mass ratio of 1:1:8 (mass/mass), after which FuGENE HD was added at a ratio of 1:3 (μg DNA: µL FuGENE HD). 1 part (vol) of FuGENE HD complexes thus formed were combined with 20 parts (vol) of HEK-293 cells suspended at a density of 2 x 10^5^ per mL in Opti-MEM containing 1% (v/v) FBS, followed by incubation in a humidified, 37°C/5% CO_2_ incubator for 18−24 hr. Following transfection, cells were washed with PBS, harvested by trypsinization, and resuspended in Opti-MEM containing 1% (v/v) FBS. BRET assays were performed in white, 96-well Non-Binding Surface plates (Corning) at a density of 2 x 10^4^ cells/well. All chemical inhibitors were prepared as concentrated stock solutions in DMSO (Sigma-Aldrich) and diluted in Opti-MEM (unless otherwise noted) to prepare working stocks. Cells were equilibrated with energy transfer probes and test compound prior to BRET measurements, with an equilibration time of 2 hours unless otherwise noted. RAS tracer was prepared first at a stock concentration of 100X in DMSO, after which the 100X stock was diluted to a working concentration of 20X in tracer dilution buffer (12.5 mM HEPES, 31.25% PEG-400, pH 7.5). For tracer dose response measurements, the RAS tracer was added to the cells in an 8 point, 3-fold dilution series starting at a final concentration of 2µM. For target engagement analysis, the RAS tracer was added to the cells at a final concentration of 1µM. To measure BRET, NanoBRET NanoGlo Substrate (Promega) was added according to the manufacturer’s recommended protocol, and filtered luminescence was measured on a GloMax Discover luminometer equipped with 450 nm BP filter (donor) and 600 nm LP filter (acceptor), using 0.5 s integration time. Unlike the full length NanoLuc protein, the signal of which can be quenched in extracellular environments using an impermeable inhibitor of NanoLuc, the NanoBiT luciferase in its current form is not amenable to extracellular quenching using the same approach. Raw BRET ratios were calculated by dividing the acceptor counts by the donor counts. Milli-BRET units (mBU) were calculated by multiplying the raw BRET values by 1000. When normalized BRET was used, mBRET values were normalized using Equation 1;

*Normalized BRET (%) = [(A-C)/(B-C)]*100*

Where A = mBRET in the presence of test compound and tracer, B = mBRET in the presence of vehicle and tracer, and C = mBRET in the presence of a saturating 20µM dose of BI-2852. Apparent tracer affinity values (EC_50_) were determined using the sigmoidal dose-response (variable slope) equation available in GraphPad Prism (Equation 2);

*Y = Bottom + (Top-Bottom)/(1+10^((LogEC50-X)*HillSlope)).*

In some cases, the RAS tracer was not saturable up to the solubility limit of the tracer, so the EC_50_ of the tracer is reported as > 1µM. For determination of test compound potency, competitive displacement data were plotted with GraphPad Prism software and data were fit to Equation 1 to determine the IC_50_ value. For fractional occupancy determination, the following equation (Equation 3) was used;

*% Occupancy = [1 – (X – Z)/( Y – Z)]*100*

where X = mBRET in the presence of the test compound and tracer, Y = mBRET in the presence of vehicle and tracer, and Z = BRET in the presence of a saturating 20µM dose BI-2852.

*Measurements of Total LgBiT-RAS levels*

Total LgBiT expression level was determined in the presence of a saturating concentration (100nM) of high affinity HiBiT peptide (Peptide 2.0), 1X NanoBRET Target Engagement substrate, and 50 mg/mL digitonin (as a permeabilization agent). Unfiltered luminescence was measured using the Nano-Glo protocol on a Glomax Discover luminometer with a 0.3 s integration time.

*Measurements of PPI disruption for KRAS(G12D):CRAF(RBD) in cells*

For the KRAS(G12D):CRAF Ras binding domain (RBD) NanoBiT interaction assay, CMV-based expression constructs were made encoding fusions of LgBiT to KRAS 4B (UNIPROT P01116-2) with the G12D mutation and SmBiT to residues 51-133 of CRAF [UNIPROT P04049-1, CRAF(RBD)). HEK293 cells (~4E6) were transiently transfected in T75 flasks with plasmids encoding a LgBiT-KRAS(G12D) and SmBiT-CRAF(RBD). Plasmids were transfected at 500 ng/construct/flask together with 9 µg of Transfection Carrier DNA (Promega) at a 3:1 lipid:DNA ratio using FuGENE HD (10 ml total volume). Following expression for 24 hours, cells were plated at 20,000 cells/well in Opti-MEM I (Thermo) containing 4% FBS and allowed to attach overnight. Serial dilutions of EX185 were made in Opti-MEM I containing 4% FBS and 1X Vivazine substrate (Promega N2581) to generate 1X solutions containing varying concentrations of EX185. Existing medium was removed by plate inversion and blotting, and 1X solutions were added to respective wells. Luminescence was measured every 5 minutes in a GloMax Discover luminometer at 37°C for 16 h using a 1 s integration time.

*Measurements of anti-proliferative activity in cells*

Anti-proliferative activity of test compounds towards human cell lines was measured as a decrease in cellular ATP levels using the CellTiter-Glo 2.0 assay (Promega) according to the manufacturer’s protocols. Briefly, cells grown to confluence in cell culture medium were harvested by trypsinization and seeded at 2,500 (HEK-293) or 5,000 (other cell types) cells per well into 96-well tissue culture treated assay plates (Corning 3917). The cells were incubated in a humidified, 37°C/5% CO_2_ incubator and allowed to adhere. After 24 hours, test compounds prepared as 10X stock solutions in culture medium were added to the cells, and the cells were incubated in a humidified, 37°C/5% CO_2_ incubator for another 48 hours. To measure total ATP levels, 2X CellTiter-Glo 2.0 (Promega) reagent was added to each well. The plates were mixed on an orbital shaker at 600 rpm for 2 min, incubated at ambient temperature for another 8 min, and then the total unfiltered luminescence was measured using the CellTiter-Glo protocol on the Glomax Discover (integration 0.3 s). Luminescence values were normalized to that of the vehicle treated controls on each plate. Vehicle normalized luminescence values were plotted as a function of compound dose. Clear monophasic behavior was fitted to Equation 1 to interpolate the anti-proliferative potency (IC_50_). To fit the biphasic antiproliferative behavior of MRTX849 in H358 and Miapaca-2 cells, the data were fit to a biphasic inhibitor model with variable hill slope (Equation 4) below

*Y = Bottom + (Top-Bottom)*Frac/(1+(IC50_1/X)^nH1) + (Top-Bottom)* (1-Frac)/(1+(IC50_2/X)^nH2)*

*Inhibition of ERK Phosphorylation*

Cellular ERK phosphorylation (Thr202/Tyr204) was quantified using Phospho-ERK (Thr202/Tyr204) cellular kit (Cisbio). SW-1990 cells (1x10^6^/mL, 50 µL/well) were plated in cell culture medium (DMEM (Gibco) + 10% FBS (Seradigm)) 12 h before the experiments. On the day of treatment, serially diluted 2X solutions were prepared in cell culture medium (9+1 points, 3:1 dilution starting from 60 µM, DMSO 0.6%). Cells in each well were treated with 2X small molecule solutions (50 µL), after which they were incubated for 4 h at ambient temperature. Media were removed by aspiration. Cells were lysed with 50 µL supplemented lysis buffer 1X (Cisbio) for 30 min at ambient temperature. Lysates were homogenized and transferred (16 µL) to a low volume 384-well detection plate (Corning 4513). Pre-mixed phospho ERK antibody solutions (4 µL, Advanced p-ERK1/2 d2 Ab 20X (19X) + Advanced p-ERK1/2 Eu Cryptate Ab 20X (19X) + detection buffer, Cisbio) were added to each well of the plates. The mixtures were incubated at ambient temperature for 4 h before reading on a TECAN plate reader using the TR-FRET mode with 60 µs Lag time and 500 µs Integration time. The ratio of the acceptor and donor emission signals for each individual well was calculated by

TR FRET Ratio = [Signal 665 nm]/[Signal 620 nm]*10,000

Total ERK data was acquired via the same procedure using Total ERK cellular kit (Cisbio) on the same cell lysates. For each marker (p-ERK or t-ERK), TR FRET Ratio was normalized to the respective DMSO control. The ratio of phosphoERK over total ERK was calculated and fit to Equation 4 above.

*Preparation of U-^15^N Ras proteins.*

The plasmids for bacterial expression of HRAS 1-166 (WT; His-TEV-N; pProEx; ampicillin resistance) and KRAS 1-169 (WT, G12C, and G12D; His-TEV-N; pJ411; kanamycin resistance) have been previously published.^1^ BL21(DE3) competent cells were transformed with 1-2 ng of plasmid, plated on LB agar containing the appropriate antibiotic (carbenicillin or kanamycin), and allowed to grow at 37 °C. Colonies were picked, and a small starter culture (100-200 ml) in M9 minimal media containing 1 g/L ^15^N ammonium chloride (99%, Cambridge Isotope Laboratories) and the appropriate antibiotic (0.1 mg/ml carbenicillin or 0.1 mM kanamycin) was grown at 37 °C with shaking at 200 rpm overnight. The starter culture was divided into shaker flasks each containing 1 L of the same medium (25-40 ml per flask, 2-4 flasks) and continued growing at 37 °C with shaking at 200 rpm. Protein expression was induced after cooling the cultures to 18 °C (A600 0.4 to 0.6) by adding 1 ml 1.0 M IPTG, and the flasks were shaken at 200 rpm overnight. Procedures for lysis and purification were followed as previously published.^1^ Nucleotide exchange from GDP to GPPNHP (Jena Biosciences) was performed prior to the final gel filtration purification and according to a published procedure comprising EDTA-mediated exchange, desalting, and cleavage of residual GDP with an alkaline phosphatase (CIP or Quick CIP, NEB).^1-4^ Final gel filtration purification was performed on a Superdex 75 column (GE) with storage buffer. Proteins were concentrated to 0.5 - 1 mM (Amicon Ultra-4, 10k MWCO, EMD), concentrations were determined by uv absorbance (ε = 13410 M^-1^cm^-1^ for HRAS 1-166 and 11920 M^-1^cm^-1^ for KRAS 1-169), and aliquots were flash frozen with liquid N_2_ and stored at -80 °C.

Storage buffer: 40 mM HEPES, 150 mM NaCl, 4 mM MgCl_2_, 5% glycerol, 7% D_2_O. Titrated to pH 7.4 with NaOH.

*Preparation of ^15^N-labelled KRAS(G12C)-MRTX849.*

A 4 ml 0.10 mM sample of U-^15^N KRAS(G12C)-GDP 1-169 (*M* = 19579) was reserved prior to the final gel filtration purification step. The concentration was determined by a bicinchoninic acid assay (Pierce, Thermo Fisher Scientific) relative to a bovine serum albumin standard. A solution of MRTX849 (120 µl, 10 mM in dmso, 3.0 equiv) was added, and the mixture was rotated at ambient temperature for 15 minutes then concentrated and purified by gel filtration as described above. The purified protein was analyzed by LC/MS to ensure complete conversion to the 1:1 protein-ligand adduct (*M* = 20182).

*^1^H-^15^N HSQC NMR sample preparation and acquisition.*

A 0.030 µmol sample of U-^15^N protein in storage buffer was diluted to 270 µl with HSQC NMR sample buffer on ice. 30 µl of a dmso-*d*_6_ solution containing the small molecule ligand was added, the sample was mixed by vortex, and the resulting solution was transferred to a 5 mm D_2_O-matched Shigemi NMR tube (BMS-3). The final concentration of protein and ligand were 100 and 200 µM, respectively. 1D ^1^H (ZGESGP) and 2D ^1^H-^15^N fast HSQC (FHSQCCF3GPPH, ns=8, tdf1=256, GARP decoupling) spectra were recorded on an 800 MHz Bruker Avance spectrometer at 298 K. ^1^H chemical shifts were corrected with an internal standard (1 mM DSS at 0 ppm). For any case in which strong chemical shift perturbations were observed, a second sample containing the same protein-ligand combination at 50 and 100 µM, respectively was also prepared and spectra acquired under the same conditions. For the mixed samples containing both nucleotide states, the scan number was doubled (ns=16) to improve SNR.

HSQC NMR buffer: 40 mM HEPES, 150 mM NaCl, 4 mM MgCl_2_, 7% D_2_O. Titrated to pH 7.4 with NaOH.

Spectra were also acquired with 0, 5, and 10% dmso-*d*_6_ and/or with minor adjustments to pH and temperature for comparison to previously published assignments; well-resolved peaks were assigned by comparison to data imported from the BMRB.

HRAS-GDP 1-166: BMRB entry 18479.^5^

HRAS-GPPNHP 1-166: BMRB entry 17678.^5^

KRAS-GDP 1-169: BMRB entry 27720.^6^

KRAS(G12C)-GDP 1-169: BMRB entry 27646.^6^

KRAS(G12D)-GDP 1-169: BMRB entry 27719.^6^

Spectra were analyzed with Bruker Topspin 4.0, CCPNMR Analysis v3,^7^ and/or MestReNova v14. The spectra images were created with CCPNMR Analysis v3. Full spectra of each protein-ligand combination (red) superimposed with the dmso-*d*_6_ control (blue) are shown in a supplementary note.

*^1^H-^15^N-^1^H NOESY-HSQC NMR sample preparation and acquisition.*

A 0.150 µmol sample of U-^15^N protein in storage buffer was diluted to 400 µl with NOESY NMR buffer on ice. The buffer was exchanged to the NOESY NMR buffer with a desalting column (5 ml HiTrap Desalting, Cytiva and AKTA FPLC, GE). The protein-containing fractions were combined (1.5 ml), concentrated to 0.3 ml (10k MWCO Amicon Ultra-4, EMD), transferred to a 5 mm Shigemi NMR tube (BMS-3), and gently sparged with Ar before sealing with parafilm (final concentration ~0.5 mM). 1D ^1^H (ZGESGP), 2D ^1^H-^15^N fast HSQC (FHSQCCF3GPPH, ns=8, tdf1=256, GARP decoupling), and 3D ^1^H-^15^N-^1^H NOESY-HSQC (NOESYHSQCF3GPWG3D, ns=16, tdf1=128, tdf2=40, GARP decoupling) NMR spectra were recorded on an 800 MHz Bruker Avance spectrometer at 298 K. A second 2D HSQC spectrum was acquired after the 3D NOESY-HSQC experiment with identical parameters to confirm sample stability during the 28 h acquisition. ^1^H chemical shifts were corrected with an internal standard (1 mM DSS at 0 ppm).

NOESY NMR buffer: 20 mM sodium phosphate, 140 mM NaCl, 15 mM MgCl_2_, 10 mM EDTA, 3 mM NaN_3_, 1 mM GDP, 1 mM DSS, 10% D_2_O. Titrated to pH 7.4 with NaOH.

3D NOESY-HSQC data were analyzed with CCPNMR Analysis v3. Sequential backbone NH ^1^H and ^15^N shifts were identified by mutual NOESY crosspeaks, and the 2D ^1^H-^15^N HSQC spectrum was assigned from this data and comparison to the published shifts for KRAS(G12C)-GDP (BMRB 27646).

*Chemical Synthesis*

General information. Solvents were purchased from Sigma Aldrich or Fisher Scientific. Solvents referred to as “dry” were either collected from a solvent system containing mol sieves or purchased anhydrous from suppliers. NanoBRET 590 SE was obtained from Promega Corp. Madison, WI. Flash column chromatography was performed with a Combiflash Rf+ (Teledyne Isco) on silica. Preparative HPLC was performed with a 30x250 mm 5 µM C18 column (Waters Xbridge). High resolution ESI MS data was obtained from LC/MS with a C18 column (Waters Acquity UPLC and Xevo G2-XS QTof). ^1^H and ^19^F NMR spectra were recorded on a Bruker Avance 400 MHz spectrometer or a Bruker Ascend 400 MHz spectrometer. Chemical shifts (δ) are quoted in parts per million (ppm) and referenced to the residual solvent peak. Multiplicities are denoted as s-singlet, d-doublet, t-triplet, q-quartet and quin-quintet and derivatives thereof (br denotes a broad resonance peak). Coupling constants are given in Hz and round to the nearest 0.1 Hz.

*Synthesis of pan RAS Tracer*

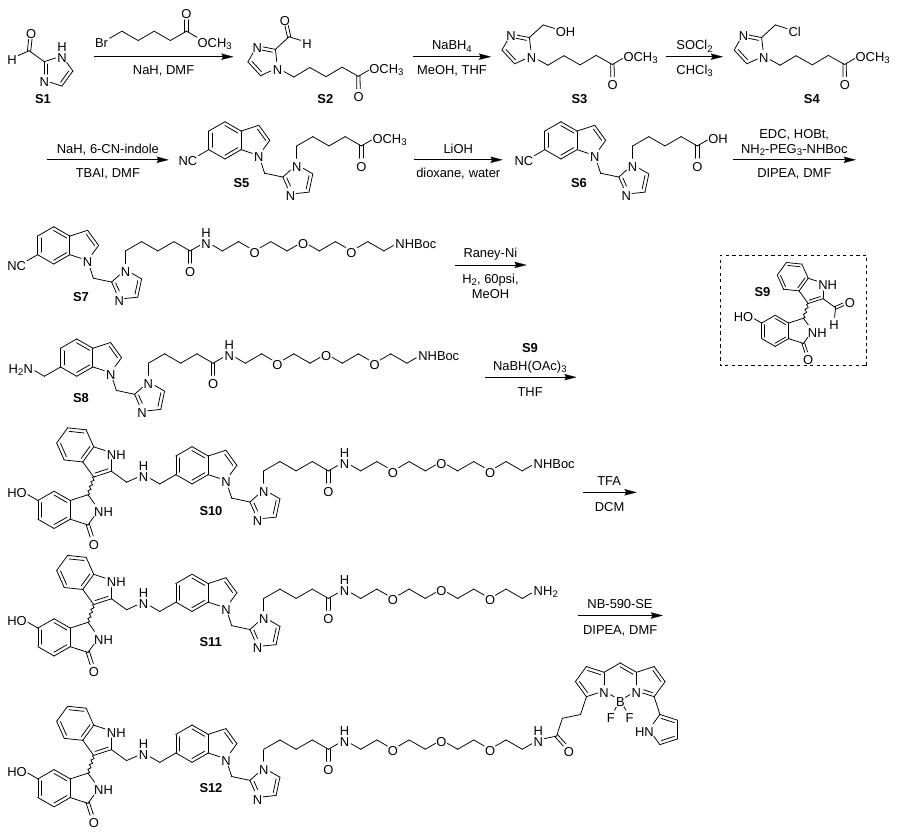

Supplementary Scheme 1. Synthesis of pan RAS tracer S12.

**Methyl 5-(2-formyl-1H-imidazol-1-yl)pentanoate (S2).** To a suspension of 1H-imidazole-2-carbaldehyde (**S1**, 1.00 g, 10.4 mmol) in DMF (20 mL) chilled with an ice bath was added sodium hydride (60% oil dispersion, 500 mg, 12.5 mmol). The mixture was warmed to ambient temperature and methyl 5-bromopentanoate (2.44 g, 12.5 mmol) was added in DMF (5mL). The mixture stirred for 4h at ambient temperature. The reaction was quenched with water, diluted with brine, and extracted with chloroform/isopropanol (3:1). The organic layers were combined, dried with sodium sulfate, filtered, concentrated, and purified with silica gel chromatography to afford the desired product (0.92 g, 42%) as a colorless oil. ^1^H NMR (400 MHz, Methanol-d4) δ 9.73 (s, 1H), 7.53 (s, 1H), 7.28 (s, 1H), 7.10 (s, rotamer), 6.88 (s, rotamer), 4.46 (t, *J* = 7.2 Hz, 2H), 4.18 (t, rotamer), 3.66 (s, 3H), 2.38 (t, *J* = 7.2, 2H), 1.90 – 1.77 (m, 2H), 1.70 – 1.55 (m, 2H); ^13^C NMR (101 MHz, MeOD) δ 182.15, 175.30, 144.39, 131.84, 128.49, 126.99 (rotamer), 122.23 (rotamer), 52.04, 34.15 (rotamer), 33.98, 31.36, 31.27 (rotamer), 22.95 (rotamer), 22.66. ESI MS m/z 211 [M + 1]+.

**Methyl 5-(2-(hydroxymethyl)-1H-imidazol-1-yl)pentanoate (S3).** To a solution of methyl 5-(2-formyl-1H-imidazol-1-yl)pentanoate (0.92 g, 4.4 mmol) in methanol/tetrahydrofuran (1:1, 50 mL) chilled with an ice bath was added sodium borohydride (0.20 g, 5.3 mmol). The mixture stirred for 30 min. The reaction was quenched with HCl (3 mL, 2M) and then the pH was adjusted to 8. The mixture was concentrated with celite and purified with silica gel chromatography to afford the desired product (0.53 g, 57%) as a colorless oil. ^1^H NMR (400 MHz, Methanol-d4) δ 7.28 (s, 1H), 7.08 (s, 1H), 4.73 (s, 2H), 4.19 – 4.09 (m, 2H), 3.67 (s, 3H), 2.40 (td, *J* = 7.5, 2.2 Hz, 2H), 1.95 – 1.80 (m, 2H), 1.73 – 1.58 (m, 2H); ^13^C NMR (101 MHz, MeOD) δ 174.01, 146.67, 123.57, 121.14, 54.84, 50.68, 45.90, 32.66, 29.58, 21.49; HRMS (ESI+) calcd for C_10_H_16_N_2_O_3_ [M + H]+ m/z 213.1239, found 213.1223.

**Methyl 5-(2-(chloromethyl)-1H-imidazol-1-yl)pentanoate (S4) .** To a solution of methyl 5-(2-(hydroxymethyl)-1H-imidazol-1-yl)pentanoate (0.53 g, 2.5 mmol) in chloroform was added thionyl chloride (3.0 g, 25 mmol). The mixture stirred for 1 h at ambient temperature and heated to 75°C for 1 h. The reaction was concentrated, and the residue was suspended in ether. Filtration afforded the crude product (0.64 g) as a white solid. ESI MS m/z 231 [M + 1]+.

**Methyl 5-(2-((6-cyano-1H-indol-1-yl)methyl)-1H-imidazol-1-yl)pentanoate (S5).** To a solution of 1H-indole-6-carbonitrile (200 mg, 1.41 mmol) in DMF (20 mL) chilled with an ice bath was added sodium hydride (84 mg, 2.1 mmol, 60%), methyl 5-(2-(chloromethyl)-1H-imidazol-1-yl)pentanoate (0.38 g, 1.4 mmol), and tetrabutylammonium iodide (52 mg, 0.14 mmol). The mixture stirred at 0 °C for 15 min and ambient temperature for 2 h. The reaction was diluted with ethyl acetate and washed with water. The organic layers were combined, dried with sodium sulfate, filtered, concentrated, and purified with silica gel chromatography to afford the desired product (370 mg, 78%) as an orange oil. ^1^H NMR (400 MHz, Methanol-d4) δ 7.95 (s, 1H), 7.71 (d, *J* = 8.2 Hz, 1H), 7.52 – 7.46 (m, 1H), 7.33 (d, *J* = 8.0, 1H), 7.15 (s, 1H), 7.03 (s, 1H), 6.65 (s, 1H), 5.56 (s, 2H), 3.88 (t, *J* = 7.0 Hz, 2H), 3.61 (s, 3H), 2.06 (t, *J* = 7.0 Hz, 2H), 1.38 – 1.20 (m, 4H); ^13^C NMR (101 MHz, MeOD) δ 173.72, 142.41, 135.11, 132.19, 131.64, 126.70, 122.12, 121.61, 121.56, 119.99, 114.64, 103.76, 102.71, 50.65, 45.61, 42.16, 32.54, 29.58, 21.30; HRMS (ESI+) calcd for C_19_H_20_N_4_O_2_ [M + H]+ m/z 337.1664, found 337.1643; HPLC 97.8% (AUC at 286 nm) 3.10 min (Synergi Max-RP, water/ACN, 0.02%TFA).

**5-(2-((6-Cyano-1H-indol-1-yl)methyl)-1H-imidazol-1-yl)pentanoic acid (S6).** To a solution of methyl 5-(2-((6-cyano-1H-indol-1-yl)methyl)-1H-imidazol-1-yl)pentanoate (0.37 g, 1.1 mmol) in dioxane (10 mL) was added lithium hydroxide (0.13 g, 5.5 mmol) and water (1 mL). The reaction was heated to 40 °C for 2 h. The mixture diluted with water, the pH was adjusted to 3 with HCl, and extracted with chloroform/isopropanol (3:1). Concentration of the organic layer afforded the crude product (0.30 g) as a white solid. ESI MS m/z 323 [M + 1]+.

***tert*-Butyl (17-(2-((6-cyano-1H-indol-1-yl)methyl)-1H-imidazol-1-yl)-13-oxo-3,6,9-trioxa-12-azaheptadecyl)carbamate (S7).** To a solution of 5-(2-((6-cyano-1H-indol-1-yl)methyl)-1H-imidazol-1-yl)pentanoic acid (0.30 g, 0.93 mmol) in DMF (10 mL) was added tert-butyl (2-(2-(2-(2-aminoethoxy)ethoxy)ethoxy)ethyl)carbamate (0.33 g, 1.1 mmol), hydroxybenzotriazole (0.28 g, 1.9 mmol), 1-ethyl-3-(3ʹ-dimethylaminopropyl)carbodiimide, HCl (0.36 g, 1.9 mmol), and diisopropylamine (0.36 g, 2.8 mmol). The reaction was heated to 60 °C for 1 h. The reaction was diluted with ethyl acetate and washed with water. The organic layers were combined, dried with sodium sulfate, filtered, concentrated, and purified with silica gel chromatography to afford the desired product (525 mg, 94%) as a light brown oil. ^1^H NMR (400 MHz, Methanol-d4) δ 7.96 (s, 1H), 7.89 (s, 1H), 7.73 (d, *J* = 8.5, 1H), 7.54 – 7.48 (m, 1H), 7.35 (d, *J* = 8.2 Hz, 1H), 7.17 (s, 1H), 7.03 (s, 1H), 6.67 (s, 1H), 6.60 (s, 1H), 5.58 (s, 2H), 3.90 (t, *J* = 7.3 Hz, 2H), 3.67 – 3.56 (m, 8H), 3.56 – 3.45 (m, 4H), 3.36 (s, 3H), 3.26 – 3.17 (m, 2H), 2.02 (t, *J* = 6.6 Hz, 2H), 1.43 (s, 9H), 1.40 – 1.24 (m, 4H); ^13^C NMR (101 MHz, MeOD) δ 173.94, 157.07, 142.37, 135.14, 132.24, 131.74, 126.72, 122.12, 121.65, 121.55, 120.02, 114.69, 103.70, 102.71, 78.69, 70.17, 70.14, 69.80, 69.76, 69.66, 69.13, 48.47, 45.65, 42.20, 39.99, 39.86, 38.93, 34.70, 29.78, 27.38, 22.25; HRMS (ESI+) calcd for C_31_H_44_N_6_O_6_ [M + H]+ m/z 597.3400, found 597.3384; HPLC 99.0% (AUC at 286 nm) 3.53 min (Synergi Max-RP, water/ACN, 0.02%TFA).

***tert*-Butyl (17-(2-((6-(aminomethyl)-1H-indol-1-yl)methyl)-1H-imidazol-1-yl)-13-oxo-3,6,9-trioxa-12-azaheptadecyl)carbamate (S8).** To a solution of tert-butyl (17-(2-((6-cyano-1H-indol-1-yl)methyl)-1H-imidazol-1-yl)-13-oxo-3,6,9-trioxa-12-azaheptadecyl)carbamate (0.52 g, 0.88 mmol) in ammonia in methanol (7 N, 20 mL) was added a scoop of Rainey Nickel suspended in water. The reaction was charged with hydrogen (60 psi) and stirred at ambient temperature for 18 h. After degassing with nitrogen, the mixture was filtered over celite. The filtrate was concentrated to afford the crude product (0.49 g) as a colorless oil. ESI MS m/z 601 [M + 1]+.

***tert*-Butyl (17-(2-((6-((((3-(6-hydroxy-3-oxoisoindolin-1-yl)-1H-indol-2-yl)methyl)amino)methyl)-1H-indol-1-yl)methyl)-1H-imidazol-1-yl)-13-oxo-3,6,9-trioxa-12-azaheptadecyl)carbamate (S10).** To a suspension of tert-butyl (17-(2-((6-(aminomethyl)-1H-indol-1-yl)methyl)-1H-imidazol-1-yl)-13-oxo-3,6,9-trioxa-12-azaheptadecyl)carbamate (0.45 g, 0.75 mmol) in THF (10 mL) was added 3-(6-hydroxy-3-oxoisoindolin-1-yl)-1H-indole-2-carbaldehyde (**S9**, 0.22 g, 0.75 mmol)^8^. The reaction stirred at ambient temperature for 1 h. Sodium triacetoxyborohydride (0.48 g, 2.3 mmol) was added and stirred at ambient temperature for 3 h. The reaction was diluted with methanol, celite was added, concentrated, and purified with silica gel chromatography to afford the desired product (0.53 g, 80%) as a brown foam. ^1^H NMR (400 MHz, Methanol-d4) δ 7.71 (d, *J* = 8.3 Hz, 1H), 7.62 (d, *J* = 8.3 Hz, 2H), 7.38 (d, *J* = 8.2 Hz, 1H), 7.27 (s, 1H), 7.18 (d, *J* = 8.2 Hz, 1H), 7.14 – 7.04 (s, 2H), 7.00 – 6.87 (m, 2H), 6.87 – 6.77 (m, 1H), 6.77 – 6.73 (m, 2H), 6.54 (s, 1H), 5.95 (s, 1H), 5.48 (s, 2H), 4.55 – 4.32 (m, 2H), 4.24 (s, 2H), 3.92 – 3.79 (m, 2H), 3.63 – 3.49 (m, 8H), 3.46 (t, *J* = 5.6 Hz, 2H), 3.40 (t, *J* = 5.6 Hz, 2H), 3.27 – 3.14 (m, 4H), 2.22 – 1.72 (m, 6H), 1.42 (s, 9H), 1.32 – 1.08 (m, 4H); ^13^C NMR (101 MHz, MeOD) δ 175.62, 174.06, 171.66, 161.97, 157.02, 150.94, 142.98, 136.69, 136.20, 129.32, 128.99, 128.73, 126.49, 126.38, 125.44, 124.47, 122.76, 122.55, 121.41, 121.29, 120.77, 119.39, 118.80, 115.88, 111.29, 111.04, 110.77, 109.77, 102.09, 78.70, 70.10, 69.76, 69.72, 69.62, 69.04, 52.85, 51.80, 45.65, 41.80, 41.33, 39.84, 38.93, 34.65, 29.76, 27.38, 22.21, 20.62; HRMS (ESI+) calcd for C_48_H_60_N_8_O_8_ [M + H]+ m/z 877.4612, found 877.4607; HPLC 96.7% (AUC at 266 nm) 2.79 min (Synergi Max-RP, water/ACN, 0.02%TFA).

**N-(2-(2-(2-(2-Aminoethoxy)ethoxy)ethoxy)ethyl)-5-(2-((6-((((3-(6-hydroxy-3-oxoisoindolin-1-yl)-1H-indol-2-yl)methyl)amino)methyl)-1H-indol-1-yl)methyl)-1H-imidazol-1-yl)pentanamide (S11).** To a solution of tert-butyl (17-(2-((6-((((3-(6-hydroxy-3-oxoisoindolin-1-yl)-1H-indol-2-yl)methyl)amino)methyl)-1H-indol-1-yl)methyl)-1H-imidazol-1-yl)-13-oxo-3,6,9-trioxa-12-azaheptadecyl)carbamate (0.030 g, 0.034 mmol) in dichloromethane (10 mL) was added trifluoroacetic acid (1 mL). The reaction stirred at ambient temperature for 1.5 h. The mixture was concentrated to give crude product as a colorless oil. ESI MS m/z 777 [M + 1]+.

**N-(15-(5,5-Difluoro-7-(1H-pyrrol-2-yl)-5H-5l4,6l4-dipyrrolo[1,2-c:2',1'-f][1,3,2]diazaborinin-3-yl)-13-oxo-3,6,9-trioxa-12-azapentadecyl)-5-(2-((6-((((3-(6-hydroxy-3-oxoisoindolin-1-yl)-1H-indol-2-yl)methyl)amino)methyl)-1H-indol-1-yl)methyl)-1H-imidazol-1-yl)pentanamide (S12).** To a solution of N-(2-(2-(2-(2-aminoethoxy)ethoxy)ethoxy)ethyl)-5-(2-((6-((((3-(6-hydroxy-3-oxoisoindolin-1-yl)-1H-indol-2-yl)methyl)amino)methyl)-1H-indol-1-yl)methyl)-1H-imidazol-1-yl)pentanamide (26 mg, 0.033 mmol) in DMF (2 mL) was added 2,5-dioxopyrrolidin-1-yl 3-(5,5-difluoro-7-(1H-pyrrol-2-yl)-5H-5l4,6l4-dipyrrolo[1,2-c:2',1'-f][1,3,2]diazaborinin-3-yl)propanoate (14 mg, 0.033 mmol) and diisopropylethylamine (34 mg, 0.27 mmol). The reaction was stirred at ambient temperature for 30 min. The mixture was diluted with methanol and purified by reverse phase preparative HPLC to afford the desired product (38 mg, quant) as a purple solid. ^1^H NMR (400 MHz, Methanol-d4) δ 7.71 (d, *J* = 8.2 Hz, 1H), 7.54 (d, *J* = 8.2 Hz, 1H), 7.46 (s, 1H), 7.32 (d, *J* = 8.6 Hz, 1H), 7.22 – 7.07 (m, 6H), 7.07 – 6.95 (m, 3H), 6.95 – 6.85 (m, 3H), 6.79 (t, *J* = 7.6 Hz, 1H), 6.75 – 6.64 (m, 2H), 6.47 (s, 1H), 6.37 – 6.27 (m, 2H), 5.92 (s, 1H), 5.41 (s, 2H), 4.00 (s, 2H), 3.89 (s, 2H), 3.72 (t, *J* = 7.5 Hz, 2H), 3.62 – 3.45 (m, 10H), 3.43-3.34 (m, 4H), 3.30 – 3.24 (d, 2H), 3.21 (t, *J* = 5.6 Hz, 2H), 2.63 (t, *J* = 7.8 Hz, 2H), 1.80 (t, *J* = 7.3 Hz, 2H), 1.25 – 1.13 (m, 2H), 1.13 – 1.01 (m, 2H); HRMS (ESI+) calcd for C_59_H_64_BF_2_N_11_O_7_ [M + H]+ m/z 1088.5129, found 1088.5124; HPLC 95.9% (AUC at 580 nm) 3.36 min (Synergi Max-RP, water/ACN, 0.02%TFA).

*Preparation of piperazine precursor of MRTX849*

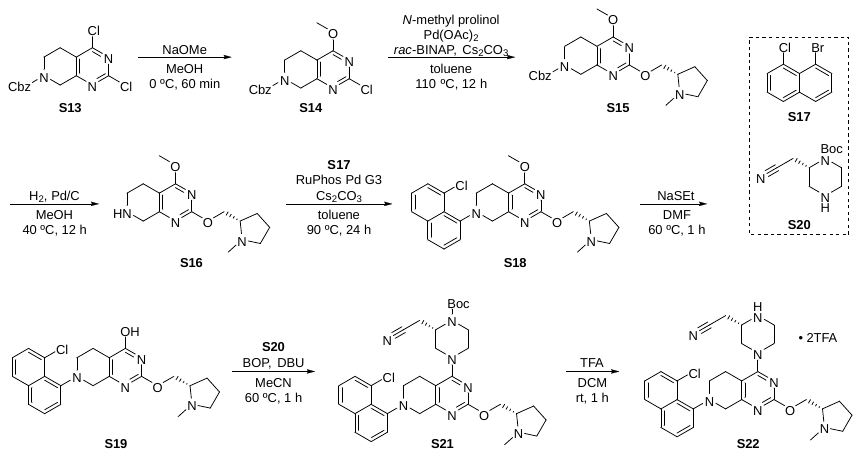

Supplementary Scheme 2. Preparation of piperazine precursor of MRTX849.

Synthesis of MRTX849 precursor was adapted from the published syntheses.^9,10^

**Benzyl 2-chloro-4-methoxy-5,8-dihydropyrido[3,4-*d*]pyrimidine-7(6*H*)-carboxylate (S14).** A 250-mL round bottom flask equipped with a stir bar was charged with benzyl 2,4-dichloro-5,8-dihydropyrido[3,4-*d*]pyrimidine-7(6*H*)-carboxylate (**S13**, 4.78 g, 14.1 mmol). Methanol (100 mL) was added, and the resulting yellow suspension was stirred at 0 °C. Sodium methoxide (25 wt% in MeOH, 3.87 mL, 17.0 mmol) was added via syringe. The resulting mixture was stirred at 0 °C for 60 min, at which point the full conversion was judged complete by TLC. The reaction mixture was neutralized by 2 N HCl and then concentrated. The residue was partitioned between EA (100 mL) and water (100 mL). The phases were separated, and the aqueous phase was extracted by EA (100 mL x 2). The organic solution was combined, washed by brine and dried on anhydrous sodium sulfate. Upon concentration, the solid crude was triturated in ether, filtered, and dried in vacuum (50 mTorr) overnight. The title product was obtained as a white powder (4.39 g, 13.1 mmol, 93% yield). ^1^H NMR (400 MHz, Chloroform-*d*) δ 7.44 – 7.28 (m, 5H), 5.16 (s, 2H), 4.59 (s, 2H), 4.02 (s, 3H), 3.73 (t, *J* = 5.8 Hz, 2H), 2.65 (s, 2H). TLC Rf 0.69 (50% EA-hexanes, UV). Accurate MS (ESI-TOF) calculated for C_16_H_17_ClN_3_O_3_ [M + H]^+^ 334.0958, found 334.1016.

**Benzyl (*S*)-4-methoxy-2-((1-methylpyrrolidin-2-yl)methoxy)-5,8-dihydropyrido[3,4-*d*]pyrimidine-7(6*H*)-carboxylate (S15).** A 100-mL round bottom flask equipped with a stir bar was charged with benzyl 2-chloro-4-methoxy-5,8-dihydropyrido[3,4-*d*]pyrimidine-7(6*H*)-carboxylate (1.67 g, 5.00 mmol), (*S*)-(1-methylpyrrolidin-2-yl)methanol (1.15 g, 10.0 mmol, 2.0 equiv), palladium acetate (112 mg, 0.500 mmol, 0.10 equiv), *rac*-BINAP (623 mg, 1.00 mmol, 0.20 equiv), cesium carbonate (4.89 g, 15.0 mmol, 3.0 equiv). Sealed with a rubber septum, the flask was repeatedly deaerated and backfilled with argon three times. Toluene (25 mL) was added. The mixture was heated to 110 °C under argon atmosphere for 12 h when LC-MS analysis indicated the full conversion of the limiting starting material, pyrimidine chloride. The reaction mixture was directly partitioned between EA (100 mL) and water (100 mL). The two phases were separated and the aqueous was further extracted with EA (100 mL x 2). The combined organic solution was washed with brine, dried over anhydrous sodium sulfate, and concentrated. The crude was purified by flash column chromatography (0–20% MeOH-DCM gradient with 0.2% ammonia, 80 g RediSep(R) Rf column, Teledyne ISCO, Lincoln, NE). The title compound was obtained as a yellow solid (1.5230 g, 3.693 mmol, 74% yield). TLC Rf 0.45 (17% MeOH-DCM, UV). Proton NMR was consistent to the earlier report^11^. Accurate MS (ESI-TOF) calculated for C_22_H_29_N_4_O_4_ [M + H]^+^ 413.2188, found 413.2235.

**(*S*)-4-Methoxy-2-((1-methylpyrrolidin-2-yl)methoxy)-5,6,7,8-tetrahydropyrido[3,4-*d*]pyrimidine (S16).** A 100-mL round bottom flask equipped with a stir bar was charged with benzyl (*S*)-4-methoxy-2-((1-methylpyrrolidin-2-yl)methoxy)-5,8-dihydropyrido[3,4-*d*]pyrimidine-7(6*H*)-carboxylate (1.24 g, 3.00 mmol), palladium on carbon (1.12 g, 10 wt% Pd, wet basis). Sealed with a rubber septum, the flask was repeatedly deaerated and backfilled with argon three times. Methanol was added via syringe. Hydrogen gas in a balloon was introduced via a long needle by submerging the needle tip in the stirred suspension. The hydrogen flow-through was maintained for 10 min before the hydrogen gas needle retracted into the headspace above the suspension and the exit needle removed from the septum. The black suspension was stirred under hydrogen atmosphere at 40 °C for 12 h when LC-MS indicated the full conversion of the starting material. An argon stream was bubbled through the reaction suspension before the latter been filtered through a tightly packed Celite column. The filter cake was washed with methanol (20 mL x 2). The combined filtrate was concentrated to give the title product as a yellow solid (752 mg, 2.70 mmol, 90% yield). Proton NMR was consistent with the earlier report.^10^ Accurate MS (ESI-TOF) calculated for C_14_H_23_N_4_O_2_ [M + H]^+^ 279.1821, found 279.1824.

**(*S*)-7-(8-Chloronaphthalen-1-yl)-4-methoxy-2-((1-methylpyrrolidin-2-yl)methoxy)-5,6,7,8-tetrahydropyrido[3,4-*d*]pyrimidine (S18).** A 100-mL round bottom flask equipped with a stir bar was charged with (*S*)-4-methoxy-2-((1-methylpyrrolidin-2-yl)methoxy)-5,6,7,8-tetrahydropyrido[3,4-*d*]pyrimidine (752 mg, 2.70 mmol), 1-bromo-8-chloronaphthalene (**S17**, 848 mg, 3.51 mmol, 1.3 equiv), RuPhos Pd G3 precatalyst (226 mg, 0.270 mmol, 0.10 equiv), and cesium carbonate (3.08 g, 9.45 mmol, 3.5 equiv). Sealed with a rubber septum, the flask was repeatedly deaerated and backfilled with argon three times. Degassed toluene (13.5 mL) was added. The resulting solution was stirred at 90 °C for 24 h when LC-MS indicated the full conversion of secondary amine starting material. The black reaction mixture was concentrated. The residue was partitioned between EA (50 mL) and water (100 mL). The two phases were separated and the aqueous was extracted by EA (50 mL x 2). The combined organic solution was washed by half-saturated brine, brine, sequentially, dried over anhydrous sodium sulfate, and concentrated. The crude product was purified by flash column chromatography (0–20% MeOH-DCM gradient with 0.2% ammonia, 80-g RediSep(R) Rf column, Teledyne ISCO, Lincoln, NE). The title compound was obtained as a yellow solid (851.0 mg, 1.939 mmol, 72% yield). Proton NMR was consistent with the earlier report.^10^ TLC Rf 0.56 (17% MeOH-DCM, UV). Accurate MS (ESI-TOF) calculated for C_24_H_28_ClN_4_O_2_ [M + H]^+^ 439.1901, found 439.1918.

**(*S*)-7-(8-Chloronaphthalen-1-yl)-2-((1-methylpyrrolidin-2-yl)methoxy)-5,6,7,8-tetrahydropyrido[3,4-*d*]pyrimidin-4-ol (S19).** A 20-mL scintillation vial equipped with a stir bar was charged with (*S*)-7-(8-Chloronaphthalen-1-yl)-4-methoxy-2-((1-methylpyrrolidin-2-yl)methoxy)-5,6,7,8-tetrahydropyrido[3,4-*d*]pyrimidine (413 mg, 0.940 mmol), and sodium ethanethiolate (158 mg, 1.88 mmol, 2.0 equiv). DMF (5 mL) was added and resulting mixture was stirred at 60 °C for 3 h. The reaction mixture was concentrated before partitioned between DCM (40 mL) and 10% aqueous sodium bicarbonate (40 mL). The phases were separated and the aqueous was extracted with DCM (40 mL x 2). The combined organic phase was dried on anhydrous sodium sulfate, filtered, and concentrated. The residue was purified by flash column chromatography (0–20% MeOH-DCM gradient with 0.2% ammonia, 24-g RediSep(R) Rf column, Teledyne ISCO, Lincoln, NE). The title compound was obtained as a yellow solid (348.2 mg, 0.819 mmol, 87% yield). Proton NMR was consistent with the earlier report.^10^ TLC Rf 0.32 (17% MeOH-DCM, UV). Accurate MS (ESI-TOF) calculated for C_23_H_26_ClN_4_O_2_ [M + H]^+^ 425.1744, found 425.1774.

***tert*-Butyl (*S*)-4-(7-(8-chloronaphthalen-1-yl)-2-(((*S*)-1-methylpyrrolidin-2-yl)methoxy)-5,6,7,8-tetrahydropyrido[3,4-*d*]pyrimidin-4-yl)-2-(cyanomethyl)piperazine-1-carboxylate (S21).** A 4-mL vial equipped with a stir bar was charged with (*S*)-7-(8-chloronaphthalen-1-yl)-2-((1-methylpyrrolidin-2-yl)methoxy)-5,6,7,8-tetrahydropyrido[3,4-*d*]pyrimidin-4-ol (187 mg, 0.440 mmol) and BOP (253 mg, 0.572 mmol, 1.3 equiv). Sealed by a screw cap with a PEFE septum, the vial was deaerated and backfilled with argon three times. Anhydrous MeCN (1.1 mL) was added, and the mixture was stirred to a brown suspension. DBU (100 µL, 0.66 mmol, 1.5 equiv) was added and the suspension, immediately, turned homogeneous. The reddish solution was stirred at ambient temperature for 10 min before a stock solution of *tert*-butyl (*S*)-2-(cyanomethyl)piperazine-1-carboxylate (**S20**, 149 mg, 0.660 mmol, 1.5 equiv) in MeCN (1.1 mL) was added via syringe. The resulting solution was allowed to stir at 60 °C overnight. The reaction mixture was concentrated. The brown crude was purified by flash column chromatography (0–15% MeOH-DCM gradient, 24-g RediSep(R) Rf column, Teledyne ISCO, Lincoln, NE). The title compound was obtained as a yellow solid (213 mg, 0.338 mmol, 77% yield). Proton NMR is consistent to the earlier report by Blake, J. F. et al.^11^ Accurate MS (ESI-TOF) calculated for C_34_H_43_ClN_7_O_3_ [M + H]^+^ 632.3116, found 632.3157.

**2-((*S*)-4-(7-(8-Chloronaphthalen-1-yl)-2-(((*S*)-1-methylpyrrolidin-2-yl)methoxy)-5,6,7,8-tetrahydropyrido[3,4-*d*]pyrimidin-4-yl)piperazin-2-yl)acetonitrile (S22).** A 20-mL scintillation vial equipped with a stir bar was charged with *tert*-butyl (*S*)-4-(7-(8-chloronaphthalen-1-yl)-2-(((*S*)-1-methylpyrrolidin-2-yl)methoxy)-5,6,7,8-tetrahydropyrido[3,4-*d*]pyrimidin-4-yl)-2-(cyanomethyl)piperazine-1-carboxylate (213 mg, 0.338 mmol). DCM (2 mL) and trifluoroacetic acid (TFA, 2 mL) were added. The resulting solution was stirred at ambient temperature for 1 h when LC-MS indicated the full conversion of the starting material to the desired product. The reaction mixture was concentrated in vacuo. The crude was triturated in ether, centrifugated and decanted to result the title compound as a di-TFA salt (254 mg, 0.335 mmol, 99% yield). The compound was used in the next step without further purification. Accurate MS (ESI-TOF) calculated for C_29_H_35_ClN_7_O [M + H]^+^ 532.2592, found 532.2592.

*Preparation of saturated amide analogs (****CPD1-6****) of MRTX849*

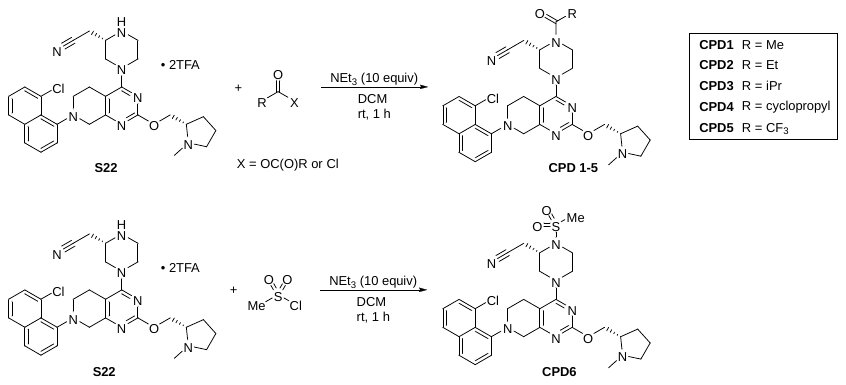

Supplementary Scheme 3. Preparation of saturated amide analogs of MRTX849.

**2-((*S*)-1-Acetyl-4-(7-(8-chloronaphthalen-1-yl)-2-(((*S*)-1-methylpyrrolidin-2-yl)methoxy)-5,6,7,8-tetrahydropyrido[3,4-*d*]pyrimidin-4-yl)piperazin-2-yl)acetonitrile (CPD1).** In a typical procedure for the synthesis of saturated amide analogs of MRTX849, A 4-mL vial equipped with a stir bar was charged with 2-((*S*)-4-(7-(8-chloronaphthalen-1-yl)-2-(((*S*)-1-methylpyrrolidin-2-yl)methoxy)-5,6,7,8-tetrahydropyrido[3,4-*d*]pyrimidin-4-yl)piperazin-2-yl)acetonitrile (5.0 mg, 0.0094 mmol) and DCM (1 mL). Triethylamine (13 µL, 0.094 mmol, 10 equiv) and acetic anhydride (4.4 µL, 0.047 mmol, 5 equiv) were added. The mixture was stirred at 23 °C for 1 h when LC-MS indicated the full conversion of the starting material. Upon concentration, the reaction crude was dissolved in 0.1 mL DMSO and diluted by 0.9 mL (50% MeCN-water). The resulting solution was filtered through a 0.45 µM PTFE syringe filter. The filtrate was purified by reverse-phase HPLC (Waters XBridge C18 column 5 µm particle size 30 x 250 mm, 5–95% acetonitrile–water + 0.1% formic acid, 40 min, 20 mL/min) to afford the title compound in its 1:1 formic acid salt form as a white fluffy solid (1.7 mg, 0.0030 mmol, 29% yield). ^1^H NMR (400 MHz, DMSO-*d*_6_) δ 7.92 (d, *J* = 8.1 Hz, 1H), 7.78 – 7.70 (m, 1H), 7.62 – 7.50 (m, 2H), 7.45 (t, *J* = 7.8 Hz, 1H), 7.34 (dd, *J* = 14.8, 6.9 Hz, 1H), 4.94 – 4.79 (m, 1H), 4.60 – 4.28 (m, 1H), 4.24 (ddt, *J* = 9.7, 4.8, 2.2 Hz, 1H), 4.16 (d, *J* = 16.1 Hz, 1H), 4.12 – 3.65 (m, 5H), 3.64 – 3.44 (m, 2H, inaccurate integrate due to HOD contamination), 3.26 – 2.62 (m, 10H), 2.55 (q, *J* = 6.8 Hz, 1H), 2.34 (d, *J* = 4.1 Hz, 3H), 2.25 – 2.12 (m, 3H), 2.07 (d, *J* = 1.6 Hz, 2H), 2.01 – 1.86 (m, 1H), 1.73 – 1.53 (m, 3H). Accurate MS (ESI-TOF) calculated for C_31_H_37_ClN_7_O_2_ [M + H]^+^ 574.2697, found 574.2703.

**2-((*S*)-4-(7-(8-chloronaphthalen-1-yl)-2-(((*S*)-1-methylpyrrolidin-2-yl)methoxy)-5,6,7,8-tetrahydropyrido[3,4-*d*]pyrimidin-4-yl)-1-propionylpiperazin-2-yl)acetonitrile (CPD2).** The title compound was synthesized via the general MRTX849 analog preparation procedure. 2-((*S*)-4-(7-(8-chloronaphthalen-1-yl)-2-(((*S*)-1-methylpyrrolidin-2-yl)methoxy)-5,6,7,8-tetrahydropyrido[3,4-*d*]pyrimidin-4-yl)piperazin-2-yl)acetonitrile (5.0 mg, 0.0094 mmol), triethylamine (13 µL, 0.094 mmol, 10 equiv) and propionyl chloride (4.1 µL, 0.047 mmol, 5 equiv) afforded the title compound in its 1:1 formic acid salt form (4.6 mg, 0.007 mmol, 77% yield) as a white fluffy solid. ^1^H NMR (400 MHz, DMSO-*d*_6_) δ 7.93 (ddt, *J* = 8.1, 1.3, 0.6 Hz, 1H), 7.75 (ddd, *J* = 8.6, 4.5, 1.1 Hz, 1H), 7.61 – 7.50 (m, 2H), 7.45 (dd, *J* = 8.1, 7.4 Hz, 1H), 7.39 – 7.29 (m, 1H), 5.00 – 4.75 (m, 1H), 4.65 – 4.32 (m, 1H), 4.24 (ddd, *J* = 9.5, 4.8, 2.8 Hz, 1H), 4.16 (d, *J* = 15.5 Hz, 1H), 4.11 – 3.88 (m, 3H), 3.78 (dd, *J* = 36.6, 14.4 Hz, 3H), 3.64 – 2.62 (m, 15H, inaccurate integrate due to HOD contamination), 2.56 (h, *J* = 5.3 Hz, 1H), 2.35 (d, *J* = 4.2 Hz, 3H), 2.19 (qd, *J* = 8.7, 3.1 Hz, 1H), 2.00 – 1.86 (m, 1H), 1.73 – 1.55 (m, 3H), 1.02 (td, *J* = 11.3, 9.8, 6.6 Hz, 3H). Accurate MS (ESI-TOF) calculated for C_32_H_39_ClN_7_O_2_ [M + H]^+^ 588.2854, found 588.2875.

**2-((*S*)-4-(7-(8-Chloronaphthalen-1-yl)-2-(((*S*)-1-methylpyrrolidin-2-yl)methoxy)-5,6,7,8-tetrahydropyrido[3,4-*d*]pyrimidin-4-yl)-1-isobutyrylpiperazin-2-yl)acetonitrile (CPD3).** The title compound was synthesized via the general MRTX849 analog preparation procedure. 2-((*S*)-4-(7-(8-chloronaphthalen-1-yl)-2-(((*S*)-1-methylpyrrolidin-2-yl)methoxy)-5,6,7,8-tetrahydropyrido[3,4-*d*]pyrimidin-4-yl)piperazin-2-yl)acetonitrile (5.0 mg, 0.009 mmol), triethylamine (13 µL, 0.094 mmol, 10 equiv) and isobutyryl chloride (4.9 µL, 0.047 mmol, 5 equiv) afforded the title compound in its 1:1 formic acid salt form (3.3 mg, 0.005 mmol, 54% yield) as a white fluffy solid. ^1^H NMR (400 MHz, DMSO-*d*_6_) δ 7.93 (dt, *J* = 8.4, 1.0 Hz, 1H), 7.75 (ddd, *J* = 8.4, 4.0, 1.1 Hz, 1H), 7.62 – 7.50 (m, 2H), 7.46 (dd, *J* = 8.1, 7.4 Hz, 1H), 7.40 – 7.30 (m, 1H), 5.01 – 4.87 (m, 1H), 4.29 – 4.12 (m, 2H), 4.10 – 3.89 (m, 3H), 3.89 – 3.68 (m, 2H), 3.64 – 2.78 (m, 20H), 2.71 (t, *J* = 14.5 Hz, 1H), 2.34 (d, *J* = 3.8 Hz, 3H), 2.17 (qd, *J* = 8.7, 2.5 Hz, 1H), 1.92 (dq, *J* = 12.1, 8.2 Hz, 1H), 1.73 – 1.52 (m, 3H), 1.11 – 0.97 (m, 6H). Accurate MS (ESI-TOF) calculated for C_33_H_41_ClN_7_O_2_ [M + H]^+^ 602.3010, found 602.3030.

**2-((*S*)-4-(7-(8-Chloronaphthalen-1-yl)-2-(((*S*)-1-methylpyrrolidin-2-yl)methoxy)-5,6,7,8-tetrahydropyrido[3,4-*d*]pyrimidin-4-yl)-1-(cyclopropanecarbonyl)piperazin-2-yl)acetonitrile (CPD4).** The title compound was synthesized via the general MRTX849 analog preparation procedure. 2-((*S*)-4-(7-(8-chloronaphthalen-1-yl)-2-(((*S*)-1-methylpyrrolidin-2-yl)methoxy)-5,6,7,8-tetrahydropyrido[3,4-*d*]pyrimidin-4-yl)piperazin-2-yl)acetonitrile (5.0 mg, 0.009 mmol), triethylamine (13 µL, 0.094 mmol, 10 equiv) and cyclopropanecarbonyl chloride (4.3 µL, 0.047 mmol, 5 equiv) afforded the title compound in its 1:1 formic acid salt form (2.4 mg, 0.004 mmol, 40% yield) as a white fluffy solid. ^1^H NMR (400 MHz, DMSO-*d*_6_) δ 7.93 (dt, *J* = 8.5, 1.2 Hz, 1H), 7.75 (ddd, *J* = 8.6, 4.5, 1.1 Hz, 1H), 7.62 – 7.50 (m, 2H), 7.46 (dd, *J* = 8.1, 7.4 Hz, 1H), 7.40 – 7.30 (m, 1H), 4.93 (s, 1H), 4.45 – 4.11 (m, 3H), 4.13 – 3.91 (m, 3H), 3.79 (dd, *J* = 43.3, 17.3 Hz, 3H), 3.58 – 2.99 (m, 16H, inaccurate integrate due to HOD contamination), 2.99 – 2.79 (m, 3H), 2.71 (s, 1H), 2.35 (d, *J* = 4.1 Hz, 3H), 2.18 (qd, *J* = 8.7, 2.8 Hz, 1H), 2.13 – 1.98 (m, 1H), 1.93 (dq, *J* = 12.1, 8.1 Hz, 1H), 1.75 – 1.54 (m, 3H), 0.90 – 0.70 (m, 5H). Accurate MS (ESI-TOF) calculated for C_33_H_39_ClN_7_O_2_ [M + H]^+^ 600.2853, found 600.2853.

**2-((*S*)-4-(7-(8-Chloronaphthalen-1-yl)-2-(((*S*)-1-methylpyrrolidin-2-yl)methoxy)-5,6,7,8-tetrahydropyrido[3,4-*d*]pyrimidin-4-yl)-1-(2,2,2-trifluoroacetyl)piperazin-2-yl)acetonitrile (CPD5).** The title compound was synthesized via the general MRTX849 analog preparation procedure. 2-((*S*)-4-(7-(8-chloronaphthalen-1-yl)-2-(((*S*)-1-methylpyrrolidin-2-yl)methoxy)-5,6,7,8-tetrahydropyrido[3,4-*d*]pyrimidin-4-yl)piperazin-2-yl)acetonitrile (5.0 mg, 0.009 mmol), triethylamine (13 µL, 0.094 mmol, 10 equiv) and trifluoroacetic anhydride (8.6 µL, 0.047 mmol, 5 equiv) afforded the title compound in its 1:1 formic acid form (1.9 mg, 0.003 mmol, 30% yield) as a white fluffy solid. ^1^H NMR (400 MHz, DMSO-*d*_6_) δ 7.96 – 7.90 (m, 1H), 7.75 (ddd, *J* = 8.4, 3.4, 1.1 Hz, 1H), 7.61 – 7.50 (m, 2H), 7.46 (dd, *J* = 8.1, 7.5 Hz, 1H), 7.40 – 7.31 (m, 1H), 4.92 (s, 1H), 4.36 – 3.57 (m, 8H), 3.50 (q, *J* = 7.2 Hz, 2H), 3.20 – 2.88 (m, 6H), 2.71 (d, *J* = 14.4 Hz, 1H), 2.56 (d, *J* = 6.1 Hz, 1H), 2.36 (d, *J* = 4.1 Hz, 3H), 2.20 (td, *J* = 8.6, 3.6 Hz, 1H), 2.01 – 1.87 (m, 1H), 1.75 – 1.53 (m, 3H), 1.15 (dt, *J* = 23.3, 7.2 Hz, 1H). Accurate MS (ESI-TOF) calculated for C_31_H_34_ClF_3_N_7_O_2_ [M + H]^+^ 628.2414, found 628.2401.

**2-((*S*)-4-(7-(8-chloronaphthalen-1-yl)-2-(((*S*)-1-methylpyrrolidin-2-yl)methoxy)-5,6,7,8-tetrahydropyrido[3,4-*d*]pyrimidin-4-yl)-1-(methylsulfonyl)piperazin-2-yl)acetonitrile (CPD6).** The title compound was synthesized via the general MRTX849 analog preparation procedure. 2-((*S*)-4-(7-(8-chloronaphthalen-1-yl)-2-(((*S*)-1-methylpyrrolidin-2-yl)methoxy)-5,6,7,8-tetrahydropyrido[3,4-*d*]pyrimidin-4-yl)piperazin-2-yl)acetonitrile (5.0 mg, 0.009 mmol), triethylamine (13 µL, 0.094 mmol, 10 equiv) and methanesulfonyl chloride (3.6 µL, 0.047 mmol, 5 equiv) afforded the title compound in its 1:1 formic acid salt form (1.2 mg, 0.002 mmol, 19% yield) as a white fluffy solid. ^1^H NMR (400 MHz, DMSO-*d*_6_) δ 7.93 (dt, *J* = 8.5, 1.2 Hz, 1H), 7.75 (ddd, *J* = 8.4, 4.8, 1.1 Hz, 1H), 7.62 – 7.50 (m, 2H), 7.46 (dd, *J* = 8.1, 7.4 Hz, 1H), 7.35 (ddd, *J* = 20.4, 7.6, 1.2 Hz, 1H), 4.36 (s, 1H), 4.32 – 4.25 (m, 1H), 4.23 – 4.08 (m, 2H), 4.02 – 3.90 (m, 2H), 3.86 – 3.64 (m, 3H), 3.51 (q, *J* = 10.6 Hz, 3H), 3.26 – 2.90 (m, 12H), 2.68 (d, *J* = 14.4 Hz, 2H), 2.42 (d, *J* = 5.8 Hz, 3H), 2.36 – 2.25 (m, 1H), 1.97 (dq, *J* = 12.1, 8.4, 8.0 Hz, 1H), 1.78 – 1.56 (m, 3H). Accurate MS (ESI-TOF) calculated for C_30_H_37_ClN_7_O_3_S [M + H]^+^ 610.2367, found 610.2371.

*Preparation of EX185*

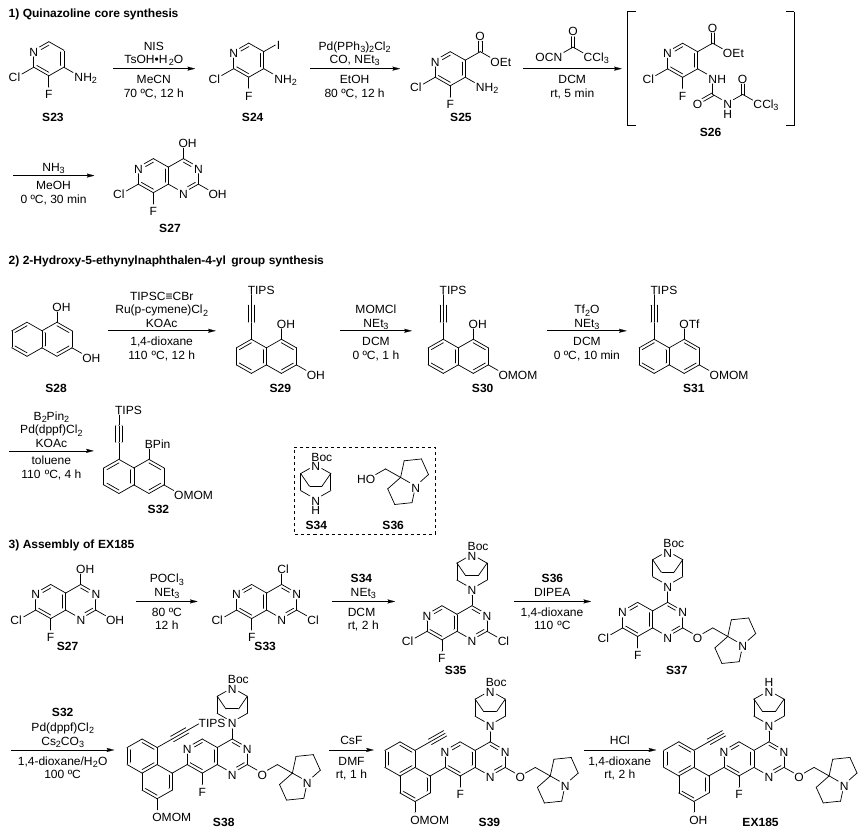

Supplementary Scheme 4. Preparation of EX185.

Synthesis of EX185 was adapted from the patent application US2020/048194^12^ with modifications.

**2-Chloro-3-fluoro-5-iodopyridin-4-amine (S24).** A 20-mL scintillation vial equipped with a stir bar was charged with commercially available 2-chloro-3-fluoro-pyridin-4-amine (**S23**, 1.00 g, 6.82 mmol), *N*-Iodosuccinimide (1.84 g, 8.19 mmol), *p*-toluenesulfonic acid monohydrate (64.9 mg, 0.341 mmol) and MeCN (5 mL). Vigorous stirring at 70 °C turned the insoluble suspension into a red color homogenous solution. The conversion was complete after 12 h. The solvent was removed and the residue was purified by flash column chromatography to give 2-chloro-3-fluoro-5-iodo-pyridin-4-amine (1.82 g, 6.67 mmol, 98% yield) as an orange crystalline solid. ^1^H NMR (400 MHz, CDCl_3_) δ 8.17 (d, *J* = 0.7 Hz, 1H), 4.85 (s, 2H). Accurate MS (ESI-TOF) calculated for C_5_H_4_ClFIN_2_ [M + H]^+^ 272.9092, found 272.9138.

**Ethyl 4-amino-6-chloro-5-fluoronicotinate (S25).** A 150-mL round bottom flask equipped with a stir bar was charged with **S24** (1.80 g, 6.60 mmol), bis(triphenylphosphine)palladium chloride (463 mg, 0.660 mmol), and ethanol (33 mL). The flask affixed with a rubber septum was deaerated and backfilled with argon three times. A balloon filled with CO gas was introduced into the reaction flask using a long needle, and the needle tip was submerged in the stirring suspension to saturate the solution with CO gas and initiate the reaction. CO gas was flowed through the suspension, with vigorous stirring, for 10 min. The exit needle was vented to the air to allow the CO gas to flow from the balloon, through the reaction mixture, and out of the vessel. The reaction mixture was stirred at 80 °C; it started from an orange solution and turned to a dark solution with some white precipitates at the bottom. LC-MS suggested complete conversion of starting material. The reaction mixture was concentrated and partitioned between EA (100 mL) and water (100 mL). The aqueous phase was further extracted with EA (50 mL x 3). The combined organic phase was washed with brine, dried on anhydrous sodium sulfate, and concentrated. The concentrated crude was purified by flash column chromatography (0–50% EA-hexanes gradient, 40-g RediSep(R) Rf column, Teledyne ISCO, Lincoln, NE) to give the title compound (1.07 g, 4.91 mmol, 74% yield) as a yellow crystalline solid. ^1^H NMR (400 MHz, CDCl_3_) δ 8.53 (s, 1H), 4.38 (q, *J* = 7.1 Hz, 2H), 1.40 (t, *J* = 7.1 Hz, 3H). ^19^F NMR (376 MHz, CDCl_3_) δ -145.50. Accurate MS (ESI-TOF) calculated for C_8_H_9_ClFIN_2_O_2_ [M + H]^+^ 219.0336, found 219.0383.

**Ethyl 6-chloro-5-fluoro-4-(3-(2,2,2-trichloroacetyl)ureido)nicotinate (S26).** A 20-mL scintillation vial equipped with a stir bar was charged with **S25** (655.8 mg, 3.000 mmol) and THF (5 mL). The vial sealed with a screw cap with silicone septum was deaerated and backfilled with argon 3 times. 2,2,2-Trichloroacetyl isocyanate (0.43 mL, 3.6 mmol) was added via a syringe and the reaction was stirred at ambient temperature for 10 min. LC-MS indicated the completion of the reaction, single peak on UV trace. The reaction mixture was concentrated. The solid red residue was triturated in diethyl ether. The title compound (1.1210 g, 2.7543 mmol, 92% yield) was obtained by filtration as a grey powder. Accurate MS (ESI-TOF) calculated for C_11_H_9_Cl_4_FN_3_O_4_ [M + H]^+^ 405.9331, found 405.9284.

**7-Chloro-8-fluoropyrido[4,3-d]pyrimidine-2,4-diol (S27).** A 20-mL scintillation vial equipped with a stir bar was charged with **S26** (1098.9 mg, 2.7000 mmol). Affixed with a rubber septum, the vial was deaerated and backfilled with argon three times. Methanol (13.5 mL) and 7N ammonia (1.54 mL, 10.8 mmol) were added. The resulting milky suspension was stirred at ambient temperature for 1 h when LC-MS indicated the completion of the reaction. The reaction mixture was concentrated *in vacuo*. The solid residue was triturated in MTBE (10 mL), filtered and further washed by MTBE (4 mL) to give the title compound (643.8 mg, 2.987 mmol, 110% yield) as an off-white solid. ^1^H NMR (400 MHz, DMSO-*d*_6_) δ 8.33 (s, 1H), 3.17 (s, 2H). Accurate MS (ESI-TOF) calculated for C_7_H_4_ClFN_3_O_2_ [M + H]^+^ 215.9976, found 215.9794.

**8-((Triisopropylsilyl)ethynyl)naphthalene-1,3-diol (S29).** A Schlenck tube equipped with a stir bar was charged with naphthalene-1,3-diol (1000.0 mg, 6.2434 mmol), 2-bromoethynyl(triisopropyl)silane (1957.4 mg, 7.4916 mmol), dichloro(*p*-cymene)ruthenium(II) dimer (382.3 mg, 0.6243 mmol), and potassium acetate (1.23 g, 12.5 mmol). The tube was deaerated and backfilled with argon three times. Degassed 1,4-dioxane (12.5 mL) was added and the reaction mixture was brought to 110 °C with an external oil bath. The stirring was allowed to continue at 110 °C under argon for 12 h. LC-MS indicated the full conversion of the starting material (12 h). The reaction mixture was partitioned between EA (100 mL) and water (100 mL). The separation of phases after agitation was difficult and slow. The organic phase was washed with brine, dried on anhydrous sodium sulfate and concentrated. The crude was purified by flash column chromatography (0–25% EA-hexanes, 24-g RediSep(R) Rf column, Teledyne ISCO, Lincoln, NE) to give the title compound as a grey solid (939.0 mg, 2.7574 mmol, 44% yield). ^1^H NMR (400 MHz, CDCl_3_) δ 9.29 (s, 1H), 7.62 (dd, *J* = 8.3, 1.2 Hz, 1H), 7.46 (dd, *J* = 7.2, 1.2 Hz, 1H), 7.29 (dd, *J* = 8.3, 7.1 Hz, 1H), 6.74 (d, *J* = 2.5 Hz, 1H), 6.62 (d, *J* = 2.5 Hz, 1H), 1.22 – 1.14 (m, 21H). Accurate MS calculated for C_21_H_29_O_2_Si [M + H]^+^ 341.1937, found 341.1924.

**3-(Methoxymethoxy)-8-((triisopropylsilyl)ethynyl)naphthalen-1-ol (S30).** A 25-mL round bottom flask equipped with a stir bar was charged with **S29** (900.1 mg, 2.643 mmol). Affixed with a rubber septum, the flask was deaerated and backfilled with argon three times. DCM (8.8 mL) and *N*,*N*-diisopropylethylamine (1.38 mL, 7.93 mmol) were added. The resulting solution was chilled to 0 °C in an ice-water bath. Chloromethyl methyl ether (3.5 M in toluene, 1.13 mL, 4.0 mmol, 1.5 equiv) was added via a syringe dropwise. The reaction mixture was stirred at 0 °C for 1 h and was directly loaded onto a silica cartridge, which was then purified by flash column chromatography (0–20% EA-hexanes gradient over 15 min then hold for 10 min, 40-g RediSep(R) Rf column, Teledyne ISCO, Lincoln, NE, 40 mL/min flow rate) to give the title compound as a yellow oil (673.1 mg, 1.750 mmol, 66% yield). ^1^H NMR (400 MHz, CDCl_3_) δ 9.25 (s, 1H), 7.72 – 7.65 (m, 1H), 7.49 (dd, *J* = 7.1, 1.2 Hz, 1H), 7.30 (dd, *J* = 8.3, 7.1 Hz, 1H), 6.97 (d, *J* = 2.4 Hz, 1H), 6.76 (d, *J* = 2.4 Hz, 1H), 5.26 (s, 2H), 3.50 (s, 3H), 1.24 – 1.14 (m, 21H).

**3-(Methoxymethoxy)-8-((triisopropylsilyl)ethynyl)naphthalen-1-yl trifluoromethanesulfonate (S31).** A 20-mL scintillation vial equipped with a stir bar was charged with **S30** (660.2 mg, 1.717 mmol). The vial affixed with a rubber septum was deaerated and backfilled with argon (three times). DCM (7.8 mL) and *N*,*N*-diisopropylethylamine (0.90 mL, 5.1 mmol, 3.0 equiv) were added. The resulting mixture was cooled to –40 °C using an external acetonitrile-dry ice bath. Trifluoromethanesulfonic anhydride (0.43 mL, 2.6 mmol, 1.5 equiv) was added slowly via a syringe with the needle immersed under the solution surface. A yellow solution was obtained and was stirred for 30 min at –40 °C. TLC indicated the full conversion of the starting material resulting in a slightly more polar spot on TLC (*R*_f_ 0.25, 17% EA-hexanes). The reaction mixture was quenched with water (5 mL) and was extracted with DCM (10 mL). The combined organic solution was dried over anhydrous sodium sulfate. The solution was concentrated; the crude was purified by column chromatography on silica (0–15% EA-hexanes, 24-g RediSep(R) Rf column, Teledyne ISCO, Lincoln, NE) to give the title compound (676.8 mg, 1.310 mmol, 76% yield) as a yellow syrup. ^1^H NMR (400 MHz, CDCl_3_) δ 7.76 – 7.71 (m, 2H), 7.46 – 7.40 (m, 2H), 7.31 (d, *J* = 2.4 Hz, 1H), 5.29 (s, 2H), 3.52 (s, 3H), 1.36 – 1.09 (m, 21H).

**Triisopropyl((6-(methoxymethoxy)-8-(4,4,5,5-tetramethyl-1,3,2-dioxaborolan-2-yl)naphthalen-1-yl)ethynyl)silane (S32).** A 4-mL screw vial equipped with a stir bar was charged with **S31** (48.0 mg, 0.0929 mmol), bis(pinacolato)diboron (47.2 mg, 0.186 mmol), [1,1’-bis(diphenylphosphino)ferrocene]palladium(II) dichloride (6.8 mg, 0.0093 mmol), and potassium acetate (31.9 mg, 0.325 mmol). The screw cap was changed to a PTFE septum and the vial was deaerated and backfilled with argon three times. Toluene (0.37 mL) was added, and the stirred mixture was brought to 110 °C, upon which the reaction turned a black mixture. After 4 h, the reaction mixture was cooled to ambient temperature and directly partitioned between EA (20 mL) and water (20 mL). The phases were separated; the aqueous solution was further extracted with EA (20 mL x 3). The combined organic solution was washed with brine and dried over anhydrous sodium sulfate. Upon filtration, the solution was concentrated in vacuo and purified by flash column chromatography (0–10% EA-Hexanes gradient, 4-g RediSep(R) Rf column, Teledyne ISCO, Lincoln, NE) to give the title compound (24.9 mg, 0.0503 mmol, 54% yield) as a yellow solid. ^1^H NMR (400 MHz, CDCl_3_) δ 7.71 – 7.66 (m, 2H), 7.47 (d, *J* = 2.6 Hz, 1H), 7.37 (d, *J* = 2.6 Hz, 1H), 7.34 (dd, *J* = 8.2, 7.2 Hz, 1H), 5.28 (s, 2H), 3.50 (s, 3H), 1.43 (s, 12H), 1.15 (d, *J* = 2.1 Hz, 21H).

***tert*-Butyl (1R,5S)-3-(2,7-dichloro-8-fluoropyrido[4,3-d]pyrimidin-4-yl)-3,8-diazabicyclo[3.2.1]octane-8-carboxylate (S35).** A 25-mL round bottom flask equipped with a stir bar was charged with **S27** (215.6 mg, 1.000 mmol). The flask affixed with a rubber septum was deaerated and backfilled with argon three times and cooled to 0 °C. Phosphoryl chloride (4.7 mL, 50 mmol, 50 equiv) and *N*,*N*-diisopropylethylamine (0.9 mL, 5 mmol) were added via syringes sequentially. The resulting mixture was heated to 110 °C for 6 h before concentrated *in vacuo*. Residual POCl_3_ was azeotropically removed with chloroform to give crude 2,4,7-trichloro-8-fluoro-pyrido[4,3-d]pyrimidine (**S33**) as a black syrup. This crude sample was used in the following step without purification or characterization.

Crude 2,4,7-trichloro-8-fluoro-pyrido[4,3-d]pyrimidine (**S33**) dissolved in DCM (10 mL) was added *tert*-butyl 3,8-diazabicyclo[3.2.1]octane-8-carboxylate (212.3 mg, 1.000 mmol) and *N*,*N*-diisopropylethylamine (0.50 mL, 3.0 mmol, 3 equiv). The resulting reddish black solution was stirred at ambient temperature for 5 min when LC-MS indicated the full conversion of the starting material. Silica gel was added directly to the reaction solution. The suspension was concentrated and loaded onto a cartridge for flash column chromatography (0–100% EA-hexanes, 12-g RediSep(R) Rf column, Teledyne ISCO, Lincoln, NE) to give the title compound (278.1 mg, 0.6493 mmol, 65% yield for two steps) as a white crystalline solid. ^1^H NMR (400 MHz, CDCl_3_) δ 8.84 (d, *J* = 0.6 Hz, 1H), 4.70 – 4.22 (m, 4H), 3.72 (s, 2H), 1.98 (dd, *J* = 8.7, 4.3 Hz, 2H), 1.67 (d, *J* = 7.7 Hz, 2H), 1.51 (s, 9H). Accurate MS (ESI-TOF) calculated for C_18_H_21_Cl_2_FN_5_O_2_ [M + H]^+^ 428.1056, found 428.1121.

***tert*-Butyl (1R,5S)-3-(7-chloro-8-fluoro-2-((tetrahydro-1H-pyrrolizin-7a(5H)-yl)methoxy)pyrido[4,3-d]pyrimidin-4-yl)-3,8-diazabicyclo[3.2.1]octane-8-carboxylate (S37).** A 20-mL vial equipped with a stir bar was charged with **S35** (364.0 mg, 0.8499 mmol). Affixed with a screw cap with a PTFE septum, the vial was deaerated and backfilled with argon three times. 1,2,3,5,6,7-Hexahydropyrrolizin-8-ylmethanol (**S36**, 240.1 mg, 1.700 mmol, 2 equiv) in 1,4-dioxane (4.3 mL) and *N*,*N*-diisopropylethylamine (0.44 mL, 2.6 mmol, 3 equiv) were added sequentially. The mixture was stirred at 80 °C for 6 h when LC-MS indicated the conversion plateaued. The reaction mixture was first concentrated, then partitioned between water and EA. The aqueous phase was extracted with EA. The combined organic solution was washed with brine, dried over anhydrous sodium sulfate and concentrated. The crude was purified by flash column chromatography (0–20% MeOH*-DCM, *MeOH contains 2% ammonia, 12-g RediSep(R) Rf column, Teledyne ISCO, Lincoln, NE) to give the title compound (325.1 mg, 0.6099 mmol, 72% yield) as a white foam. ^1^H NMR (400 MHz, CDCl_3_) δ 8.74 (d, *J* = 0.5 Hz, 1H), 5.30 (s, 1H), 4.80 (s, 2H), 4.40 (s, 3H), 3.92 (dq, *J* = 12.9, 6.6 Hz, 2H), 3.81 (d, *J* = 7.7 Hz, 1H), 3.70 (dq, *J* = 12.8, 6.7 Hz, 2H), 3.04 – 2.88 (m, 3H), 2.47 – 1.91 (m, 15H), 1.86 – 1.66 (m, 3H), 1.63 (s, 9H). Accurate MS (ESI-TOF) calculated for C_26_H_35_ClFN_6_O_3_ [M + H]^+^ 533.2443, found 533.2467.

***tert*-Butyl (1R,5S)-3-(8-fluoro-7-(3-(methoxymethoxy)-8-((triisopropylsilyl)ethynyl)naphthalen-1-yl)-2-((tetrahydro-1H-pyrrolizin-7a(5H)-yl)methoxy)pyrido[4,3-d]pyrimidin-4-yl)-3,8-diazabicyclo[3.2.1]octane-8-carboxylate (S38).** A 20-mL scintillation vial equipped with a stir bar was charged with **S37** (53.3 mg, 0.100 mmol), **S32** (64.3 mg, 0.130 mmol, 1.3 equiv), [1,1’-bis(diphenylphosphino)ferrocene]palladium(II) dichloride (14.6 mg, 0.0200 mmol, 0.2 equiv), and cesium carbonate (97.8 mg, 0.300 mmol, 3 equiv). The vial capped with a screw cap with a silicone septum was deaerated and backfilled with argon three times. 1,4-Dioxane (3 mL) and degassed water (1 mL) were added. The resulting mixture was stirred at 100 °C for 2 h. LC-MS indicated the 98% conversion of the starting material. The reaction crude was purified by prep-HPLC (5–95% MeCN-H_2_O gradient with 0.1% formic acid, 40 min method) to give the title compound in its 1:1 formic acid salt form (15.5 mg, 0.0179 mmol, 18% yield) as a white fluffy solid. ^1^H NMR (400 MHz, DMSO) δ 9.15 (s, 1H), 8.01 (dd, *J* = 8.4, 1.4 Hz, 1H), 7.69 (d, *J* = 2.6 Hz, 1H), 7.61 (dd, *J* = 7.2, 1.4 Hz, 1H), 7.53 (dd, *J* = 8.2, 7.2 Hz, 1H), 7.28 (d, *J* = 2.6 Hz, 1H), 5.37 (s, 2H), 4.74 (d, *J* = 12.8 Hz, 1H), 4.38 – 4.15 (m, 3H), 4.00 (s, 2H), 3.74 (d, *J* = 12.4 Hz, 1H), 3.43 (s, 3H), 2.93 (dt, *J* = 10.1, 5.4 Hz, 2H), 2.58 – 2.53 (m, 1H), 1.95 – 1.52 (m, 12H), 1.46 (s, 8H), 0.81 (dd, *J* = 7.5, 6.3 Hz, 16H), 0.47 (p, *J* = 7.5 Hz, 3H). Accurate MS (ESI-TOF) calculated for C_49_H_66_FN_6_O_5_Si [M + H]^+^ 865.4848, found 865.4904.

***tert*-Butyl (1R,5S)-3-(7-(8-ethynyl-3-(methoxymethoxy)naphthalen-1-yl)-8-fluoro-2-((tetrahydro-1H-pyrrolizin-7a(5H)-yl)methoxy)pyrido[4,3-d]pyrimidin-4-yl)-3,8-diazabicyclo[3.2.1]octane-8-carboxylate (S39).** A 4-mL vial equipped with a stir bar was charged with **S38** (15.5 mg, 0.0179 mmol) and cesium fluoride (27.2 mg, 0.179 mmol, 1 equiv). DMF (0.5 mL) was added and the reaction mixture was stirred at ambient temperature. Full conversion was achieved within 30 min as indicated by LC-MS. Majority of solvent was removed in vacuo. The residue was dissolved in 70% MeCN-H_2_O (4 mL) and was purified by prep-HPLC (5–95% MeCN-H_2_O gradient with 0.1% formic acid, 40 min method) to give the title compound in its 1:1 formic acid salt form (6.6 mg, 0.0087 mmol, 49% yield) as a white solid. ^1^H NMR (400 MHz, CDCl_3_) δ 9.00 (s, 1H), 7.83 (dd, *J* = 8.3, 1.3 Hz, 1H), 7.60 (dd, *J* = 7.2, 1.3 Hz, 1H), 7.53 (d, *J* = 2.6 Hz, 1H), 7.39 (dd, *J* = 8.2, 7.2 Hz, 1H), 7.33 (d, *J* = 2.6 Hz, 1H), 5.37 – 5.28 (m, 2H), 4.86 – 4.33 (m, 7H), 3.97 – 3.58 (m, 4H), 3.52 (s, 3H), 3.03 – 2.82 (m, 2H), 2.67 (s, 1H), 2.46 – 2.32 (m, 2H), 2.29 – 2.16 (m, 2H), 2.09 (dt, *J* = 15.0, 7.5 Hz, 2H), 2.03 – 1.89 (m, 5H), 1.76 – 1.55 (m, 11H), 1.52 (s, 9H). Accurate MS calculated for C_40_H_46_FN_6_O_5_ [M + H]^+^ 709.3514, found 709.3590.

**4-(4-((1R,5S)-3,8-Diazabicyclo[3.2.1]octan-3-yl)-8-fluoro-2-((tetrahydro-1H-pyrrolizin-7a(5H)-yl)methoxy)pyrido[4,3-d]pyrimidin-7-yl)-5-ethynylnaphthalen-2-ol (EX185).** A 4-mL vial equipped with a stir bar was charged with **S39** (2.2 mg, 0.0029 mmol). The vial affixed with a screw cap and PTFE septum was deaerated and backfilled with argon three times. 1,4-Dioxane (0.5 mL) was added and the resulting solution was cooled to 0 °C. Into the reaction vial, concentrated HCl solution (0.5 mL) was added and the mixture was stirred at 0 °C for 1 h when LC-MS indicated the full conversion of the starting material. The reaction mixture was diluted in 5 mL 50% MeCN-H_2_O, filtered, and directly injected into a prep-HPLC for purification (5–95% MeCN-H_2_O gradient with 0.1% formic acid, 40 min method). The fraction collected with target MS was flash frozen in liquid nitrogen and lyophilized to give the title compound in its 1:2 formic acid salt form (1.6 mg, 0.0024 mmol, 84% yield) as a fluffy white solid. ^1^H NMR (400 MHz, MeOD) δ 9.10 (d, *J* = 4.9 Hz, 1H), 8.29 (s, 1H), 7.85 (dd, *J* = 8.4, 1.3 Hz, 1H), 7.53 (dd, *J* = 7.2, 1.3 Hz, 1H), 7.42 (dd, *J* = 8.3, 7.1 Hz, 1H), 7.36 (dd, *J* = 2.6, 1.4 Hz, 1H), 7.18 (d, *J* = 2.6 Hz, 1H), 4.81 (s, 2H), 4.69 (t, *J* = 4.3 Hz, 2H), 4.63 – 4.39 (m, 2H), 4.12 (s, 2H), 3.93 (t, *J* = 14.5 Hz, 2H), 3.71 (dt, *J* = 12.3, 6.9 Hz, 2H), 3.05 (s, 1H), 2.41 – 2.06 (m, 12H). Accurate MS (ESI-TOF) calculated for C_33_H_34_FN_6_O_2_ [M + H]^+^ 565.2727, found 565.2740.

**QUANTIFICATION AND STATISTICAL ANALYSIS**

Data from multiple independent experiments (N) are presented as mean values +/- standard error of the mean (SE) and data involving technical replicates are presented as mean +/- standard deviation (SD) as indicated in the figure legends. The number of experimental or technical replicates for each experiment is also described in each individual figure legend. Apparent affinity values were determined using the sigmoidal dose-response (variable slope) equation available in GraphPad Prism (Version 8).
